# The sodium channel SCN2A regulates cortical excitatory and inhibitory neurogenesis

**DOI:** 10.1101/2025.01.28.635170

**Authors:** Jarryll A. Uy, Zahra Dargaei, Sarah Geahchan, Laura Botler, Yi Pan, Chad O. Brown, Jennifer L. Howe, Stephen W. Scherer, Jaideep S. Bains, Karun K. Singh

## Abstract

Voltage-gated sodium channels regulate neuronal excitability and synaptic transmission in the postnatal and adult brain. The gene *SCN2A*, encoding the sodium channel Na_v_1.2, regulates synaptic development and variants in *SCN2A* are associated with autism spectrum disorders (ASD) and a broad spectrum of epilepsy phenotypes, including early-onset developmental and epileptic encephalopathies. The expression pattern of *SCN2A* begins during prenatal cortical development, prior to the onset of synaptic transmission, but it is unknown whether SCN2A regulates early cortical development through mechanisms independent of synaptic transmission. Here we reveal that isogenic and ASD patient-derived human forebrain organoids modelling a loss of SCN2A function display impaired excitatory and inhibitory neurogenesis, leading to a developmental imbalance. Unexpectedly, we find precocious generation of cortical inhibitory neurons is driven by elevated Sonic hedgehog signaling and is reversible through pharmacological inhibition. Functionally, these developmental phenotypes arise due to Na_v_1.2-dependent sodium channel dysfunction and reduced action potential generation, leading to abnormal neuronal network activity. Our results identify a mechanism for cortical excitatory and inhibitory neurogenesis involving *SCN2A*, and reveal that early neurogenesis deficits precede postnatal neural circuit dysfunction in SCN2A-associated disorders.

## Introduction

Voltage-gated sodium channels (VGSCs) are critical regulators in the generation and propagation of action potentials, the electrophysiological unit of activity in neurons^1^. Mutations that alter VGSC function can result in channelopathies such as neurodevelopmental disorders. *SCN2A*, encoding Na_v_1.2, is one of the primary sodium channels expressed in the central nervous system that regulates action potentials in developing glutamatergic neurons of the brain^2^. Variants in *SCN2A* lead to three primary disorders: 1) autism spectrum disorder (ASD)/intellectual disability (ID) with some individuals developing seizures and 2) benign familial neonatal-infantile seizures and 3) developmental epileptic encephalopathies such as Ohtahara syndrome or West syndrome^3–5^. Large scale sequencing studies have identified *SCN2A* as a recurrent top ASD risk gene with variants typically associated with loss-of-function (LOF)^6–10^, highlighting its importance in neurodevelopment.

The canonical role of *SCN2A* has been primarily studied in postnatal and adult mouse neurons. *SCN2A* is expressed in the neocortex in glutamatergic neurons and GABAergic neurons^11–13^ in the axon initial segment, the site of action potential initiation, where it regulates neuronal excitability^2,14–18^. ASD-associated variants in SCN2A have been found to impair channel function and result in reduced neuronal excitability^19^. Mouse models of heterozygous loss of *Scn2a* display haploin sufficient phenotypes of reduced axonal and dendritic excitability, impaired synaptic transmission and absence-like seizures^2,14,15,20,21^. Scn2a has also been shown to regulate action potential backpropagation in neuronal dendrites, providing instructive cues that may influence synaptic plasticity^15,21^. Similar findings have been reported in human pluripotent stem cell (PSC)-derived neurons from either *SCN2A* patient cells or CRISPR-engineered cells^22–25^. While these studies support the neurophysiological role of *SCN2A* in postnatal neurons, it remains unknown whether *SCN2A* has non-synaptic roles during neurodevelopment.

*SCN2A* is among the top ranked ASD risk genes to date^9,10^ and is developmentally regulated in the brain^26^. Interestingly, human brain atlas studies reveal that SCN2A expression begins in neurons during prenatal time points, prior to the postnatal onset of fast synaptic transmission^27–29^. Mutations in other sodium channels such as *SCN3A* (Na_v_1.3) have been reported to alter channel physiology and disrupt early processes such as neuronal migration^30,31^. Further, screening of ASD risk genes in *Xenopus tropicalis* identified that loss of *scn2a* caused deficits in neuron production^32^. Given the early expression pattern of SCN2A and previous studies on related sodium channels, it is unknown whether SCN2A regulates human prenatal cortical development independently of its canonical role in neuronal excitability and synaptogenesis.

To study SCN2A during early cortical development, we used human forebrain organoid models that recapitulate early cortical development. We generated ASD patient and engineered human pluripotent stem cell lines that are homozygous (*SCN2A^-/-^)* or heterozygous for *SCN2A* loss-of-function alleles (*SCN2A^+/G^*^1744^*^X^* and *SCN2A^+/R^*^607^*^X^*)^23^ (**Fig. 1a**). We performed single-cell RNA sequencing (scRNAseq), and uncovered profound transcriptional vulnerabilities in excitatory neurons involving neurogenesis, leading to deficits in excitatory neuron production. Unexpectedly, *SCN2A^+/G^*^1744^*^X^*and *SCN2A^-/-^*organoids also revealed precocious production, abnormal morphology and function of inhibitory neurons (INs). Abnormal inhibitory neuron production occurrs due to elevated Sonic hedgehog (SHH) signaling and is reversible with pharmacological inhibition of SHH signaling. Electrophysiology experiments revealed dysfunctional sodium channel activity and action potential generation in *SCN2A* mutant organoids, which led to subsequent abnormal neuronal network activity via calcium activity. Overall, our work delineates a mechanism for cortical neurogenesis whereby early neural activity, driven by SCN2A, regulates the genesis of both cortical excitatory and inhibitory neurons. These data indicate that SCN2A shapes excitatory-inhibitory balance through a neurogenic mechanism, prior to its postnatal role in regulating neuronal excitability and synaptogenesis.

**Fig. 1.**
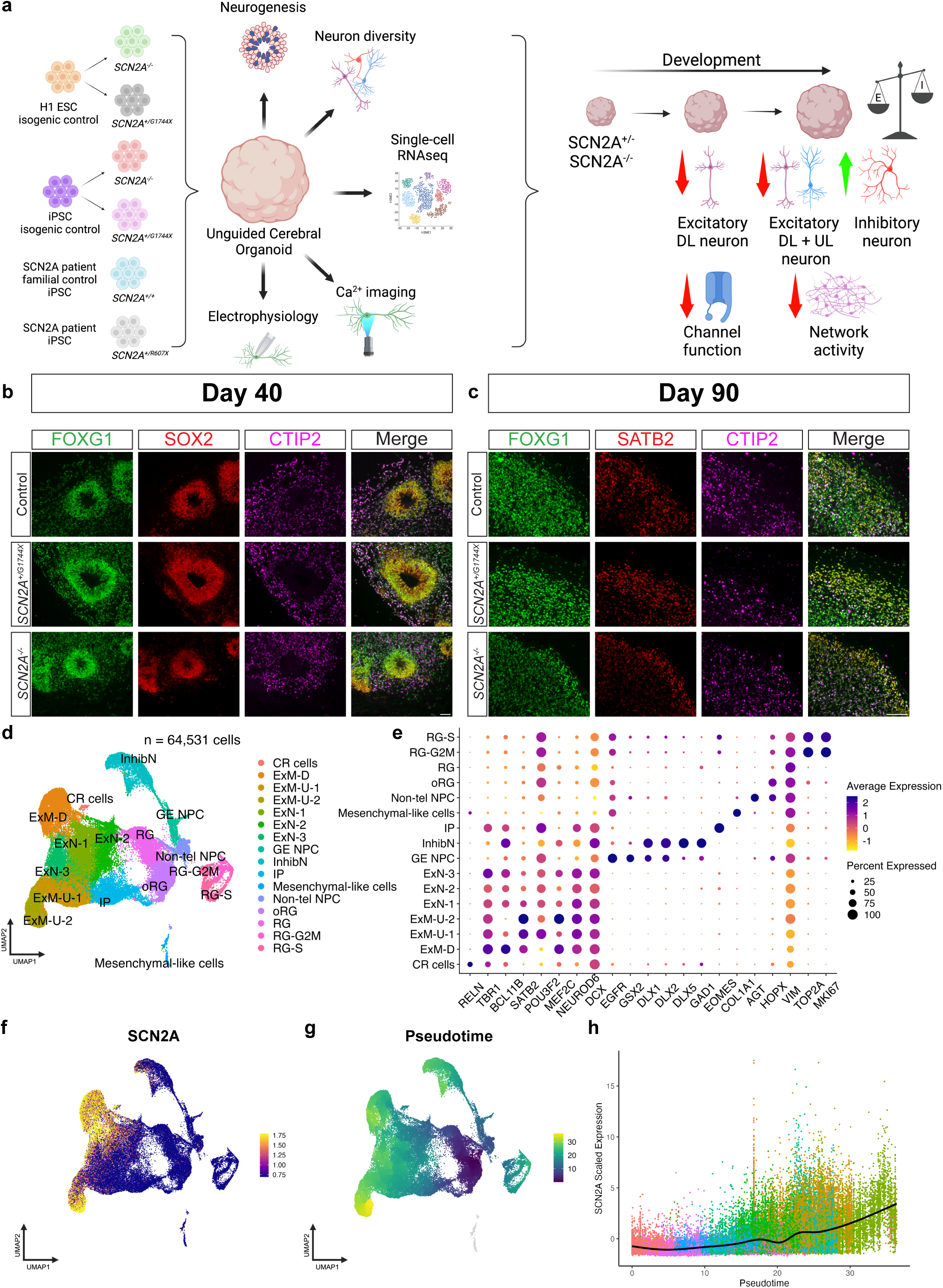
Generation and characterization of hESC-derived isogenic *SCN2A* mutant cerebral organoids. **a)** Schematic describing generation of cerebral organoids from *SCN2A* patient or CRISPR engineered hPSCs and experimental overview. A combination of immunohistochemistry, single-cell RNAseq, functional imaging and electrophysiology uncover a neurogenic role for *SCN2A* in human prenatal neurodevelopment through regulating the excitatory/inhibitory balance. **b)** Expression of telencephalic marker FOXG1, neural progenitor marker SOX2 and excitatory deep layer neuron marker CTIP2 in Day 40 cerebral organoids. N=3 independent batches with at least 3 organoids per genotype. Scale bar = 50 μm. **c)** Expression of telencephalic marker FOXG1, excitatory upper layer neuron marker SATB2 and excitatory deep layer neuron marker CTIP2 in Day 90 cerebral organoids. N=3 independent batches with at least 3 organoids per genotype. Scale bar = 100 μm. **d)** Uniform manifold approximation and projection (UMAP) dimensional reduction and unbiased clustering of 64,531 cells showing distribution of annotated cell types from Day 90 cerebral organoids. (N=3 independent batches) Abbreviations: CR cells, Cajal-Retzius cells; ExM-D, maturing excitatory neurons - deep layer; ExM-U, maturing excitatory neurons – upper layer; ExN, newborn excitatory neurons; GE NPC, ganglionic eminence-like neural progenitor cells; InhibN, inhibitory neuron; IP, intermediate progenitors; Mesenchymal-like cells; Non-tel NPC, non-telencephalic neural progenitor cells; RG, radial glia; RG-G2M, G2M-phase cycling radial glia; RG-S, S-phase cycling radial glia. **e)** Dot plot showing expression of selected well-known marker genes used for cell-type annotation. **f)** Feature plot showing *SCN2A* expression is enriched in the excitatory and inhibitory neuron clusters. **g)** UMAP plot showing distribution of pseudotime inferred using Monocle3 with radial glia as the starting node. **h)** Scaled expression of *SCN2A* across pseudotime in the integrated scRNAseq dataset. Trend-line shows the expression of *SCN2A* as a function of pseudotime. Higher pseudotime values indicate greater maturity as indicated by multiple neuronal classes. Created in BioRender. Singh, K. (2026) https://BioRender.com/0otxclw.

## Results

### *SCN2A* is expressed in glutamatergic and GABAergic neurons throughout the developing human cortex

To understand the role of SCN2A in early (prenatal) neurodevelopment, we examined expression patterns in cortical tissues from previously published transcriptome data. Analysis of the developmental transcriptome data from the BrainSpan Atlas of the Developing Human Brain^28^ showed *SCN2A* expression begins as early as postconceptional week (PCW) 16 of the developing human cortex and peaks at around birth (**Fig. S1a, b**). Consistent with this pattern, scRNAseq data of mid-gestation PCW 17-18 prenatal cortex^29^ showed an enrichment of *SCN2A* in maturing excitatory deep and upper layer neurons (**Fig. S1c**). Further, leveraging the scApex atlas^33^ that contains scRNAseq data of developing cerebral organoids, we found *SCN2A* to be expressed primarily in later-stage mature organoids (>2 months; **Fig. S1d**). We also examined the expression of *SCN2A* in the adult cortex using data from the Allen Brain Atlas^34^ and found *SCN2A*, along with the other CNS neuronal VGSCs, to be expressed in both glutamatergic and GABAergic lineages (**Fig. S1e**). Additionally, SCN2A protein is expressed across human brain development including prenatal time points^27^. These data are in line with a previous study demonstrating *SCN2A* is expressed in the prenatal cortex^26^. Together, these gene expression patterns across human brain development indicate *SCN2A* is expressed in the prenatal brain and suggest a potential role during early neurodevelopment.

### Generation and characterization of hESC-derived isogenic *SCN2A* mutant cerebral organoids

To investigate the human neurodevelopmental impact of SCN2A, we used CRISPR-Cas9 on the well-characterized human H1 embryonic stem cell (ESC) line to generate an isogenic homozygous knockout of SCN2A (*SCN2A^-/-^*) and a heterozygous knock-in of a late-truncating ASD patient mutation at position G1744 (*SCN2A^+/G^*^1744^*^X^)* with known late-onset seizures (**Fig. S2a**). We adopted an isogenic experimental design on the basis of eliminating confounding effects of genetic background as well as improving large-scale transcriptomics precision^35^. Each H1 cell line edit was validated through sanger sequencing or droplet digital PCR (ddPCR) and were karyotypically normal (**Fig. S2b-g**). Further, SCN2A protein levels were analyzed in cerebral organoids using mass spectrometry, validating heterozygous expression for *SCN2A^+/G^*^1744^*^X^* and a lack of protein in *SCN2A^-/-^*organoids (**Fig. S2h**).

We generated cerebral organoids as previously described for their cortical forebrain identity, cellular diversity and reproducibility^33,36–38^. Early cerebral organoids (Day 40) developed as expected by expressing the telencephalic marker FOXG1, SOX2 for neural progenitors in ventricular-like progenitor zones, and CTIP2 for deep layer excitatory neurons (**Fig. 1b**). In mature organoids (Day 90), most progenitor zones dissipated to produce cortical neurons that were upper-layer SATB2^+^ and deep-layer CTIP2^+^ supporting typical development of these organoids (**Fig. 1c**).

Having established a reliable organoid protocol and isogenic model, we matured H1 ESC-derived cerebral organoids to Day 90, where there is more cell diversity and maturation^36,39^ and performed scRNAseq. We used 3 independent differentiations (N=3 batches) composed of pooled samples that contained 3 organoids per genotype to control for inter-organoid variability within a batch (3 organoids of the same genotype pooled as 1 sample). For simplicity, the isogenic H1 control is referred to as “Control”, *SCN2A^+/G^*^1744^*^X^* as “SCN2A G1744X”, and *SCN2A^-/-^*as “SCN2A KO” for our scRNAseq experiment. After quality control filtering, we recovered 64,531 cells from 27 cerebral organoids for analysis (**Fig. 1d**). We performed principal component analysis, integration and UMAP clustering resulting in 16 distinct clusters that were annotated using a combined approach based on previously published guidelines^40^. We first used scClassify, a semi-automated annotation method^41^, with two reference datasets (2 month-old cerebral organoids^33^ and mid-gestation prenatal cortex^29^; **Fig. S3a-d**). We used a panel of well-known cell type markers^29,33,42–44^ (**Fig. 1e**) as verification of the cell type predictions. To confirm cellular and regional identity of our organoids, we performed unbiased spatial mapping using VoxHunt^45^ onto the Allen Brain Atlas embryonic day 15.5 mouse brain and found a high correlation to the dorsal telencephalon (**Fig. S4a, b**). When mapped to the BrainSpan atlas^28^, our cerebral organoids closely resemble neocortical areas of the prenatal brain (**Fig. S4c, d**). We further corroborated forebrain identity by examining the expression of known markers of the midbrain (PAX5 and GBX2)^46,47^ and hindbrain (HOXB2 and ISL1)^48,49^ which showed very low expression (**Fig. S4e**). After annotation, we examined the expression of *SCN2A* and found it to be expressed in the excitatory neuron populations as well as INs (**Fig. 1f**). In addition, we used trajectory inference analysis (monocle3^50^) using radial glia (RG) as the root node to quantify the expression of *SCN2A* across organoid development (**Fig. 1g**). As expected, we found *SCN2A* expression to increase across development (higher pseudotime values) in the newborn excitatory neurons (ExN) and deep layer and upper layer maturing excitatory neurons (ExM-D, ExM-U; **Fig. 1h**), with little to no expression in progenitor cells (lower pseudotime values). Thus, our organoid model is able to recapitulate the major prenatal brain cell types and expression of *SCN2A* during development.

### Single-cell RNA sequencing reveals impaired developmental trajectories of excitatory neurons caused by loss of *SCN2A*

To uncover cellular and molecular impairments caused by the loss of *SCN2A*, we performed trajectory inference and differentially expressed gene (DEG) analysis of H1 isogenic cerebral organoids. By plotting each condition along a pseudotime density plot, we found a significant difference in pseudotime distribution (**Fig. 2a**). Specifically, we looked at ExM-D (deep-layer neurons) and ExM-U-2 (upper-layer neurons), the principal cell types expressing *SCN2A*, and found a significant deviation from the Control distribution (**Fig. 2b, c**). These altered pseudotime distributions suggest the developmental dynamics and maturation in *SCN2A* mutant organoids may be impaired.

**Fig. 2.**
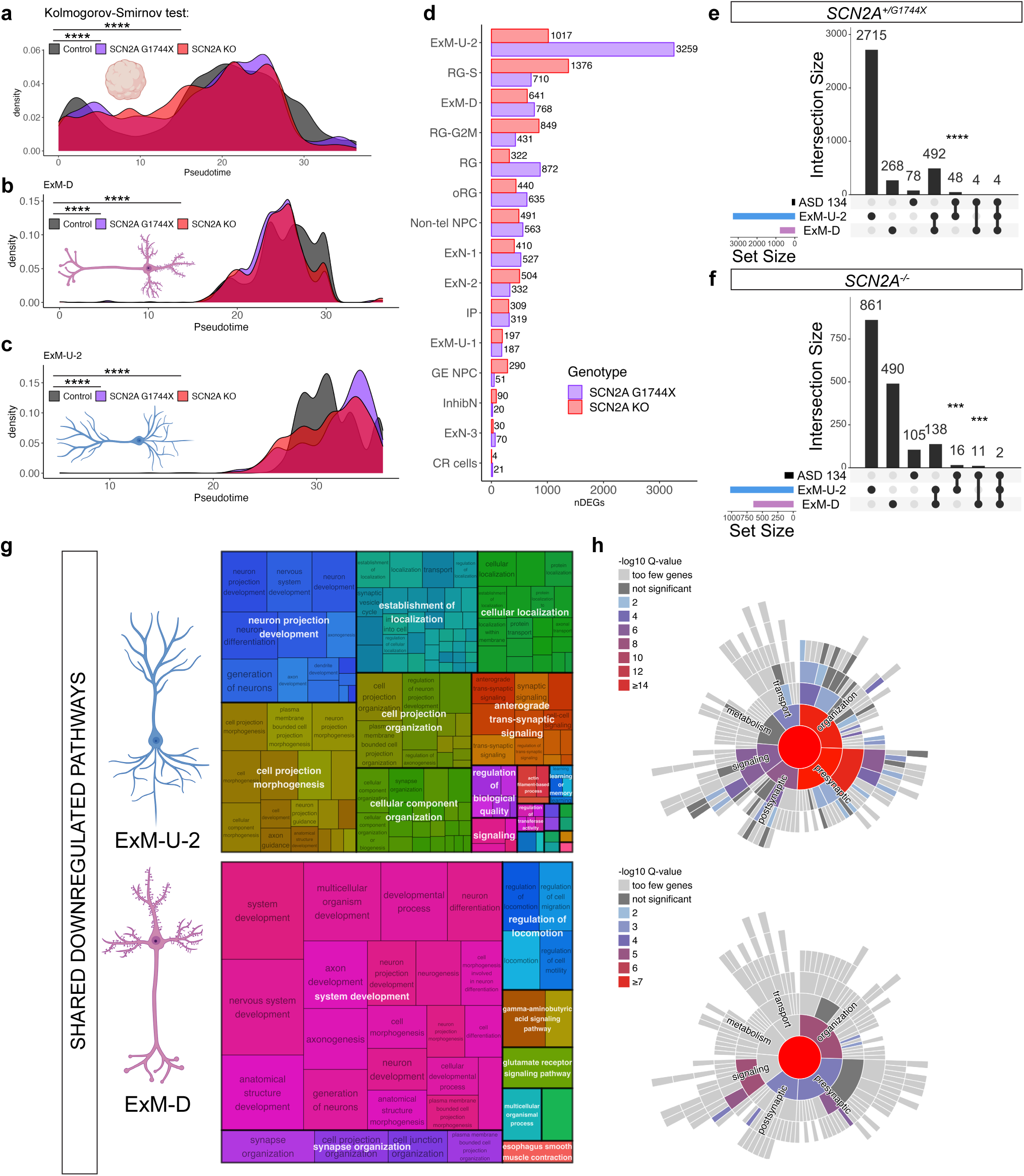
Single-cell RNAseq reveals developmental perturbations and vulnerabilities of excitatory neurons caused by loss of *SCN2A*. **a-c)** Density plot showing pseudotime distribution of all cell types (a), deep layer neurons (b) or upper layer neurons (c). (a) Control vs *SCN2A^+/G^*^1744^*^X^* p=1.69×10^-61^; Control vs *SCN2A^-/-^* p=1.98×10^-94^. (b) Control vs *SCN2A^+/G^*^1744^*^X^* p < 2.2×10^-16^; Control vs *SCN2A^-/-^* p < 2.2×10^-16^. (c) Control vs *SCN2A^+/G^*^1744^*^X^* p < 2.2×10^-16^; Control vs *SCN2A^-/-^*p=1.99×10^-7^. A two-sided Kolmogorov-Smirnov test was used for all distributions. **d)** Number of differentially expressed genes (DEGs) by cell type between each *SCN2A* mutant versus Control. DEG counts are derived from Tables S1-S2. **e, f)** Number of ASD risk genes enriched in deep and upper layer neurons of *SCN2A^+/G^*^1744^*^X^*(e) or *SCN2A^-/-^* organoids (f) using one-sided Fisher’s exact test (GeneOverlap). (e) ExM-U-2 p=1.1×10^-8^; ExM-D p=0.21 (n.s.). (f) ExM-U-2 p=5.2×10^-4^; ExM-D p=9.6×10^-4^. **g)** Pathway enrichment of shared downregulated DEGs in deep and upper layer neurons for each *SCN2A* mutant. In each treemap plot, the overarching biological theme is shown in the white text large rectangles, and its size is dependent on the number of GO biological process terms shared while black text smaller rectangles size is dependent on the significance of that term. **h)** Sunburst plots of enriched SynGO terms using shared downregulated DEGs of either deep or upper layer neurons. The color of the box shows the ratio for enrichment (q-value < 0.05). *p < 0.05, **p < 0.01, ***p < 0.001, ****p < 0.0001; n.s., not significant. Source data are provided as a Source Data file. Created in BioRender. Singh, K. (2026) https://BioRender.com/0otxclw.

To identify cell type-specific vulnerabilities, we performed DEG analysis (p-adj < 0.05) in each cell type. We found that both deep and upper layer neurons were among the highest in number of DEGs present (**Fig. 2d, Tables S1** and **S2**). We then performed a gene overlap analysis of DEGs in those cell types and ASD risk genes from Autism Speaks’ MSSNG database^9^ and found a significant overlap primarily in the upper layer neurons (**Fig. 2e, f**). We performed a GO term enrichment analysis using g:Profiler^51^ combined with semantic reduction using REVIGO^52^ to uncover impaired biological themes. We used a combined list of DEGs shared between both genotypes per cell type as we predicted their effects to be similar. Our GO analysis revealed that both upper and deep layer neurons shared downregulated pathways involving neuron development, neurogenesis, synaptic signaling and neuron projection development (**Fig. 2g** and **Tables S3-S6**). We then subjected our shared list of downregulated DEGs to SynGO analysis^53^ to identify synaptic GO terms. SynGO analysis revealed an enrichment in presynase GO terms, concordant with the role of *SCN2A* in action potential generation (**Fig. 2h**). Altogether, our DEG analysis suggests loss of *SCN2A* consistently perturbed neuron development and maturation in deep and upper layer excitatory neurons.

### Complete or partial loss of *SCN2A* causes early and late disruptions in excitatory neuron production

*SCN2A* is known to regulate neuronal properties such as excitability^14–17,21,23–25^, however, its role outside of the action potential and during early neurodevelopment remain unknown. Unexpectedly, our transcriptomics revealed neurogenesis and neuron development to be impaired (**Fig. 2g**) suggesting *SCN2A* may regulate these functions. We next examined proportion changes in our organoid model but given the dynamic and contiguous differentiation occurring in organoids, simple discrete proportion tests may be limited in statistical power. To overcome those limitations, we performed differential abundance (DA) testing using Milo^54^. DA testing revealed that loss of *SCN2A* leads to a decrease in deep and upper layer excitatory neurons (**Fig. 3a, b**). Notably, we found an increase in INs suggesting *SCN2A* may regulate the generation of both neuronal lineages.

**Fig. 3.**
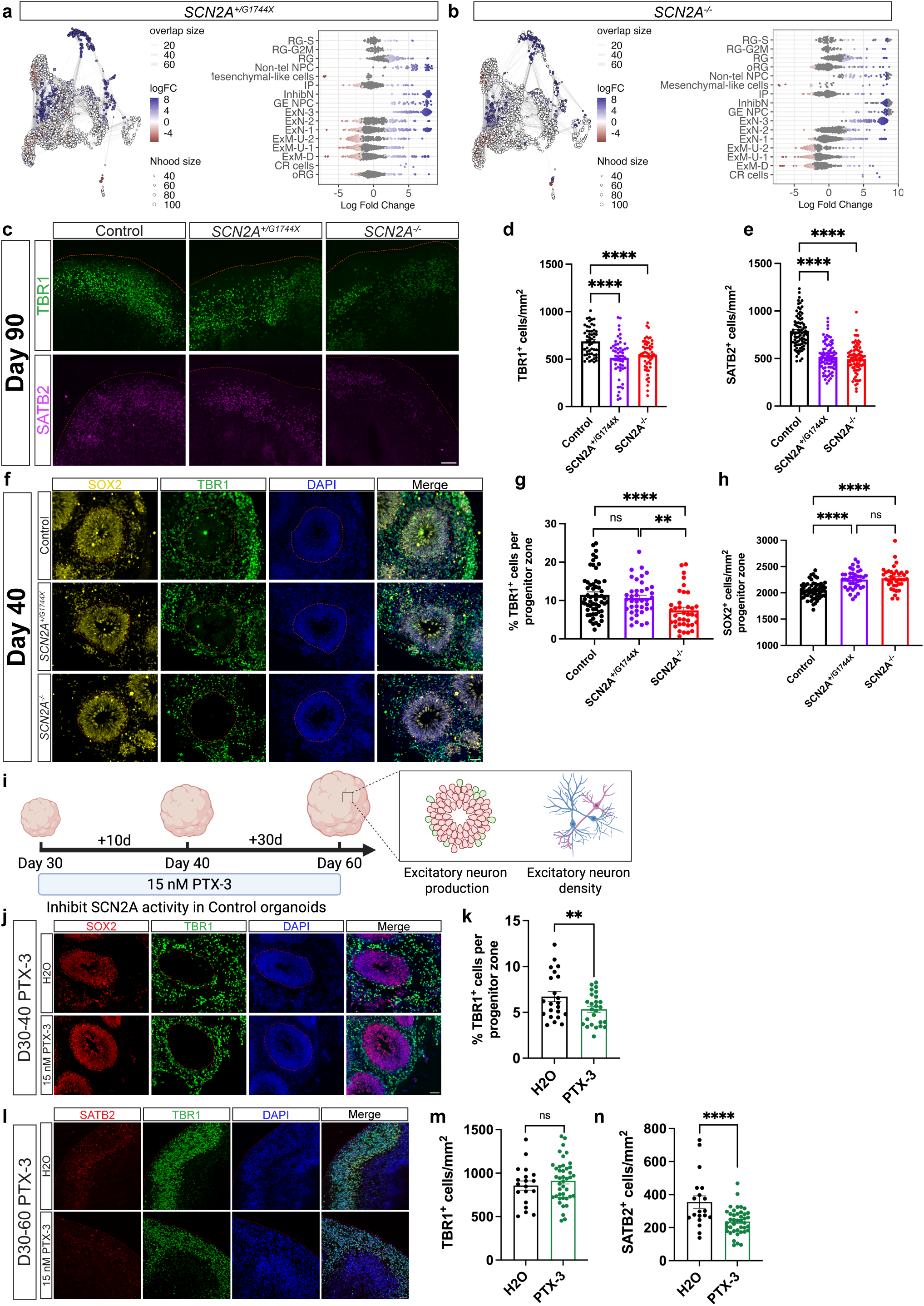
Complete or partial loss of *SCN2A* causes early and late disruptions in excitatory neuron production. **a, b)** Differential abundance of neural cell types in *SCN2A^+/G^*^1744^*^X^* (a) or *SCN2A^-/-^*(b) versus control organoids. UMAP: colored neighborhoods are differentially abundant at spatial FDR < 0.1. Beeswarm: differentially abundant cell types. **c)** Immunostaining for the deep layer neuron marker TBR1 and upper layer neuron marker SATB2. Scale bar = 100 μm. **d, e)** Density of TBR1^+^ (d) and SATB2+ (e) neurons in Day 90 organoids. n = 54/62/47 (d) and 84/83/83 (e) FOVs for Control/*SCN2A^+/G^*^1744^*^X^*/*SCN2A^-/-^*, from 18/17/12 organoids across 3 independent batches; mixed-effects model (Fixed effect: Genotype, random effect: batch) with Tukey post-hoc. (d) Genotype χ^2^(2)=31.82, p=1.2×10^-7^; Control vs *SCN2A^+/G^*^1744^*^X^* p=8.7×10^-5^; Control vs *SCN2A^-/-^* p=2.4×10^-6^; *SCN2A^+/G^*^1744^*^X^* vs *SCN2A^-/-^* p=0.46. (e) Genotype χ^2^(2)=135.4, p=4.0×10^-30^; Control vs *SCN2A^+/G^*^1744^*^X^* p=4.0×10^-14^; Control vs *SCN2A^-/-^* p=2.7×10^-14^; *SCN2A^+/G^*^1744^*^X^* vs *SCN2A^-/-^*p=0.18. **f)** Immunostaining for the deep layer neuron marker TBR1 and neural progenitor marker SOX2 in Day 40 organoids. Scale bar = 50 μm. **g)** Percentage of TBR1^+^ neurons in SOX2^+^ progenitor zones, Day 40 organoids. n = 58/39/37 progenitor zones for Con-trol/*SCN2A^+/G^*^1744^*^X^*/*SCN2A^-/-^*, from 20/18/18 organoids across 3 independent batches; mixed-effects model (Fixed effect: Genotype, random effect: batch) with Tukey post-hoc. Genotype χ^2^(2)=19.80, p=5.0×10^−5^; Control vs *SCN2A^+/G^*^1744^*^X^* p=0.84; Control vs *SCN2A^-/-^* p=5.5×10^−5^; *SCN2A^+/G^*^1744^*^X^* vs *SCN2A^-/-^* p=2.0×10^−3^. **h)** Density of SOX2^+^ cells in progenitor zones (same n and design as in g). χ^2^(2)=99.41, p=2.6×10^−2^^2^; Control vs *SCN2A^+/G^*^1744^*^X^* p=2.5×10^−1^^4^; Control vs *SCN2A^-/-^* p=1.1×10^−1^^3^; *SCN2A^+/G^*^1744^*^X^* vs *SCN2A^-/-^* p=0.47. **i)** Schematic of pharmacological inhibition of SCN2A in Control organoids using 15 nM of PTX-3. **j)** Immunostaining for TBR1 and SOX2 in Day 40 organoids. Scale bar = 50 μm. **k)** Percentage of TBR1^+^ neurons in SOX2^+^ progenitor zones of PTX-3-treated organoids. n = 22/26 progenitor zones for H2O/PTX-3, from 3/3 organoids (1 batch); mixed-effects model (Fixed effect: Condition, random effect: organoid). H2O vs PTX-3 p=1.7×10^−3^. **l)** Immunostaining for TBR1 and SATB2 in Day 60 PTX-3-treated organoids. Scale bar = 50 μm. **m, n)** Density of TBR1^+^ (m) and SATB2^+^ (n) neurons in H2O- or PTX-3-treated Day 60 organoids. n = 19/42 FOVs for H2O/PTX-3, from 2/2 organoids (1 batch); mixed-effects model (Fixed effect: Condition, random effect: organoid). (m) H2O vs PTX-3 p=0.75. (n) H2O vs PTX-3 p=2.1×10^−7^. All data are reported as mean ± SEM. Comparisons used mixed-effects models; the Genotype fixed effect was assessed by Type III Wald χ^2^ tests and pairwise comparisons by two-sided post-hoc Tukey tests; two-group comparisons (k–n) used the two-sided Wald test on the condition term. *p < 0.05, **p < 0.01, ***p < 0.001, ****p < 0.0001; n.s., not significant. Source data are provided as a Source Data file. Created in BioRender. Singh, K. (2026) https://BioRender.com/0otxclw.

Given our transcriptomics suggest a cell type imbalance and gene expression changes involving neurogenesis, we performed immunostaining validation in organoids at Day 90. We stained for TBR1, a marker of deep layer excitatory neurons and SATB2, a marker for upper layer excitatory neurons^55–57^. We observed in Day 90 *SCN2A* mutant organoids a reduction in both TBR1^+^ deep layer neurons and SATB2^+^ upper layer neurons, with no difference in cell death (**Fig. 3c-e**, **S5a, b**). Because our transcriptomics indicate neurogenesis may be impacted, we investigated early neurogenesis in organoids. At Day 40, a time of high neurogenesis^36^, we found a marked reduction in TBR1^+^ early born neurons within progenitor zones of *SCN2A^-/-^* organoids but not *SCN2A^+/G^*^1744^*^X^*(**Fig. 3f, g**). Further, there was an increase in SOX2^+^ neural progenitors with a reduction in early apoptotic cells (**Fig. 3h, S5c** and **S5d**) in agreement with our DA analysis (**Fig. 3a, b**). We confirmed these findings by repeating these experiments in two previously published and independent isogenic *SCN2A^-/-^*iPSC lines (named 19-2-2^58^ and 50B^59^; hereafter each “Control” refers to the isogenic control of the respective cell line). We validated a reduction in TBR1^+^ deep layer neurons, however, there was no change in SATB2^+^ upper layer neurons at Day 90 in the 50B background and an increase in the 19-2-2 background likely due to cell line differences (**Fig. S6a-c**). At Day 40, both isogenic iPSC lines show a reduction in TBR1^+^ early born neurons in progenitor zones (**Fig. S6d, e**). We next asked if changes in deep-layer neuron production due to loss of SCN2A function could be recapitulated by pharmacologically inhibiting sodium channels and thus, reducing neuronal activity during early development. We applied a low concentration of tetrodotoxin (TTX) in Control organoids at Day 40 to partially block sodium channels (**Fig. S7a)** and mimic reduced action potential kinetics (peak dV/dt) in neurons as previously reported in a mouse model of *Scn2a* haploinsufficiency^15^. Consistent with *SCN2A^-/-^*organoids, we observed a reduction in TBR1^+^ neurons within progenitor zones (**Fig. S7b, c**). In addition, we found no differences in neuronal (NeuN+) or progenitor (SOX2+) cell death compared to the vehicle control (**Fig. S7d-f**). We further linked these TTX-associated phenotypes to SCN2A by using a selective inhibitor PTX-3 at its reported Na_v_1.2-specific dose of 15 nM^60^ through acute and chronic treatment in Control organoids (**Fig. 3i**). We find that acute treatment of PTX-3 from Day 30-40 led to a reduction in TBR1^+^ neurons in progenitor zones (**Fig. 3j, k**). We further found that chronic treatment from Day 30-60 resulted in reductions in SATB2^+^ upper-layer neurons but not TBR1^+^ deep-layer neurons (**Fig. 3l-n**). To confirm if the decreases in excitatory neuron production were due to changes in proliferation or a shift in neuronal identity, we pulsed Control and *SCN2A* mutant organoids with BrdU for 24h and measured cell cycle exit by examining the colocalization of BrdU with Ki67^61^. Cells positive for BrdU but negative for Ki67 indicate those cells have exited the cell cycle within 24h of BrdU exposure. We found no significant changes in the percent of cells exiting the cell cycle, and proliferating progenitors, however, a slight decrease in the percentage of BrdU-labelled cells in *SCN2A^+/G^*^1744^*^X^*organoids (**Fig. S8a-d**). We further validated this finding in isogenic iPSC-derived organoids and similarly found no changes in neural progenitor proliferation or cell death (**Fig. S9a-h** and **S10a-d**). Lastly, we tested patient-derived *SCN2A^+/R^*^607^*^X^* organoids, harboring an early truncation mutation in SCN2A, compared to the familial-matched neurotypical control, and found no changes in neural progenitor proliferation (**Fig. S11a-d**). Collectively, these results suggest an *SCN2A*-dependent effect on early excitatory neurogenesis and maturation without changes in neural progenitor proliferation, and that these early changes are a consequence of reduced neuronal activity by SCN2A.

### Loss of *SCN2A* leads to precocious production of CGE-like INs

Cerebral organoids capture many of the developmental milestones and features of the telencephalon such as cell type regional diversity^62^. While cerebral organoids are abundantly composed of excitatory lineages, inhibitory lineages have been previously identified in these organoids and other neural organoid models^39,63–69^. Further, cortical inhibitory neurogenesis occurs primarily in the ventral cortex^70–72^, however, recent studies suggest a dorsal origin of a population of human cortical INs (in primary tissue and neural organoids) that resemble cells derived from the caudal ganglionic eminence (CGE)^69,73,74^. We wondered if the same observations occur in our cerebral organoid models. We performed bioinformatic characterization of our INs showing expression of the ventral lineage transcription factors DLX1/2 and DLX5 (**Fig. S12a**). We compared a panel of markers to differentiate cells from the medial ganglionic eminence (MGE) and the CGE that showed CGE-like expression patterns (**Fig. S12b**). Further, we found very low to no expression of the MGE lineage marker NKX2-1 (**Fig. S12c**). As confirmation of these CGE-like INs, our cell type annotation using mid-gestation prenatal cortex showed the highest identification for CGE INs versus MGE INs (**Fig. S3c**). Thus, our cerebral organoids are capable of producing CGE-like INs as previously described^69^.

We observed an increase in the number of INs in both *SCN2A* mutants (**Fig. 4a**), suggesting an imbalance in neuron production. To validate the increase in IN numbers, we used an AAV-mediated viral labelling method for INs with GFP (AAV-mDlx-GFP) at Day 120 given the low numbers identified in the Control condition at Day 90 (**Fig. 4b**). Viral labelling confirmed the increased number of INs in both *SCN2A* mutant organoids as well as GABAergic CGE-like identity through immunostaining (**Fig. 4c, d** and **S13a, b**). We performed morphological analyses of the INs and found *SCN2A* mutants showed increased local neurite branching (**Fig. 4e-g**) compared to the isogenic Control. We further confirmed the increase in IN population by staining for the ventral lineage marker DLX5 (**Fig. 4h, i**). Similarly, there was an increase in DLX5^+^ neurons when Control organoids are treated with PTX-3 (from the same batch as in Fig. 3) to inhibit SCN2A (**Fig. 4j, k**). We then performed the same viral labelling experiment in organoids derived from the isogenic iPSC line 19-2-2 to account for cell line variability. We found a similar increase in INs; however, not to the same degree as the hESC-derived organoids (**Fig. S14a-c**). Additionally, we labelled isogenic 50B iPSC-derived organoids and patient-derived *SCN2A^+/R^*^607^*^X^* organoids at Day 90 and found an increase in GFP-labelled INs (**Fig. S14d-g**). To confirm whether the loss of *SCN2A* alters inhibitory neurogenesis across organoid models, we generated dorsal forebrain organoids based on previous guided protocols^64,75,76^ and performed viral labelling of INs (**Fig. S15a**). Again, we found that in an independent dorsal organoid model, both *SCN2A* mutants displayed increased numbers of INs as well as DLX5^+^ INs (**Fig. S15b-e**). Lastly, we infected D130 organoids with AAV-DLX-GCaMP6f to target INs for calcium imaging to understand their functional properties (**Fig. 4l**). We found that *SCN2A^-/-^* INs have reduced amplitude but *SCN2A^+/G^*^1744^*^X^*INs showed no difference (**Fig. 4m, n**). Further, when comparing calcium event frequency distributions, both *SCN2A^-/-^*and *SCN2A^+/G^*^1744^*^X^*INs show a left-shifted distribution (**Fig. 4o**) but only *SCN2A^-/-^* show a difference in mean frequency compared to the isogenic control (**Fig. 4p**). Altogether, these data indicate that loss of SCN2A leads to precocious production of dysfunctional CGE-like INs in cerebral and dorsal forebrain organoid models, revealing a previously unknown role for *SCN2A* in neurogenesis.

**Fig. 4.**
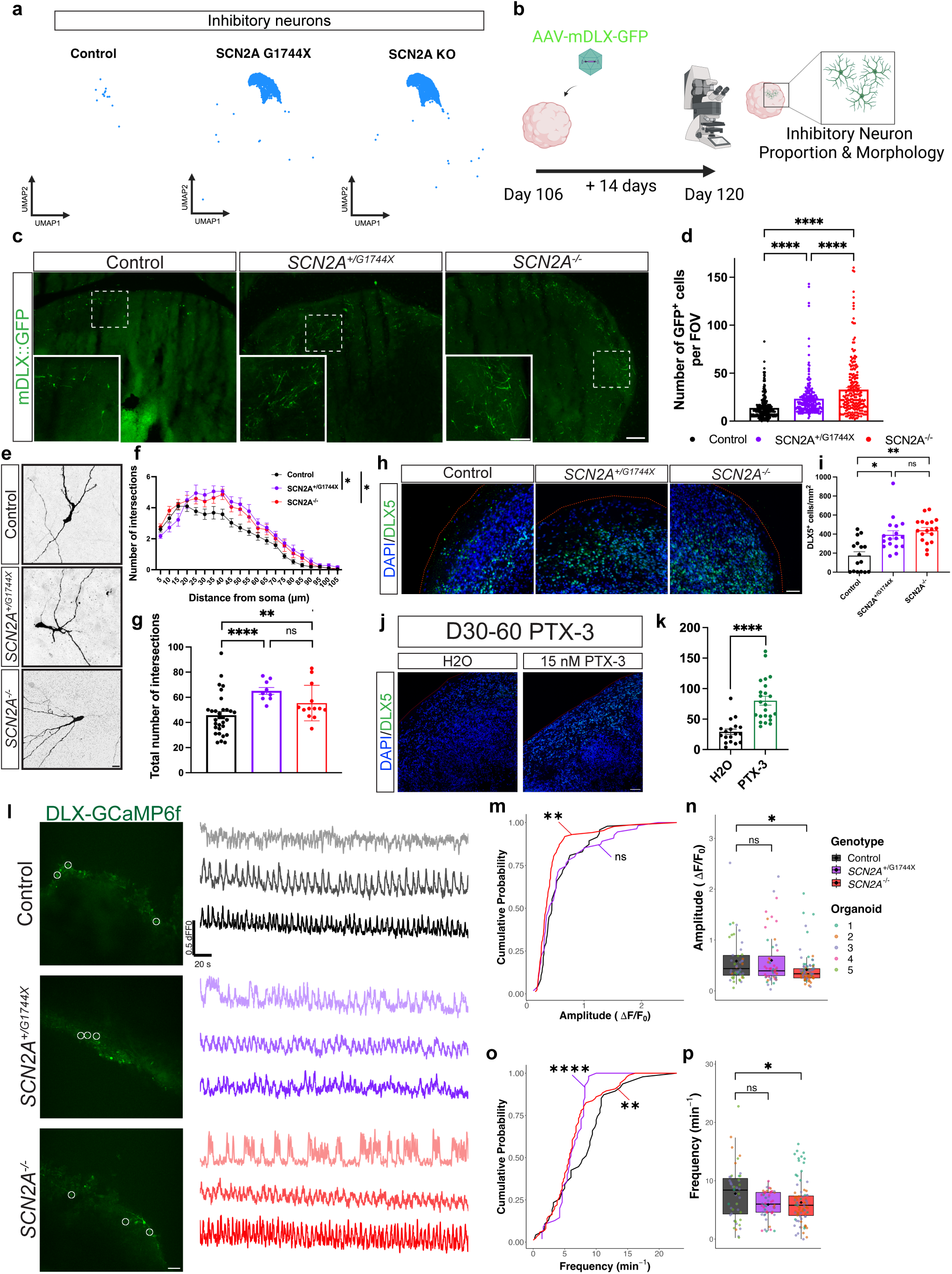
Loss of *SCN2A* leads to precocious production of CGE-like inhibitory neurons in cerebral organoids. **a)** UMAP plot showing inhibitory neurons (InhibN) by genotype. **b)** Schematic of inhibitory neuron viral labelling with AAV-GFP in Day 120 organoids. **c)** Immunostaining for AAV-mDLX-GFP-labelled inhibitory neurons. Scale bar for low magnification=200 μm and inset scale bar = 100 μm. **d)** Number of GFP^+^ cells per FOV, Day 120 organoids. n = 358/251/220 FOVs for Control/*SCN2A^+/G^*^1744^*^X^*/*SCN2A^-/-^*, from 12 organoids each across 3 independent batches; mixed-effects model (Fixed effect: Genotype, random effect: batch) with Tukey post-hoc. Genotype χ^2^(2)=7.58×10^5^, p<2.2×10^−1^^6^; Control vs *SCN2A^+/G^*^1744^*^X^*, Control vs *SCN2A^-/-^*, and *SCN2A^+/G^*^1744^*^X^* vs *SCN2A^-/-^*all p<2.2×10^−1^^6^. **e)** Immunostaining for AAV-mDLX-GFP-labelled inhibitory neurons used for Sholl analysis. Scale bar = 10 μm. **f)** Sholl analysis of inhibitory neuron morphology. n = 31/18/22 neurons for Control/*SCN2A^+/G^*^1744^*^X^*/*SCN2A^-/-^*, from 12 organoids each across 3 independent batches; mixed-effects model with a Type III Wald chi-square test and post-hoc Tukey correction (Fixed effect: Genotype and Distance, random effect: batch). Genotype F(2,68)=6.02, p=3.9×10^−3^; Distance×Gen-otype F(40,1360)=2.36, p<1×10^−4^. **g)** Total number of intersections. n = 31/18/13 neurons for Control/*SCN2A^+/G^*^1744^*^X^*/*SCN2A^-/-^*, from 12 organoids each across 3 independent batches; mixed-effects model (Fixed effect: Genotype, random effect: batch) with Tukey post-hoc. χ^2^(2)=23.03, p=1.0×10^−5^; Control vs *SCN2A^+/G^*^1744^*^X^* p=7.7×10^−5^; Control vs *SCN2A^-/-^* p=1.1×10^−3^; *SCN2A^+/G^*^1744^*^X^* vs *SCN2A^-/-^* p=1.0. **h)** Immunostaining for the inhibitory neuron marker DLX5 in Day 90 organoids. Scale bar = 100 μm. **i)** Density of DLX5^+^ neurons in Day 90 organoids. n = 16/17/18 FOVs for Control/*SCN2A^+/G^*^1744^*^X^*/*SCN2A^-/-^*, from 4 organoids each (1 batch); mixed-effects model (Fixed effect: Genotype with nested FOVs) with Tukey post-hoc. χ^2^(2)=10.79, p=4.6×10^−3^; Control vs *SCN2A^+/G^*^1744^*^X^* p=0.023; Control vs *SCN2A^-/-^* p=6.5×10^−3^; *SCN2A^+/G^*^1744^*^X^* vs *SCN2A^-/-^* p=0.92. **j)** Immunostaining for the inhibitory neuron marker DLX5 in Day 60 organoids. Scale bar = 50 μm. **k)** Density of DLX5^+^ neurons in PTX-3-treated Day 60 organoids. n = 18/23 FOVs for H2O/PTX-3, from 2/2 organoids (1 batch); mixed-effects model (Fixed effect: Genotype with nested FOVs). H2O vs PTX-3 p=4.9×10^−1^^1^. **l)** Representative fluorescent images of DLX-GCaMP6f-infected organoids and normalized GCaMP6f intensity traces recorded over 5 minutes in Day 130 organoids. White circles indicate neurons recorded in representative traces. **m, n)** Cumulative distribution and amplitude of inhibitory neuron spontaneous calcium transients, Day 130 organoids. n = 47/55/85 neurons for Control/*SCN2A^+/G^*^1744^*^X^*/*SCN2A^-/-^*, from 4/4/3 organoids (1 batch). KS (cumulative): Control vs *SCN2A^-/-^* D=0.294, p=8.0×10^−3^; Control vs *SCN2A^+/G^*^1744^*^X^* D=0.142, p=0.62. Quantification (mixed-effects model (Fixed effect: Genotype, random effect: organoid) with Tukey post-hoc): Genotype χ^2^(2)=8.22, p=0.016; Control vs *SCN2A^+/G^*^1744^*^X^*p=0.35; Control vs *SCN2A^-/-^* p=0.013; *SCN2A^+/G^*^1744^*^X^* vs *SCN2A^-/-^* p=0.32. Boxplots are median, quartiles and 1.5 IQR whiskers, black diamond represents mean of the data. **o, p)** Cumulative distribution and frequency of inhibitory-neuron spontaneous calcium transients (same n and design as in m, n). KS (cumulative): Control vs *SCN2A^-/-^* D=0.324, p=1.6×10^−3^; Control vs *SCN2A^+/G^*^1744^*^X^* D=0.438, p=3.97×10^−5^. Quantification (mixed-effects model with Tukey post-hoc): Genotype χ^2^(2)=5.81, p=0.055; Control vs *SCN2A^+/G^*^1744^*^X^* p=0.23; Control vs *SCN2A^-/-^* p=0.042; *SCN2A^+/G^*^1744^*^X^* vs *SCN2A^-/-^*p=0.67. All data are reported as mean ± SEM except in (m-p). Mixed-effects models with fixed effects assessed by Type III Wald χ^2^ tests and pairwise comparisons by two-sided post-hoc Tukey tests unless otherwise stated; cumulative distributions compared with two-sided Kolmogorov–Smirnov tests. *p < 0.05, **p < 0.01, ***p < 0.001, ****p < 0.0001; n.s., not significant. Source data are provided as a Source Data file. Created in BioRender. Singh, K. (2026) https://BioRender.com/0otxclw.

### *SCN2A* regulates inhibitory neurogenesis through Sonic hedgehog signaling

To determine the developmental origins of the CGE-like INs and their precocious production in *SCN2A* mutants, we mined our single cell RNA sequencing dataset for ventralizing factors, and found potential alterations to sonic hedgehog (SHH) signaling. SHH signaling is a potent ventralizing factor during neurodevelopment, inducing GABAergic fate in the forebrain^64,65,77,78^. We found elevated SHH expression in both *SCN2A* mutants primarily coming from the excitatory neuron populations (**Fig. 5a**). We hypothesized that there may be a crosstalk effect from excitatory neurons to radial glia given that *SCN2A* is only expressed in neurons and not in progenitors (**Fig. 1f**, and **S1c, d**). We performed cell-cell communication analysis using CellChat^79^ to infer perturbed intercellular communication networks caused by the loss of *SCN2A*. We observed that SHH signaling was a differentially expressed pathway and displayed a shift towards proliferating radial glia in the SCN2A KO organoids (**Fig. 5b**). As we hypothesized, CellChat identified a shift of SHH signaling from excitatory neurons to the proliferating radial glia (**Fig. S16a**). DEG analysis further confirmed significantly elevated expression of the SHH receptor *PTCH1* in proliferating radial glia and reduction in *GLI3*, a negative regulator of the SHH pathway^80^ (**Fig. S16b**). We next validated our transcriptomic data by staining Day 90 organoids for the SHH ligand and GLI3. We observed a significant increase in SHH intensity in both NeuN+ neurons and SOX2+ progenitors (**Fig. 5c, d**). Further, we observed a decrease in GLI3 intensity in both SOX2+ progenitors and SOX2+Ki67+ proliferating progenitors (**Fig. 5e, f**) suggesting decreased negative regulation of SHH signaling in these cells. Lastly, we confirmed elevated extracellular SHH by infecting HEK293T cells with a lentiviral GLI:GFP reporter and treatment with conditioned media from Day 60-70 organoids (**Fig. S16c**). We further report that acute and chronic inhibition of SCN2A using 15 nM PTX-3 elevated SHH signaling (**Fig. S16d-f**). Together, these results indicate that loss of *SCN2A* in excitatory neurons leads to elevated intracellular and extracellular SHH signaling. We next investigated if the elevated SHH signal is driven by excitatory neurons and if the excess IN neurogenesis can be reversed. We transduced *SCN2A^-/-^* organoids with an AAV expressing EGFP and either a scramble shRNA or shRNA against SHH under the CaMKIIa promoter to block the signal (**Fig. S17a**). We found robust expression in neurons after 21 days post-transduction (**Fig. S17b**) and observed reduced SHH expression by immunostaining and our SHH reporter (**Fig. S17c-f**). We observed a reduction in DLX5^+^ INs in both shSHH conditions suggesting that by blocking SHH signal only in excitatory neurons is sufficient to reduce the precocious production of INs (**Fig. S17g, h**). Next, to test if we can rescue the increased production of INs in *SCN2A* mutant organoids by systemically inhibiting SHH signaling, we chronically added 5 μM cyclopamine from D60, where INs begin to be produced^69^, until Day 90 and then labelled INs with AAV-mDLX-GFP (**Fig. 5g**). Cyclopamine treatment led to reductions in GFP^+^ INs in both *SCN2A* mutants (**Fig. 5h, i**). However, cyclopamine treatment did not rescue reduced excitatory neuron numbers (TBR1+ or SATB2+ neurons) in *SCN2A* mutant organoids (**Fig. 5j-l**). Additionally, we investigated other factors influencing ventral fate commitment such as LIF signaling^69^ but found no changes in expression of its effectors such as *LIF* and *LIFR* (**Table S1** and **S2**). Thus, these data indicate that a loss of *SCN2A* in excitatory neurons regulates inhibitory neurogenesis through activation of SHH signaling, primarily driven by neurons, which can be rescued by inhibition of SHH using cyclopamine.

**Fig. 5.**
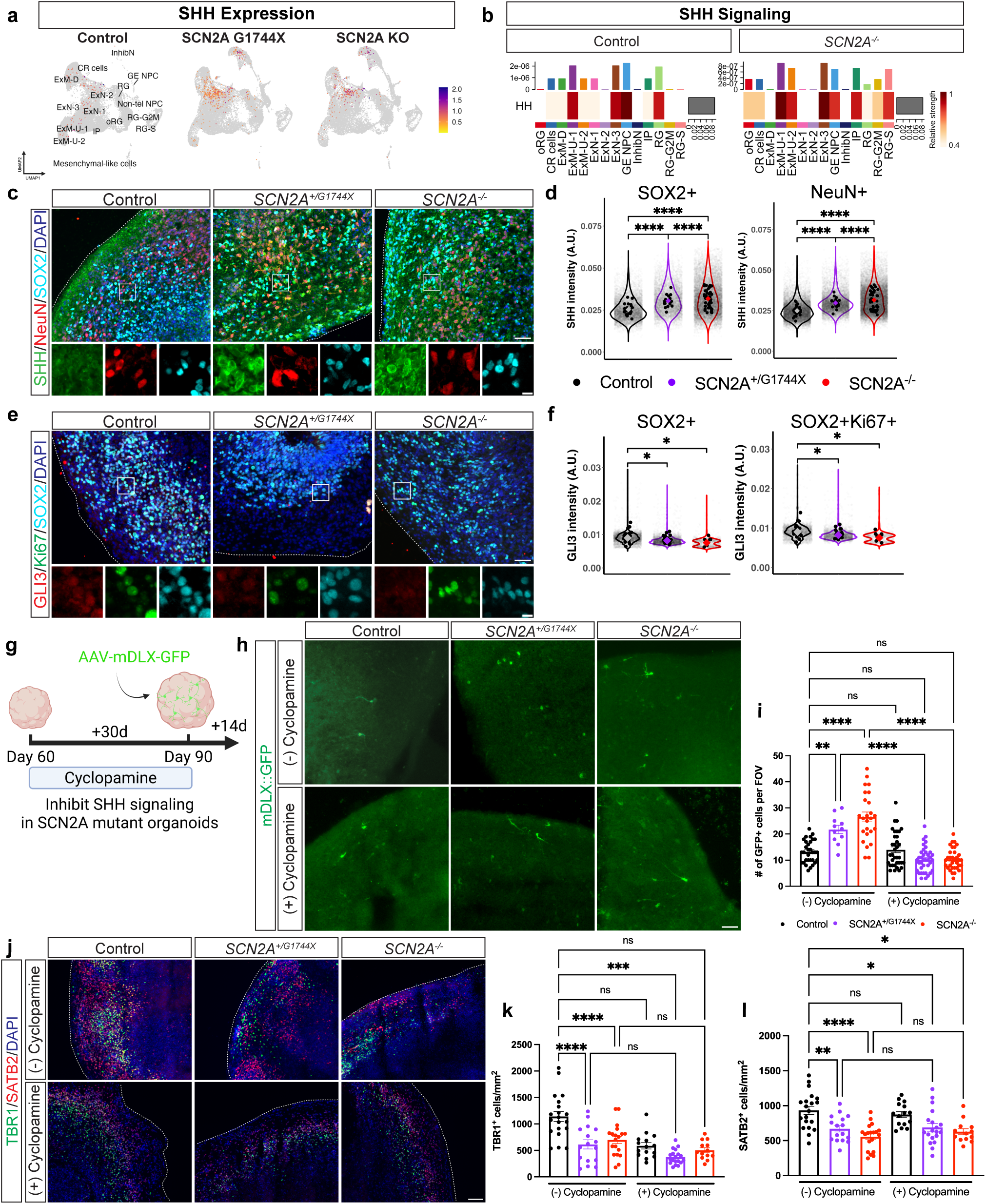
*SCN2A* regulates inhibitory neurogenesis through Sonic hedgehog signaling. **a)** Feature plots showing expression of SHH ligand between genotypes. **b)** Heatmap of overall signaling pattern for the SHH pathway between Control and SCN2A KO organoids. **c)** Immunostaining for SHH, NeuN and SOX2 in Day 90 cerebral organoids. Scale bars = 100 μm, 20 μm. **d)** SHH intensity in SOX2^+^ cells and NeuN^+^ neurons, Day 90 organoids. n = 27/19/42 FOVs for Con-trol/*SCN2A^+/G^*^1744^*^X^*/*SCN2A^-/-^*, from 3 organoids each (1 batch); mixed-effects model (Fixed effect: Genotype, random effect: organoid). SOX2^+^: Genotype and all pairwise p<2.2×10^−1^^6^. NeuN^+^: Genotype and all pairwise p<2.2×10^−1^^6^. Colored diamonds represent the mean of the data, black dots represent FOVs per genotype and transparent dots represent individual cells per FOV. **e)** Immunostaining for GLI3, SOX2 and Ki67 in Day 90 cerebral organoids. Scale bars = 100 μm, 20 μm. **f)** GLI3 intensity in SOX2^+^ cells and SOX2^+^Ki67^+^ proliferating cells, Day 90 organoids. n = 24/25/7 FOVs for Con-trol/*SCN2A^+/G^*^1744^*^X^*/*SCN2A^-/-^*, from 3/3/2 organoids (1 batch); mixed-effects model (Fixed effect: Genotype, random effect: organoid) with Tukey post-hoc. SOX2^+^: χ^2^(2)=9.03, p=0.011; Control vs *SCN2A^+/G^*^1744^*^X^* p=0.029; Control vs *SCN2A^-/-^*p=0.047; *SCN2A^+/G^*^1744^*^X^* vs *SCN2A^-/-^* p=0.78. SOX2^+^Ki67^+^: χ^2^(2)=9.52, p=8.6×10^−3^; Control vs *SCN2A^+/G^*^1744^*^X^* p=0.028; Control vs *SCN2A^-/-^*p=0.034; *SCN2A^+/G^*^1744^*^X^*vs *SCN2A^-/-^* p=0.71. Colored diamonds represent the mean of the data, black dots represent FOVs per genotype and transparent dots represent individual cells per FOV. **g)** Schematic of chronic inhibition of SHH using cyclopamine followed by viral labelling of INs. **h)** Immunostaining for AAV-mDLX-GFP-labelled inhibitory neurons in untreated and cyclopamine-treated organoids. Scale bar = 50 μm. **i)** Inhibitory neuron density in untreated and cyclopamine-treated organoids. n (FOVs, 3 organoids each, 1 batch) for Con-trol/*SCN2A^+/G^*^1744^*^X^*/*SCN2A^-/-^*: untreated 31/11/25, cyclopamine 35/37/33; mixed-effects model (Fixed effect: Genotype and Treatment, random effect: batch) with Tukey post-hoc. Genotype χ^2^(2)=17.14, p=1.9×10^−4^; Treatment χ^2^(1)=0.21, p=0.65; Genotype×Treatment χ^2^(2)=63.64, p=1.5×10^−1^^4^. Tukey: Control (Untreated) vs *SCN2A^+/G^*^1744^*^X^* (Untreated) p=2.5×10^−3^; Control (Untreated) vs *SCN2A^-/-^*(Untreated) p=6.8×10^−1^^2^; *SCN2A^+/G^*^1744^*^X^* (Cyclopamine) vs *SCN2A^+/G^*^1744^*^X^* (Untreated) p=1.7×10^−6^; *SCN2A^-/-^* (Cyclopamine) vs *SCN2A^-/-^* (Untreated) p=7.3×10^−1^^4^. **j)** Immunostaining for TBR1 and SATB2 in untreated and cyclopamine-treated organoids. Scale bar = 100 μm. **k, l)** TBR1^+^ (k) and SATB2^+^ (l) neuron density in untreated and cyclopamine-treated organoids, 2 independent batches. n (FOVs) for Control/*SCN2A^+/G^*^1744^*^X^*/*SCN2A^-/-^*: untreated 20/16/19 from 6 organoids each; cyclopamine 15/19/13 from 3/6/6 organoids; mixed-effects model (same as i) with Tukey post-hoc. (k) Genotype χ^2^(2)=21.06, p=2.7×10^−5^; Genotype×Treat-ment χ^2^(2)=5.82, p=0.055; Tukey: Control (Untreated) vs *SCN2A^+/G^*^1744^*^X^*(Untreated) p=4.4×10^−5^; Control (Untreated) vs *SCN2A^-/-^* (Untreated) p=5.2×10^−6^; *SCN2A^+/G^*^1744^*^X^*(Cyclopamine) vs Control (Untreated) p=2.1×10^−4^. (l) Genotype χ^2^(2)=11.67, p=2.9×10^−3^; Genotype×Treatment χ^2^(2)=1.68, p=0.43; Tukey: Control (Untreated) vs *SCN2A^+/G^*^1744^*^X^*(Untreated) p=4.2×10^−3^; Control (Untreated) vs *SCN2A^-/-^* (Untreated) p=6.2×10^−6^; *SCN2A^+/G^*^1744^*^X^* (Cyclopamine) vs Control (Untreated) p=0.033; *SCN2A^-/-^* (Cyclopamine) vs Control (Untreated) p=0.022. All data are reported as mean ± SEM except for scRNAseq data or unless otherwise stated. Mixed-effects models with fixed effects assessed by Type III Wald χ^2^ tests and pairwise comparisons by two-sided post-hoc Tukey tests. *p < 0.05, **p < 0.01, ***p < 0.001, ****p < 0.0001; n.s., not significant. Source data are provided as a Source Data file. Created in BioRen-der. Singh, K. (2026) https://BioRender.com/0otxclw.

### Loss of SCN2A impairs sodium channel function, neuronal excitability and neuronal network activity

Given that a loss of SCN2A activity leads to an imbalance in excitatory/inhibitory neurogenesis, we sought to assess if there are subsequent functional consequences in neuronal connectivity. We performed whole-cell patch clamp electrophysiology measurements in Day 60 organoid slices (**Fig. 6a**) as we hypothesized early loss of SCN2A function preceded the late neurogenesis deficits. We recorded functional sodium currents in putative pyramidal neurons where we find a decrease in current densities, normalized to membrane capacitance, across *SCN2A* mutant neurons (**Fig. 6b, c**, and **S18a, b**). Further, we find loss of SCN2A leads to a depolarized shift in voltage-dependent activation of sodium currents compared to the isogenic control, as well as a hyperpolarized shift in sodium current inactivation (**Fig. S18c**). These results confirm homozygous loss of SCN2A severely impairs sodium currents and that the heterozygous expression of the G1744X variant reduces sodium current densities to ∼50% of the control as expected in early cortical neurons. We interpret these results with caution as we and others have encountered space-clamp issues that persist in PSC-derived neural cultures^81–86^, however, our results and interpretation remain the same due to genotype-specific effects. We further tested sodium current densities with 5 nM TTX as previously used in our TTX-neurogenesis experiment (**Fig. S7**) and found that per-voltage-step analysis revealed significant reductions in Na⁺ current density across the −50 to 0 mV range in both the *SCN2A^+/G^*^1744^*^X^* and Control + 5 nM TTX conditions compared to Control supporting that 5 nM TTX partially reduces sodium channel activity to levels comparable to the *SCN2A^+/G^*^1744^*^X^* variant (**Fig. S19a-c**). In addition, we find both *SCN2A* mutant neurons in organoids display reduced cell membrane capacitance (**Fig. S18d**). Together, these results suggest loss of SCN2A in early neurodevelopment impairs inward sodium currents and biophysical properties of cerebral organoid neurons which may lead to decreased neuronal excitability.

**Fig. 6.**
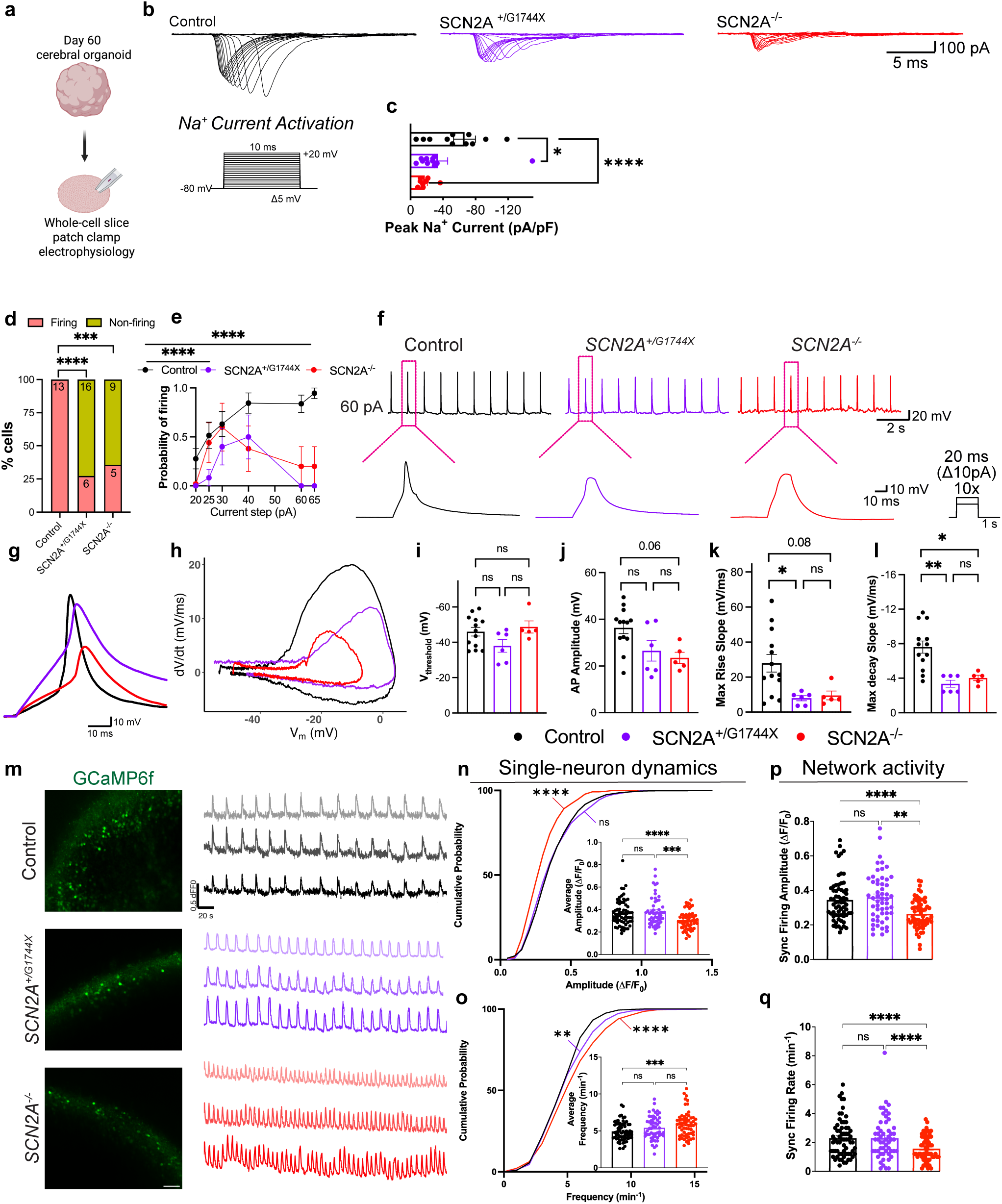
Impaired sodium channel function, neuronal excitability and altered network activity in *SCN2A* mutant cerebral organoids. **a)** Schematic of whole-cell patch clamp electrophysiology in sliced cerebral organoids at Day 60. **b)** Representative sodium current activation traces. **c)** Peak sodium current density normalized to capacitance, Day 60 organoids. n = 12/11/8 neurons for Con-trol/*SCN2A^+/G^*^1744^*^X^*/*SCN2A^-/-^*, from 4/4/3 organoids across 2 independent batches; mixed-effects model (Fixed effect: Genotype, random effect: organoid) with Tukey post-hoc. Genotype χ^2^(2)=18.41, p=1.0×10^−4^; Control vs *SCN2A^+/G^*^1744^*^X^* p=0.047; Control vs *SCN2A^-/-^* p=6.2×10^−5^; *SCN2A^+/G^*^1744^*^X^* vs *SCN2A^-/-^* p=0.10. **d)** Percentage of cells firing an action-potential-like depolarization versus non-firing. n = 13/22/14 neurons for Con-trol/*SCN2A^+/G^*^1744^*^X^*/*SCN2A^-/-^*, from 3 organoids each across 2 independent batches; two-sided Fisher’s exact test, Bonferroni-corrected. Control vs *SCN2A^+/G^*^1744^*^X^* p=5×10^−5^; Control vs *SCN2A^-/-^* p=3×10^−4^. **e, f)** Probability of action-potential-like depolarizations per current step (representative traces at 60 pA in f). n = 13/6/6 neurons for Control/*SCN2A^+/G^*^1744^*^X^*/*SCN2A^-/-^*, from 3 organoids each across 2 independent batches; mixed-effects model (Fixed effect: Genotype and current step, random effect: organoid) with Tukey post-hoc. Genotype χ^2^(2)=76.25, p=2.8×10^−1^^7^; Control vs *SCN2A^+/G^*^1744^*^X^* p=3.3×10^−1^^4^; Control vs *SCN2A^-/-^* p=2.8×10^−6^; *SCN2A^+/G^*^1744^*^X^* vs *SCN2A^-/-^*p=6.4×10^−4^. **g, h)** Representative action potential-like depolarization waveform and phase-plane plots. **i-l)** Action-potential-like depolarization threshold (i), amplitude (j), max rise slope (k) and max decay slope (l); same n and design as in (e, f); mixed-effects model (Fixed effect: Genotype, random effect: organoid) with Tukey post-hoc. (i) χ^2^(2)=5.10, p=0.078; Control vs *SCN2A^+/G^*^1744^*^X^* p=0.23; Control vs *SCN2A^-/-^* p=0.85; *SCN2A^+/G^*^1744^*^X^* vs *SCN2A^-/-^*p=0.13. (j) χ^2^(2)=8.00, p=0.018; Control vs *SCN2A^+/G^*^1744^*^X^* p=0.19; Control vs *SCN2A^-/-^* p=0.064; *SCN2A^+/G^*^1744^*^X^* vs *SCN2A^-/-^* p=0.79. (k) χ^2^(2)=11.11, p=3.9×10^−3^; Control vs *SCN2A^+/G^*^1744^*^X^* p=0.046; Control vs *SCN2A^-/-^* p=0.085; *SCN2A^+/G^*^1744^*^X^* vs *SCN2A^-/-^*p=0.98. (l) χ^2^(2)=22.98, p=1.0×10^−5^; Control vs *SCN2A^+/G^*^1744^*^X^* p=3.0×10^−3^; Control vs *SCN2A^-/-^* p=0.016; *SCN2A^+/G^*^1744^*^X^* vs *SCN2A^-/-^* p=0.85. **m)** Representative fluorescent images of GCaMP6f-infected organoids and colored normalized GCaMP6f intensity traces recorded over 5 minutes of Day 90 organoids. **n, o)** Cumulative distribution and quantification (insets) of amplitude (n) and frequency (o) of spontaneous Ca^2+^ transients, Day 90 organoids. FOV-level n = 65/54/61 for Control/*SCN2A^+/G^*^1744^*^X^*/*SCN2A^-/-^*, from 10 organoids each across 2 independent batches; neuron-level n for KS in (n) 1095/720/1021 and in (o) 998/731/1011. (n) Two-sided KS: Control vs *SCN2A^+/G^*^1744^*^X^* D=0.050, p=0.23; Control vs *SCN2A^-/-^* D=0.203, p<2.2×10^−1^^6^. Inset (mixed-effects model (Fixed effect: Genotype, random effect: batch) with Tukey post-hoc): Genotype χ^2^(2)=23.65, p=7.3×10^−6^; Control vs *SCN2A^+/G^*^1744^*^X^*p=0.99; Control vs *SCN2A^-/-^* p=7.5×10^−5^; *SCN2A^+/G^*^1744^*^X^* vs *SCN2A^-/-^* p=1.0×10^−4^. (o) Two-sided KS: Control vs *SCN2A^+/G^*^1744^*^X^* D=0.087, p=3.5×10^−3^; Control vs *SCN2A^-/-^* D=0.143, p=1.9×10^−9^. Inset: Genotype χ^2^(2)=16.31, p=2.9×10^−4^; Control vs *SCN2A^+/G^*^1744^*^X^* p=0.076; Control vs *SCN2A^-/-^* p=1.7×10^−4^; *SCN2A^+/G^*^1744^*^X^* vs *SCN2A^-/-^* p=0.22. **p, q)** Synchronous firing amplitude (p) and frequency (q), Day 90 organoids (n and design as in n, o); mixed-effects model (Fixed effect: Genotype, random effect: batch) with Tukey post-hoc. (p) Genotype χ^2^(2)=21.45, p=2.2×10^−5^; Control vs *SCN2A^+/G^*^1744^*^X^* p=0.91; Control vs *SCN2A^-/-^* p=7.0×10^−5^; *SCN2A^+/G^*^1744^*^X^* vs *SCN2A^-/-^*p=1.0×10^−3^. (q) Genotype χ^2^(2)=32.50, p=8.7×10^−8^; Control vs *SCN2A^+/G^*^1744^*^X^* p=1.0; Control vs *SCN2A^-/-^* p=1.6×10^−6^; *SCN2A^+/G^*^1744^*^X^* vs *SCN2A^-/-^* p=5.9×10^−6^. Data are reported as mean ± SEM. Patch-clamp and average calcium-transient quantifications were analysed with linear or generalized linear mixed-effects models; the Genotype fixed effect was assessed by Type III Wald χ^2^ tests and pairwise comparisons by two-sided post-hoc Tukey tests; firing/non-firing proportions (d) used two-sided Fisher’s exact tests with Bonferroni correction; cumulative distributions (n, o) used two-sided Kolmogorov–Smirnov tests. *p < 0.05, **p < 0.01, ***p < 0.001, ****p < 0.0001; n.s., not significant. Source data are provided as a Source Data file. Created in BioRender. Singh, K. (2026) https://BioRender.com/0otxclw.

Given reduced sodium currents produced by both SCN2A mutants, it is likely that there are impairments in neuronal excitability. We examined the repetitive firing and action potential properties of *SCN2A* mutant neurons. While we find no difference in resting membrane potential (**Fig. S18e**), we found a significant inability of *SCN2A* mutant neurons to fire action potential-like depolarizations as well as a reduced probability to fire multiple action potential-like depolarizations in response to increasing current injection steps (**Fig. 6d-f**, see Methods). We next examined the action potential properties using phase-plane analysis (**Fig. 6g, h**), which compares the rate of change in voltage during action potentials to the voltage^87^. While we find no difference in threshold potential or rheobase (**Fig. 6i** and **S18f**), we find reduced action potential-like depolarization amplitude in *SCN2A^-/-^* neurons but not *SCN2A^+/G^*^1744^*^X^* neurons (**Fig. 6j**). Interestingly, both *SCN2A* mutants display reduced maximum action potential-like depolarization rise and decay slopes (**Fig. 6k, l**) while only *SCN2A^+/G^*^1744^*^X^*neurons display increased rise and decay times (**Fig. S18g, h**). Thus, complete or partial deletion of *SCN2A* impairs neuronal excitability and action potential kinetics in cerebral organoid neurons.

We next examined neuronal network activity using live Ca^2+^ imaging in Day 90 organoids using AAV-hSyn-GCaMP6f under basal conditions (without stimulation, **Fig. 6m**). Organoids were infected two weeks prior to imaging and maintained in BrainPhys^TM^ media one week prior to promote neuronal activity^88,89^. We measured the amplitude and frequency of Ca^2+^ transients that represent spontaneous activity of individual GCaMP6f-positive neurons. Further, we measured synchronous firing which indicates the frequency of groups of neurons firing together, thereby eliciting synchronized activity. The spontaneous calcium transients obtained from GCaMP6f-infected neurons were also TTX-sensitive (**Fig. S20a-d**). Single-neuron analysis revealed no difference in amplitude and frequency between Control and *SCN2A^+/G^*^1744^*^X^*, however, *SCN2A^+/G^*^1744^*^X^* neurons displayed a shift towards a higher frequency distribution (**Fig. 6n, o**). In *SCN2A^-/-^* neurons, there was a reduction in amplitude but an increase in calcium transient frequency (**Fig. 6n, o**). *SCN2A^+/G^*^1744^*^X^* neurons showed no change in baseline network activity but *SCN2A^-/-^* neurons displayed reductions in both synchronous firing rate and amplitude (**Fig. 6p, q**). Together, these data suggest that a complete or partial loss of *SCN2A* causes early deficits in sodium currents and neuronal excitability that likely lead to abnormal calcium dynamics at the single-neuron and network levels in mature organoids.

## Discussion

*SCN2A* is known for its postnatal functions in regulating neuronal excitability and action potential generation^2,15–17,19,21,23–25^. While prior studies characterized a role for *SCN2A* in early postnatal and mature neurons, if and how *SCN2A* regulates earlier stages of prenatal development remain unknown given it is expressed at early stages. Here, we provide evidence supporting a neurogenic role for *SCN2A* in early (prenatal) neurodevelopment using human neural organoid models generated from isogenic and patient-derived PSC lines and define its cellular and molecular origins. Our study indicates that *SCN2A* regulates the timing of the genesis of excitatory and inhibitory neurons through modulation of SHH signaling. Further, we show loss of SCN2A severely impacts neuronal sodium currents and action potential generation. Lastly, our work also highlights that ASD-linked variants in *SCN2A* may impair brain development much earlier than previously known, altering neurogenesis prior to modulating fast synaptic transmission in the postnatal brain. Together, these results outline a previously unappreciated role for *SCN2A*’s contribution and timing toward ASD pathogenesis and the role of sodium channels in neurogenesis.

There is a growing body of evidence suggesting that some ASD risk genes, classically known for their role in postnatal neuronal communication, may regulate early developmental mechanisms. For example, the synaptic protein *SYNGAP1* regulates radial glial cell proliferation and neuronal maturation in cortical organoids^90^. Further, the synaptic organizer *NRXN1* displayed cell type-specific transcriptional impairments and vulnerabilities in organoid-derived neural progenitor cells^66^. These studies highlight the potential unexpected roles of ASD risk genes involved in neuronal communication in early brain development. In addition, the neural organoid model has allowed for the investigation of prenatal brain development of other neurodevelopmental risk genes such as *CHD8*^63,89^, *PACS1*^67^, *KIF26A*^91^, *ARID1B*^68^, and others^76,81,92–94^. In the case of *SCN2A*, its expression is well established in mature neurons^2,11,16,17,20^, however, transcriptomic data from human prenatal cortex and neural organoids indicate *SCN2A* is expressed at prenatal time points^26,29,33^. We thus hypothesized that complete or partial loss of *SCN2A* impairs early neurodevelopment by disrupting neuronal maturation. We show using human neural organoids that loss of *SCN2A* resulted in progressive deficiencies in the production of early-born excitatory deep layer neurons and late-born upper layer neurons. These shifts in excitatory neuron populations were concomitant with an increase in CGE-like INs supporting a neurogenesis-derived shift in excitatory-inhibitory (E-I) balance. Taken together, our findings contribute to the growing evidence of unexpected roles of ASD risk genes in early neurodevelopment.

Using single-cell transcriptomics, we uncovered several impairments in the excitatory neuron development of both *SCN2A* mutant organoids. Our analyses revealed vulnerabilities in multiple neural cell types with an emphasis on the maturing excitatory neurons. Unexpectedly, we found that across organoid development, there was impaired production of both deep and upper layer neurons. This finding supports a previous large scale clinical study identifying nearly 25% of individuals with *SCN2A* mutations with microcephaly^95^ and a subset having developmental brain malformations^96^, suggesting VGSCs such as *SCN2A* may influence neuronal production and migration. Similarly, pathogenic variants in *SCN3A* have been shown to impair cortical folding and neuronal migration^31^ underscoring the role of VGSCs in brain development. It is important to note, we did not find any expression of *SCN2A* in radial glial cells and thus these effects are likely to be non-cell autonomous by *SCN2A* mutant neurons. We further did not find changes in cell cycle exit of radial glia suggesting a change in neuronal identity (upon differentiation) rather than accelerated differentiation^97,98^. This notion is also supported by the lack of cell death observed in mature organoids. We note that we use the term neurogenesis throughout this study to refer broadly to the production and specification of neuronal subtypes from progenitor cells, rather than progenitor proliferation per se. Our data indicate that SCN2A does not regulate progenitor cell division or survival, but rather influences the fate of neurons generated from those progenitors through a non-cell-autonomous mechanism. In addition, through pharmacological blockade of sodium channels using TTX or PTX-3, we observed a reduction in early neuron production suggesting that early neural activity driven by sodium channels may regulate neurogenesis without changes in cell death. A central question arises as to what downstream pathways are directly regulated by *SCN2A* that govern those neurogenic programs. Our study and others have demonstrated that loss of *SCN2A* can disrupt spontaneous and evoked calcium transients ^14,15,18^ and given the importance of calcium activity in neurodevelopment^99^, in particular action potential-evoked calcium events, it is possible that abnormal neurogenesis phenotypes are initiated in part by dysregulated calcium activity. Spontaneous calcium transients have been shown to occur prenatally and perturbation of this activity can lead to impaired circuit formation and neuronal migration^100^. Further, it has been shown that hyperpolarization of neural progenitor cell membrane potential changes the laminar position of their postmitotic daughter cells^101^. Thus, altering the activity profile or membrane properties of early cortical cells can impair normal brain development by disrupting the tempo of neurogenesis.

The loss of *SCN2A* resulted in a striking E-I shift in cerebral and dorsal forebrain organoids where a reduction in excitatory neurons were concomitant with precocious production of INs. First, we identified the INs on the basis of their expression of ventral lineage markers such as the DLX transcription factors^102^. We further identified the INs as CGE-like given their similar expression profiles to CGE-derived INs and lack of MGE marker Nkx2-1. A majority of embryonic INs are derived from the ventral forebrain in the ganglionic eminences^103^, however, a subset of dorsal forebrain progenitors have been shown capable of producing GABAergic INs^104^. Recent studies using single-cell lineage tracing using human prenatal cortical tissue demonstrated that dorsal-derived progenitors were capable of producing both excitatory and inhibitory neurons^73^ whereas in mouse cortical progenitors were largely restricted to a single lineage^74^. Furthermore, analysis of adult human brain identified a substantial portion of dorsal cells have the potential to differentiate into both cortical excitatory and inhibitory neurons^105^. Lastly, LIF signaling can induce inhibitory neurogenesis in cortical tissue and organoids with CGE-like identity^69^. A remarkable finding between these studies is that the INs produced most resembled CGE-derived INs, similar to the INs produced in cerebral organoids in this study. This implies that dorsal forebrain progenitors have the intrinsic potential to differentiate into GABAergic CGE-like INs, and *SCN2A* may regulate this process through SHH signaling. Lastly, inhibition of SHH signaling by cyclopamine or genetic inhibition reversed the increased production of INs in SCN2A mutant organoids but did not rescue deficits in excitatory neuron production. We speculate that the reduced neuronal activity in newborn excitatory neurons may have a negative-feedback effect on progenitors to reduce excitatory neurogenesis, potentially independent or upstream of elevated SHH signaling that causes precocious inhibitory neurogenesis. It is also possible that reduced neuronal activity may interrupt the normal developmental sequence of cortical neurogenesis through a parallel signaling program. For example, it has been shown that inhibition of SHH signaling in early cerebral organoids reduces upper layer neuron production^106^ suggesting SHH signaling can directly affect dorsal cell fate. Interestingly, we observed a decrease in GLI3 mRNA and protein levels in *SCN2A* mutant radial glia. GLI3 has been shown to be a negative regulator of SHH signaling and a determinant of dorsal fate through regulation by Wnt signaling^107^. While we did not directly examine Wnt signaling, there may be some interplay between Wnt and SHH signaling in which loss of SCN2A disrupts this balance which disrupts cortical fate decisions. Future studies will need to determine how *SCN2A* regulates cortical neurogenesis and whether this is directly causing changes to SHH signaling or another causal pathway.

Our study reveals a link between neuronal activity during prenatal brain development and SHH signaling. While SHH signaling has been shown to mediate neural plasticity in the postnatal brain^108^, our study indicates interaction between these pathways at a much earlier time point. We posited that aberrant neuronal activity caused by loss of *SCN2A* is upstream of the neurogenic defects. We previously demonstrated that mature *SCN2A* KO neurons at baseline are hypoexcitable but are intrinsically hyperexcitable and capable of firing high-frequency bursts of action potentials^23^. In this study, we further show that embryonic excitatory neurons lacking SCN2A display impaired sodium currents and action potential-like depolarization properties as well as INs display mild deficiencies by calcium activity. While our electrophysiology data suggest a hypoexcitable nature, our calcium data suggest that some *SCN2A* mutant excitatory neurons are capable of producing higher frequency calcium events which could have an influence on morphogen signaling. Interestingly, *SCN2A* mutant CGE-like INs display reduced calcium transient frequency which may reflect reduced excitatory drive onto these neurons. Early developmental GABAergic signaling can influence neurogenesis^109^ and E-I balance^110^, and therefore reduced IN activity could have downstream consequences for cortical circuit maturation. How exactly this altered E-I balance impacts overall network dynamics in organoids remains to be fully characterized. Nevertheless, this shift could be a transient effect during prenatal brain development and may be resolved later in postnatal development, given the *Scn2a* heterozygous mouse model does not display excess numbers of GABAergic neurons in the adult brain^15^. However, given CGE-derived INs have not been examined in *Scn2a*-deficient mice, future studies should investigate the functional properties of these neurons during development and how they influence excitatory neuron activity. Alternatively, the effect with *SCN2A* and neurogenesis could be a human-enriched event given the *SCN2A* mutants produced CGE-derived INs from dorsal progenitors^69,73,105^. Lastly, our pharmacological experiments using the pan-Na_v_ blocker TTX and selective Na_v_1.2 blocker PTX-3 demonstrate an underappreciated role for sodium channels in regulating neurogenic events. The results of this study open up the possibility of differential regulation of neuronal activity and neurogenesis with a potential link to downstream events such as gliogenesis in the embryonic brain.

There remain some limitations to the current study. First, while we captured mostly dorsal forebrain development, there lacks the ventral forebrain that produces a large portion of interneurons of the cortex^70^ and therefore does not fully recapitulate brain development in that aspect. The use of the assembloid model, composed of regionalized neural organoids, such as dorsal-ventral^64,78,111^ or thalamocortical^92,112^, would more faithfully recapitulate normal brain development. Additionally, other critical cell types are not present such as microglia. Microglia in organoids have been shown to modulate the maturation and development of the neural cell types^113–115^. Of note, it has been shown that microglia in Scn2a-deficient mice present abnormal pruning of synapses and their ablation can improve synaptic transmission deficits. While cerebral organoids lack microglia, we cannot rule out the possibility of their contribution to modulate excitatory neurons or progenitors. Furthermore, recent work has raised questions regarding the selectivity of PTX-3 for Na_v_1.2 over Na_v_1.6^116^. However, our scRNA-seq data indicate that SCN8A (Na_v_1.6) expression is minimal in excitatory neurons at the developmental stages examined, with SCN2A being the predominant neuronal sodium channel, making off-target effects of PTX-3 on Na_v_1.6 unlikely in our system. Lastly, we find that GABAergic CGE-like INs in SCN2A mutant organoids show decreased activity by calcium imaging. In the mouse brain, CGE-derived INs receive inputs from other excitatory neurons of the cortex and synapse onto MGE-derived somatostatin or parvalbumin-expressing INs^117–119^. As previously mentioned, the cerebral or dorsal forebrain organoid model lack IN diversity from the MGE, the majority embryonic source of INs in the cortex and therefore the normal synaptic targets of CGE neurons are not present which may affect overall system activity. Furthermore, the shared genesis of excitatory and CGE-like inhibitory neurons from dorsal progenitors appears to be present in humans but not the mouse^73,74^ and thus how *Scn2a* regulates neurogenesis in the mouse brain may be different than what occurs in humans. Future studies should also address the alternative sources of SHH in *SCN2A* mutant organoids and whether its effects are downstream or parallel to the excitatory neurogenic defects. Additionally, SHH has been demonstrated to contribute to epileptiform activity while its suppression can reduce that elevated activity^120^. While we did not address whether excitability was impacted by blockade of SHH signaling with cyclopamine, it remains pertinent for future studies to address these neurophysiological features.

In summary, our study demonstrates an unexpected embryonic neurogenic role for *SCN2A* by regulating cortical excitatory-inhibitory balance through SHH signaling. We show that loss of *SCN2A* can lead to abnormal effects on neurogenesis and that these effects are likely to be non-cell autonomous by *SCN2A* mutant neurons. Our results may provide an explanation to circuit dysfunction caused by loss of *SCN2A* that stem from abnormal neuron development from the prenatal stages.

## METHODS

### Human pluripotent stem cell culture

This research was performed in compliance with relevant ethical regulations and approved by the University Health Network Research Ethics Board (21-5860). All human iPSC lines (with the exception of the 19-2-2 *SCN2A^+/G^*^1744^*^X^*) used were generated in previous studies^23,58^. Generation of isogenic *SCN2A* mutant ESC lines were derived from the commercially available wildtype H1 ESC line (WA01, WiCell). All iPSC/ESC lines are male-derived. All experiments performed using the above stem cell lines were cultured at 37°C in a humidified incubator on Matrigel-coated plates (Corning) in mTeSR1 medium (STEMCELL Technologies). Cells were passaged using ReLeSR (iPSCs, STEMCELL Technologies) or Versene (ESCs, ThermoFisher).

### CRISPR/Cas9 gene editing

For generating *SCN2A^+/G^*^1744^*^X^*cells, H1 hESCs (WiCell) were genome edited with dual sgRNA:Cas9 RNP complexes to insert a single nucleotide edit of c.5230C>T in the *SCN2A* gene (gRNA 1: GACCCTGACAAAGATCACCC, gRNA 2: GCTCAGTTAAAGGAGACTGT). DNA oligos that included Alt-R ™ sgRNA (IDT) and Alt-R™ HDR single stranded oligodeoxynucleotide (ssODN: GCA CCT ATT CTT AAT AGT GGA CCT CCA GAC TGT GAC CCA GAC AAA GAC CAC CCT GGA AGC TCA GTT AAA TGA GAC TGT GGG AAC CCA TCT GTT GGG ATT TTC TTT TTT GTC AGT TAC ATC) (IDT) were prepared to a final concentration of 10 μM in nuclease-free water (Corning). On the day of nucleofection, the RNP complex was prepared by creating a mastermix of 1 μL of 61 μM stock HiFi Cas9 (IDT), 11 μL of 10 μM sgRNA, and 100 μL of nucleofector solution with its’ supplement (P3 Primary Cell 4D-Nucleofector X Kit, Lonza) and allowed to incubate for 20 minutes. After incubation, 22 μL of 10 μM ssODN was added to the RNP mastermix. H1 ES cells were then dissociated into single cells using Accutase (STEMCELL Technologies), centrifuged and resuspended in mTeSR1

+ Y-27632. The cells were counted using an automated cell counter (Cellometer Auto T4, Nexcelom) and added to a Lonza nucleofector cuvette at 1,000,000 cells in nucleofector solution. The cuvette was placed in a 4D-Nucleofector X Unit (Lonza) using program ‘CA 137’ for nucleofecting PSCs. The cells were allowed to incubate for 10 minutes post-nucleofection at room temperature. Next, nucleofected cells were plated onto 6-well Matrigel-coated wells at 500,000 cells per well in mTeSR1 + Y-27632. The plates were placed in a humidified incubator at 37°C for 24 hours. This same method was used to generate the *SCN2A^+/G^*^1744^*^X^* in the 19-2-2 iPSC line background.

For generating *SCN2A^-/-^* cells, H1 hES cells (WiCell) were genome edited with dual sgRNA:Cas9 RNP complexes targeted to exon 2 to create a frameshift deletion in exon 3 of the *SCN2A* gene (gRNA 1: GAGACCCAAACAGGAACGCAAGG, gRNA 2: CCTTGCTGCTATTGAACAACGCA). The same procedure for nucleofection was followed as described above with the exception of no ssODN was used.

### Purification of CRISPR-edited cells

Nucleofected cells were allowed to grow to 70% confluency where single colonies were picked under a brightfield microscope using a P20 pipette and plated in individual wells of a Matrigel-coated 24-well plate. Once single colonies had reached confluency, the cells were split 1:2 into a new well of a 24-well plate with each well having a sister well. Next, after the cells reached 50% confluency, half the plates were harvested for droplet digital PCR (ddPCR). The cells were lifted using Accutase and neutralized with DNA lysis buffer (10 mM Tris pH 7.5, 10 mM EDTA pH 8.0, 10 mM N-lauroylsarcosine (Millipore Sigma), and freshly added 1 mg/ml proteinase K). The cells were then gently transferred to a 96-well PCR plate containing equal volume DNA lysis buffer and sticker sealed. The sealed PCR plate was incubated in a thermocycler (Bio-Rad) at 70°C for 10 minutes. The plate was then chilled on ice for 4 minutes and 100 μL/well of DNA precipitation solution (95% EtOH, 5% H20, and 75 mM NaCl chilled at −80°C) was added to each well. The PCR plate was then left at room temperature for 1 hour then centrifuged at 1800 x g for 4 minutes. The plate was then flicked quickly upside down to remove the supernatant. Two gentle washes were performed with 70% EtOH at 100 μL/well and flicked in between each wash. The PCR plate was then placed back into the thermocycler unsealed and incubated at 70°C for 3 minutes to evaporate the excess EtOH. DNA was resuspended in 20 μL of UltraPure Distilled Water (ThermoFisher), heated at 70°C for 10 minutes and vortexed briefly. Custom Taqman MGB probes (ThermoFisher) were designed and optimized for detection of our gene targets (6FAM probe to detect the mutant allele and a VIC probe to detect the wildtype allele) based on Bio-Rad’s ddPCR manual recommendations. Primers for ddPCR were designed to bind outside the ssODN region. ddPCR was performed by The Centre for Applied Genomics at The Hospital for Sick Children. Once the ddPCR data was received, the well with the highest edited frequency by the FAM probe were selected and replated as single cells supplemented with CloneR^TM^2 (STEMCELL Technologies) as per the manufacturer’s protocol. Once the single cell colonies had grown after 3-4 days, they were passaged into smaller colonies and plated into a new 24-well plate with each well having a sister well. This process was repeated as described until we obtained a pure population of edited cells.

### Validation of CRISPR-edited cells and authentication

Once we had achieved our desired edited cells based on ddPCR data, we validated the cells using ddPCR or sanger sequencing. For *SCN2A^-/-^* cells, sanger sequencing was performed using primers flanking the sgRNA target site at exon 2 to confirm a 50 bp indel edit. For *SCN2A^+/G^*^1744^*^X^*cells, copy number analysis was performed using ddPCR with the autosomal RNaseP gene which appears at 2 copies. Secondly, sanger sequencing using primers flanking the edited region on exon 27 were used to confirm the heterozygosity of the edit. Off-target effects were detected using sanger sequencing using primers targeting the top 2 off-target genes provided by Benchling as well as for SCN3A and SCN8A. Confirmed positive clones were expanded and subjected to downstream validation steps. Once each cell line had been validated by sequencing, they were subjected to karyotyping, short-tandem repeat (STR) profiling (The Centre for Applied Genomics) and mycoplasma testing. See Table S7.

### Generation of cerebral organoids

Cerebral organoids were derived from hPSC cultures using the STEMdiff^TM^ Cerebral Organoid Kit (STEMCELL Technologies, cat #08570) based on the protocol developed by Lancaster et al^36^. Cerebral organoids were generated according to the manufacturer’s protocol. In brief, to generate embryoid bodies (EBs), hPSCs are grown to 70-80% confluency in a 6-well plate and dissociated using the Gentle Cell Dissociation Reagent at 37°C for 8-10 minutes. The cells were gently resuspended and transferred to a 15ml conical tube and centrifuged at 300 x g for 5 minutes. The cells were then resuspended in EB Seeding Medium (EB Formation Medium + 10 μM Y-27632) and counted using a Cellometer Auto T4 (Nexcelom) and viability tested using Trypan Blue staining. The cells were then plated onto a 96-well ultra-low attachment plate (Corning) in EB Seeding Medium at 9000 cells/well and centrifuged at 100 x g for 3 minutes. The cells were then placed in a 37°C incubator for 24 hours with media changes at day 2 and 4 using EB Formation Medium. On day 5, EBs were transferred into individual wells of a 24-well ultra-low attachment plate (Corning) containing Induction medium and incubated for 48 hours. On day 7, Matrigel (Corning) was thawed on ice for 1-2 hours. Individual EBs were embedded into 20 μL Matrigel droplets on Organoid Embedding Sheets (STEMCELL Technologies) and incubated at 37°C for 25 minutes to allow the Matrigel to polymerize. Matrigel droplets containing EBs were then washed into a 6-well ultra-low attachment plate (Corning) containing Expansion Medium. Embedded organoids were then incubated at 37°C for 3 days. On day 10, organoid media was changed to Maturation Medium and placed in a 37°C incubator on a Celltron orbital shaker (INFORS HT) at 65 rpm. Media was changed every 3-4 days until organoids reached the specified experimental timepoint.

### Generation of dorsal forebrain organoids

Dorsal forebrain organoids derived from H1-ESCs were generated using the STEMdiff™ Dorsal Forebrain Organoid Differentiation Kit (STEMCELL Technologies, cat #08620) based on the protocols developed by the Pasca Lab^64,75,111^.

### Tissue processing for immunostaining

Organoid samples were fixed in 4% paraformaldehyde shaking at 4°C overnight. Organoids were then incubated in PBS containing 30% sucrose overnight. The organoids were then incubated in gelatin solution (7.5% w/v gelatin, 10% sucrose, PBS) for 1 hour at 37°C then placed in embedding molds containing gelatin solution and snap frozen using liquid nitrogen and placed in a −80°C freezer for later cryosectioning. Cryosections were cut at 16 μm thickness unless otherwise stated.

### Immunostaining

Organoid sections were washed 3 x 10 minutes with 1X PBS + 0.1% Tween 20 (PBS-T) and blocked and permeabilized in 10% normal donkey serum + 10% Triton-X-100 in PBS (blocking/permeabilization (BP) solution) for 1 hour at room temperature. Organoid sections were then incubated with primary antibodies in BP solution overnight. The next day, sections were washed 3 x 20 minutes in PBS-T and incubated for 2 hours at room temperature with fluorophore-conjugated secondary antibodies at 1:500 dilution then washed for 3 x 20 minutes in PBS-T. DNA nuclear counterstaining was performed using 300 nM DAPI for 10 minutes and washed for 20 minutes in PBS-T. Slides were mounted using Prolong Gold antifade mountant. Primary antibodies and dilutions are listed in the reporting summary.

The following antigens required antigen retrieval: FOXG1, SATB2, CTIP2, BrdU, and Ki67. Antigen retrieval was performed by placing section slides in a food steamer (Hamilton-Beach 37530) in citrate buffer (10 mM tri-sodium citrate dihydrate, 0.05% Tween 20) for 20 minutes followed by 3 x 10-minute washes in PBS-T then continued onto the immunostaining workflow above.

### Bromodeoxyuridine labelling in cerebral organoids

Organoids were transferred to a new 6-well ultra-low attachment plate with maturation media supplemented with 10 μM BrdU (Sigma) and incubated at 37°C for 1 hour. Organoids were then washed 3 times with 1X PBS and incubated in maturation media for a further 24 hours and then processed for immunostaining.

### Tetrodotoxin and Phrixotoxin-3 treatment in cerebral organoids

Control organoids were transferred to a new 6-well ultra-low attachment plate with maturation media supplemented with 5 nM TTX or PTX-3 (Tocris) or water and incubated at 37°C on Day 30 until Day 40 or 60. Media was changed every 3-4 days with fresh drug or water added. At Day 40 and 60, organoids were then washed 3 times with 1X PBS then processed for immunostaining.

### Sholl analysis

For AAV-mDLX-GFP infected organoids, confocal images were taken on a Zeiss LSM 880 at a resolution of 2048 x 2048 pixels using a Plan-Apochromat 63x/1.4 oil objective and analyzed with ImageJ2 (v2.14.0/1.54f). Sholl analysis was performed using the Simple Neurite Tracer (SNT) plugin^121^. Briefly, concentric rings are drawn radiating from the center of the neuron soma with a constant increasing 5 μm radius step size and then dendritic intersections are counted. Neurons with at least 3 primary neurites were analyzed.

### Image processing and quantification

Images for analysis and figures were manually adjusted for brightness/contrast in ImageJ and exported as .tif format. All organoid images with the exception of the ones used for sholl analysis were taken on a fluorescence microscope (Revolve, ECHO). All pixel units were converted to millimeters using ImageJ2.

For Day 40 TBR1 progenitor zone analysis, CellProfiler (v4.2.6)^122^ was used to quantify nuclear markers. A region of interest (ROI) was drawn around the progenitor zone using SOX2 as a guide. Next, a gaussian filter (*sigma* = 1) was applied to each image and a mask was drawn using the ROI on the input image. The *IdentifyPrimaryObjects* module was used to identify nuclei using the following criteria: *diameter* = 20 to 40 pixel units, global threshold strategy, Otsu thresholding for 3 classes, threshold smoothing scale of 4, and threshold correction factor of 0.9. The *CalculateMath* module was used to determine the percentage of TBR1^+^ cells within the SOX2^+^ progenitor zone.

For Day 40 cell cycle exit analysis with BrdU, we used a modified CellProfiler pipeline to analyze cell cycle exit. An ROI was drawn around a progenitor zone using SOX2 as a guide. Next, a gaussian filter (*sigma* = 1) applied to each image channel and a mask was drawn using the ROI. The *IdentifyPrimaryObjects* module, with the same criteria as the TBR1 progenitor zone analysis, was applied to each nuclear antigen: BrdU, Ki67, SOX2, and DAPI. Next, to identify double positive cells (ie. BrdU^+^Ki67^+^), the *RelateObjects* module and *FilterObjects* module were used with BrdU as the parent object and Ki67 as the child object. To classify the cells as either single or double marker positive, we used the *ClassifyObjects* module as set BrdU to be the object to be classified. Lastly, we used the *CalculateMath* module to calculate the percentage of BrdU^+^Ki67^-^ over the total number of BrdU^+^ cells. A similar process was applied for identifying CC3+ neuronal (NeuN+) or progenitor (SOX2+) cells in TTX-treated organoids.

For Day 60 and 90 nuclear staining, we used a modified CellProfiler pipeline to count the number of marker-positive cells over the area measured. An ROI was drawn around the organoid (denoted by the dotted line in the figures) and the previously mentioned workflow for Day 40 TBR1 progenitor zone analysis was used for each nuclear marker (TBR1, SATB2), with the exception that diameter size ranged from 16-40 pixel units. To calculate the nuclei marker density, the total number of nuclei marker was divided by the ROI area measured in mm^2^. The same process was applied for Figure 5i-k for cyclopamine-treated organoids.

For counting GFP^+^ cells in organoids infected with AAV-mDLX-GFP, we manually counted neurons in each field of view (FOV) using the Cell Counter plugin in ImageJ. For isogenic and patient-derived iPSC organoids, images were zoomed in from 4X objective images for visibility.

For Day 90 SHH intensity staining, we used a modified CellProfiler pipeline to identify neuronal (NeuN+) or radial glial nuclei (SOX2+) using the *IdentifyPrimaryObjects* module. SHH+ objects were then identified using the *IdentifySecondaryObjects* module for the respective nuclei and average cell intensity was measured using the *MeasureObjectIntensity* module. Similarly, for GLI3 intensity staining, progenitor nuclei and GLI3+ objects were identified using the *IdentifyPrimaryObjects* module. GLI3+ progenitor cells were identified using *RelateObjects* and *ClassifyObjects* and average GLI3 intensity per cell was measured using *MeasureObjectIntensity*.

### Cell lysis and mass spectrometry

Cerebral organoids were matured to Day 90 and prepared for mass spectrometry and analyzed at SPARC Biocentre, Hospital for Sick Children, Toronto, Canada. Raw mass spectrometry data was analyzed and statistics calculated using Spectronaut™ (Biognosys). Briefly, DIA analysis and quantification were done using Spectronaut 18.6 (Direct DIA+). Database searched against Uniprot_UP000005640_Human_SP_20231026 (20411 Entries) with the following parameters: Enzyme: Trypsin Max Missed Cleavages: 3 Fixed Modifications: Carbamidomethylation: C (+57.02) Variable Modifications:Oxidation: M (+15.99) Acetylation: peptide N-term (+42.01)

### Prenatal brain gene expression visualization

Visualization of the spatiotemporal expression pattern of *SCN2A* was performed by taking the expression data from the BrainSpan database^28^. The data was plotted using the cerebroviz package (v1.0)^123^.

### *SCN2A* gene expression analysis of published transcriptomic datasets

To view *SCN2A* expression in human mid-gestation prenatal cortex single-cell RNAseq data, we used the data from Polioudakis et al.^29^ available on their web portal (CoDEx, http://geschwindlab.dgsom.ucla.edu/pages/codexviewer). To view *SCN2A* expression in cerebral organoids, we used the data from Kanton et al.^33^ available on their web portal (scApeX, https://bioinf.eva.mpg.de/shiny/sample-apps/scApeX/). For expression patterns in the adult human cortex, we used the Allen Brain Atlas’s Transcriptomic Explorer with the Human M1 – 10X Genomics (2020) dataset from Hodge et al.^34^

### Dissociation of organoids for fixed-cell single-cell RNA sequencing

Organoids were dissociated as previously described^124^ using the Papain Dissociation Kit (Worthington) per the manufacturer’s protocol. In brief, organoids were transferred to a 6-well plate containing 2.5 mL of Papain + DNase solution. A sterile razor blade was used to mince organoids in small pieces. Minced organoids were transferred to an orbital shaker at 65 rpm inside a humidified incubator at 37°C and 5% CO2 for 30 minutes. After, organoids were gently triturated using a P1000 pipette and placed back into the orbital shaker for 10 more minutes. After incubation, a 10 mL serological pipette was used to gently break up the minced organoids further. The entire well volume was transferred to a 15 mL conical tube and debris were allowed to settle for 1-3 minutes. The cell suspension was then transferred to a new conical tube containing 5 mL of Earle’s medium plus 3 mL of reconstituted ovomucoid inhibitor, inverted 3-5 times and centrifuged at 300 x g for 7 minutes. The supernatant was aspirated, and the cell pellet was gently resuspended in 1 mL of 1x PBS + 0.04% BSA. The resuspended cells were then passed through a 40 μm cell strainer. The cells were then counted and diluted to 1000 cells/μL. Dissociated cells were fixed using the formaldehyde-based Chromium Next GEM Single Cell Fixed RNA Sample Preparation Kit (10X Genomics, 1000414). The samples were processed according to the 10X Genomics protocol. Single cell libraries were prepared using the Human Probe Set (10X Genomics) and sequenced using an Illumina NovaSeq 6000.

### Single-cell RNA sequencing (scRNAseq) processing

Generation of feature-barcode matrices were generated using Cell Ranger 7.0.1 (10X Genomics) with the reference genome GRCh38. Sample feature-barcode matrices were loaded using Seurat v4^125^ as raw data. To perform initial quality control, we removed cells with less than 1000 unique molecular identifiers (UMI). Low quality or dying cells typically exhibit heightened mitochondrial RNA contamination, however, the 10X Genomics Human Probe Set accounts for mitochondrial protein coding genes by adjusting fold coverage for these genes. As a precaution, we removed cells with more than 10% mitochondrial transcripts.

After quality control, we obtained a total of 33,743 genes and 64,531 high-quality cells, including 20,777 cells from Control organoids, 22,421 cells from SCN2A KO organoids and 21,333 cells from SCN2A G1744X organoids. We performed normalization using SCTransform^126^ (version 0.4.1) with mitochondrial read as a variable to regress and default parameters. Principal Component Analysis (PCA) was performed using default parameters. We then applied Harmony integration^127^ to handle batch-to-batch variability using default parameters. UMAP embeddings were calculated using the Harmony reduction and SCT assay.

### Cell type annotation

Cell type annotation was performed through a combined approach of manual and semi-automated annotation. In our manual approach, we use canonical cell type markers of neural cell types from literature. For our automated approach, we used two reference scRNAseq datasets, a cerebral organoid atlas^33^ and a mid-gestation prenatal cortex atlas^29^ and used the package scClassify^41^ to estimate each cell type. Reference data sets were imported from feature-barcode matrices or Seurat objects and preprocessed as per their source reference. Frequency scores outputted by scClassify were used to guide annotation in conjunction with canonical cell type marker expression. To focus on neural cell type and neurodevelopment, mesenchymal-like cells and unknown (unk) classified clusters were removed from downstream analysis.

### Spatial mapping and regional identification

Unbiased spatial mapping was performed using Voxhunt (v1.1.0)^45^. The top 50 most variable features from the ISH Allen Brain Atlas dataset of the E15 mouse brain were used to create similarity maps and plotted in sagittal view. The BrainSpan transcriptomic database was also used with VoxHunt default parameters.

### Differential abundance testing of cell types

To perform differential abundance testing, we used the Milo R package (v1.10.0)^54^. For each comparison (ie. Control vs SCN2A KO and Control vs SCN2A G1744X) of differential abundance across all cell types, we created a Milo object and used the *buildGraph* function with *k* = 30 and *d* = 30. We then used *makeNhoods* and set *prop=*0.05 followed by *calcNhoodDistance*. For plotting, significance for differentially abundant neighbourhoods is reported as spatial FDR less than 0.1.

### Differentially expressed gene analysis and gene ontology analysis

Differentially expressed gene (DEG) analysis was performed by comparing Control and SCN2A KO datasets and Control and SCN2A G1744X data sets within each cell type. We used the *FindMarkers* function with the Wilcoxon test and post-hoc test set to FDR. Genes with FDR-adjusted p-values lower than 0.05 in DE tests were considered significant. Gene Ontology (GO) term enrichment was performed using g:Profiler’s GOSt function (v0.2.2) using the default *g_SCS* correction method and GO:BP database^51^. GO enrichment was performed on each cell type between each comparison. Significant GO terms were defined as having a p-value less than 0.05.

To gain a more comprehensive view of what biological themes may be affected, we employed the use of REVIGO (v1.14.1)^52^ which summarizes GO terms by finding a representative subset of terms using semantic similarity measures. Results were obtained using default parameters and visualized using treemap plots for each cell type.

To understand what synaptic processes may be affected, we used SynGO. DEGs were imported to the SynGO website: https://www.syngoportal.org for GO term analysis and where the resulting sunburst plots were obtained.

### Gene overlap analysis of SCN2A mutant organoid single-cell RNAseq

DEG overlap analysis was performed using DEGs from upper-layer neurons (ExM-U-2) and deep-layer neurons (ExM-D) from either SCN2A KO or SCN2A G1744X genotypes. ASD risk genes list was obtained from Trost et al.^9^. Gene overlap statistical significance was assessed using the GeneOverlap R package (v1.38.0) as previously described^128^ and results were visualized on UpSet plot made from the UpSetR (v1.4.0) package.

### Trajectory inference and pseudotime analysis

Trajectory inference and pseudotime analysis was performed using monocle3 (v1.3.1)^50^. We converted Seurat objects into Monocle3 *cell_data_set* objects. We then learned trajectories on the MAP using the *learn_graph* function and get the pseudotemporal order of cells using the *order_cells* function. We chose radial glia (RG) as the start point of the trajectory. We plotted the pseudotemporal value of each cell using a density plot. The resulting pseudotime distributions for all cells and each cell types between genotypes were compared using the Kolmogorov-Smirnov test.

### Cell-cell communication analysis

Cell communication analysis was conducted using the CellChat^79^ package. Processed scRNAseq data were loaded into Cell-Chat. The ligand-receptor pairs in the human version of the database CellChatDB was used for calculating cell interactions. The downstream analyses and generation of plots were performed as described in Jin et al.^76^ The default “trimean” threshold was used for more stringent identification of interactions and the default minimum cell number per group of 10 was used.

### Viral labelling of inhibitory neurons in organoids

Organoids were transduced either at Day 90 or Day 106 with AAV-mDLX-GFP (Addgene, 83900-AAV9) at 1:1000 stock virus into 500 μL of conditioned cerebral maturation media in a 24-well ultra-low attachment plate in individual wells. The next day, 1 mL of fresh maturation media was added to each well. Three days post-transduction, organoids were transferred into a 6-well ultra-low attachment plate in maturation media while spinning on an orbital shaker. Organoids were matured for another 2 weeks post-transduction to allow high expression of GFP. After, organoids were processed for immunostaining using 50 μm thick sections.

### Calcium imaging

Organoids were transduced with AAV-hSyn-GCaMP6f (Addgene, 100837-AAV1) or AAV-mDLX-GCaMP6f (Addgene, 83899-AAV9) at 1:1000 or 1:500 stock virus into 500 μL of conditioned cerebral maturation media in a 24-well ultra-low attachment plate in individual wells. The next day, 1 mL of fresh maturation media was added to each well. Three days post-transduction, organoids were transferred into a 6-well ultra-low attachment plate in maturation media while spinning on an orbital shaker. After 7 days post-transduction, the media was switched to BrainPhys (STEMCELL Technologies, 08605) supplemented with 1 μg/μL laminin. The organoids were cultured in BrainPhys for at least 2 weeks prior to imaging.

One day prior to imaging, organoids were transferred to a 1.5 coverslip 35 mm dish (MATTEK, P35G-1.5-14-C) in fresh BrainPhys. Imaging was performed using an LSM880 confocal microscope with Airyscan (Zeiss) with a heating chamber and controller (Live Cell Instruments) to maintain physiological conditions (37°C, 5% CO2). A Plan-Apochromat 10x/0.45 NA objective was used with 2X digital zoom to obtain a 712 x 803 frame size with bi-directional scanning at 5.2 frames per second. Time lapse videos of spontaneous calcium activity was recorded for 5 minutes.

For calcium imaging of DLX-GCaMP6f organoids, a Leica STELLARIS confocal microscope was used with a heating chamber to maintain physiological conditions (37°C, 5% CO2). A 10x objective was used with 2X digital zoom at a resolution of 512 x 512 with a line averaging of 2 and resonance scanning at 7.55 frames per second. Spontaneous calcium activity was recorded for 5 minutes.

### Calcium imaging analysis

Analysis was performed using a stimulation-free MATLAB (MathWorks) protocol as previously described^129,130^. In brief, .czi files obtained from live imaging were converted to .tif format and loaded into scripts in MATLAB R2023a. Regions of interest (ROIs) were placed on neuron somas to calculate the raw GCaMP6f intensity over time. For transient detection, the raw trace intensity was normalized to baseline fluorescence (Δ*F*/*F*_0_). For single-neuron dynamics: 1) amplitude was calculated from normalized GCaMP6f intensity for all detected transients in each trace per minute of recording (average Δ*F*/*F*_0_ of detected transients per neuron) and 2) frequency was calculated as the number of detected transients in each trace per recording minute. For network activity, synchronous firing rate was calculated as the number of detected synchronous transients from all ROIs in one FOV per recording minute.

### Slice preparation

Acute slices for electrophysiology experiments were obtained from 2-month-old organoids. Organoids were placed in ice-cold cutting solution, consisting of the following (in mM) 105 Choline-Cl, 2.5 KCl, 1.25 NaH2PO4, 25 NaHCO3, 25 glucose, 11 Na-ascorbate, 3 Na-pyruvate, 0.5 CaCl2·2H2O, and 7 MgCl2.6H2O. Titrate pH to 7.3–7.4 and osmolarity of 295–300 mOsmol/L and were embedded in 2-3% low-melting gel agarose (A9045, Sigma) prepared in PBS. The embedded samples were then mounted onto a slicing chamber of a vibratome (Leica VT 1200) using glue. The chamber was filled with slicing solution, and slices were cut to a thickness of 150-200 µm. The slices were transferred to an artificial cerebrospinal fluid (aCSF) at 32-34 °C for a minimum of 30 minutes, containing the following (in mM): 124 NaCl, 2.5 KCl, 1.25 NaH2PO4, 1 MgCl2.6H2O, 2 CaCl2.2H2O, 25 NaHCO3, and 25 glucose, 1 Na-ascorbate, 1 Na-pyruvate with pH 7.3–7.4 and osmolarity of 295–300 mOsmol/L. All the solutions were saturated with 95% O2 and 5% CO2.

### Whole-cell patch clamp recordings

During recordings, individual slices were placed in a submersion chamber and perfused with aCSF at a rate of 1.5-2 ml/min in room temperature. Organoid slices were observed using an upright microscope (Olympus U-TV1XC) equipped with a water-immersion long-working distance objective (40x; differential interference contrast, DIC) and an infrared CCD camera (DAGE-MTI IR-2000). Data was collected using a Multiclamp 700B amplifier (Molecular Devices) and digitized at 10 kHz using Digidata 1550B and pClamp 11.1 software (Molecular Devices). The recordings were low pass filtered at 1 kHz. The access resistance was continuously monitored throughout the experiment, and data were included only if the holding current and series resistance remained stable (with less than a 25% change in the initial series resistance). Neuron demonstrating pyramidal-like morphology were selected for the experiments.

Action potential-like depolarizations were defined as events exhibiting a regenerative upstroke (maximum dV/dt ≥ 10 mV/ms) and a distinct peak, with at least two of the following criteria: amplitude ≥ 20 mV (from threshold to peak) and/or overshoot ≥ −10 mV. Events not meeting these criteria were excluded from analysis. In addition, all events classified as action potentials were required to exhibit a distinct peak, defined as a clear local maximum with a reversal of dV/dt from positive to negative, ensuring separation from passive depolarizing responses.

To characterize intrinsic electrical properties of neurons, borosilicate glass pipettes (5–8 MΩ) were filled with intracellular solution consisting of following (in mM): 110 K-gluconate, 10 KCl, 0.2 EGTA, 10 HEPES, 4 MgATP, 0.3 NaGTP, and 10 phosphocreatine (pH 7.3 with KOH, 290–295 mOsmol/L). Whole cell patch clamp recording was performed and resting membrane voltage was recorded in the current clamp configuration. A small bias current was applied, when necessary, to maintain the baseline membrane voltage between −55 and −60 mV, facilitating action potential firing. Single action potential properties were extracted using Clampfit 11.1.

Phase-plot analysis was performed to assess the dynamics of action potential initiation and upstroke, using Clampfit. The resulting dV/dt values were plotted against the corresponding membrane voltage (V) to generate phase plots. The maximum dV/dt slope was defined as the peak values of the dV/dt trace during the rising phase of the first action potential evoked at rheobase. The maximum decay slope was defined as peak negative dv/dt value during the repolarization phase.

To elicit multiple action potential firing, a series of brief depolarizing current pulses were delivered in current clamp. Square current injections of 20 ms duration were applied in 10 pA increments, starting at 10 pA. For each current amplitude, 10 consecutive pulses were delivered, with a 1-second inter-stimulus interval to allow the membrane potential to recover between stimuli.

Na^+^ currents were recorded using an internal solution containing (in mM): 120 Cs-methanesulfonate, 5 CsCl, 10 HEPES, 0.5 EGTA, 2 MgCl_2_, 4 MgATP, 0.3 NaGTP, and 10 phosphocreatine (pH 7.3 with CsOH and 290–295 mOsmol/L). 100 µM CdCl_2_ (Sigma, 202908), and 100 µM 4-Aminopyridine (504-24-5, hellobio) were added to the aCSF to block voltage-gated Ca^2+^ and K^+^ channels, respectively.

For activation properties, a voltage step protocol was applied from a holding potential of −80 mV. Cells were stepped to test potentials ranging from −90 mV to + 40 mV in 5 mV increments for 10 ms. Na^+^ current density was calculated by dividing the peak inward Na^+^ current by the whole cell capacitance, yielding values in pA/pF. For steady-state inactivation, a two-step protocol was used. Cells were held at −80 mV, then subjected to a 100 ms pre-pulse ranging from −90 mV to 0 mV, followed by a test pulse −20 mV for 10 ms to assess availability of Na^+^ channels. Peak currents during the test pulse were normalized to maximum and plotted as a function of the pre-pulse voltage. The activation and inactivation curves were fitted using the Boltzmann equation. All recordings were leak-subtracted using a P/4 protocol.

In a subset of Na^+^ current recordings, 5 nM TTX (Tocris, 1069) was added to the aCSF, and the Na^+^ current activation protocol was tested in control organoids.

### Cyclopamine treatment

To inhibit SHH signaling during organoid development, organoids were treated with 5 μM Cyclopamine (Cyclopamine, V.californicum, Millipore) as previously described^131^ with some modifications. Organoids were treated from day 60 to 90 continuously with each media change. On day 90, organoids were transduced with AAV-mDLX-GFP as described above and allowed to mature for 2 weeks then processed for immunostaining with 50 μm sections.

### Sonic hedgehog reporter assay

HEK293T cells were cultured on glass coverslips at a density of 250k cells/well. One day after plating cells were transduced with a GLI-TAG-Puro lentivirus (LipoExoGen, SKU LTV-0012-1S) at a 1:500 dilution and left to recover for 24 hours. The next day, conditioned media from mutant or drug-treated organoids or 1 μM purmorphamine were added to infected or uninfected cells and incubated for 24 hours. After, cells were fixed with 4% PFA, whole mounted and imaged on a Leica STELLARIS confocal microscope. CellProfiler was used to count cells using brightfield and GFP channels.

### Sonic hedgehog shRNA assay

Custom AAV9-CaMKIIa(short)-shRNA-EGFP were ordered from VectorBuilder. Organoids were transduced at D74 as described above. Expression was evaluated by EGFP expression at 21 and 29 days post-transduction. After 29 days, conditioned media from infected organoids were used for the SHH reporter assay and organoid were processed for immunostaining. The sequence information for shRNAs are: Scramble (CCCTAAGGTTAAGTCGCCCTCGTAGTGAAGCCACAGATGTACGAGGGCGACTTAACCTTAGGT, Vector ID: VB250806-1526scd), shSHH1 (CCTACGAGTCCAAGGCACATATTAGTGAAGCCACAGATGTAATATGTGCCTTGGACTCGTAGT, Vector ID: VB250806-1529eyf), shSHH2 (ACTTTAGCCTACAAGCAGTTTATAGTGAAGCCACAGATGTATAAACTGCTTGTAGGCTAAAGG, Vector ID: VB250925-1710nty).

### Statistics and mixed model analysis

All statistical analyses were performed using RStudio and R version 4.3.0. No statistical methods were used to pre-determine sample sizes. For all experiments, we used a nested (hierarchical) mixed model analysis using the *lme4* (v1.1-35.1) and *lmerTest* (v3.1-3) packages and depending on the normality of the data, checked using a Shapiro-Wilk test (p<0.05 considered not normal), a linear mixed model was used and if the data were not normal, a generalized linear mixed model was used when comparing multiple groups, experimental conditions (ie. Genotype, treatment) were used as fixed effects. In experiments with multiple batches, a random effect for batches of differentiation was used and random effects for organoid and field of view were included for single batch experiments. For experiments with more than two groups, the fixed effect (Genotype or treatment) was assessed by a Type III Wald chi-square test (analysis of deviance), and pairwise comparisons were performed on the estimated marginal means with Tukey’s adjustment. For comparisons between only two conditions, the p-value was taken directly from the model. Distributions were tested using the Kolmogorov-Smirnov test. Significance was assessed by p < 0.05. All error bars represent mean ± S.E.M. unless otherwise stated. A Student’s t-test was performed to compare SCN2A mass spectrometry levels. All statistical tests, sample sizes, fixed and random effects are described in the figure legend results and available in the Source Data file.

## Data availability

scRNAseq data and mass spectrometry data from cerebral organoids are available through GEO <u>GSE292644</u> and PRIDE <u>PXD064388</u>, respectively. Two month-old cerebral organoid scRNAseq data used for reference mapping is available from Mendeley Data (https://data.mendeley.com/datasets/z4jyxnx3vp/3). Second trimester prenatal cortex data is available from the CoDex viewer (http://solo.bmap.ucla.edu/shiny/webapp/). All raw data quantified and statistical analyses for each figure generated in this study are provided in the Source Data file. Source data are provided with this paper.

## Code availability

This paper does not report original code, however, primary software packages used for analysis are listed in the reporting summary. Any additional information required to reanalyze the data reported in this paper is available from the lead contact upon reasonable request.

## Acknowledgements

We thank members of the Singh and Bains labs for reading the manuscript. ImageTwin was used as a pre-submission quality-control tool to screen figures for potential duplication.

## Funding

K.K.S. was supported by the Canadian Institutes of Health Research (CIHR - 148814, 161487, 183919, 197899), the Natural Sciences and Engineering Research Council of Canada (NSERC RGPIN 2019 06332), the Ontario Brain Institute (OBI), the Donald K. Johnson Eye Institute (DKJEI), the Autism Research Institute (ARI 1028010), and the Krembil Research Institute (KRI). J.A.U. was supported by the Vision Science Research Program (VSRP).

## Author contributions

J. A. U. and K. K. S. conceived and designed the project. J. A. U. and K. K. S. wrote the paper with input from Z. D. and J. S. B. All experiments, data analysis, and figures were done by J. A. U. unless otherwise specified. C. O. B. generated the 19-2-2 *SCN2A^+/G^*^1744^*^X^* iPS cell line. S. G. and L. B. performed and analyzed cell cycle exit and apoptosis immunofluorescence experiments. Y. P. generated and maintained the dorsal forebrain organoids. Z. D. performed all electrophysiology experiments with input from J. S. B. J. L. H. and S.W. S. provided patient access and samples.

## Competing Interests

The authors declare no competing interests.

## Supplemental information

Document S1. Figures S1-S20.

Table S1. scRNAseq: DEGs in *SCN2A^+/G^*^1744^*^X^* organoids, related to Figure 2. DEGs were identified per cell type with the Seurat FindMarkers function using a two-sided Wilcoxon rank-sum test, with multiple-comparison correction by the Benja-mini–Hochberg false discovery rate (FDR); genes with FDR-adjusted p < 0.05 were considered significant.

Table S2. scRNAseq: DEGs in *SCN2A^-/-^* organoids, related to Figure 2. DEGs were identified per cell type with the Seurat FindMarkers function using a two-sided Wilcoxon rank-sum test, with multiple-comparison correction by the Benjamini–Hochberg false discovery rate (FDR); genes with FDR-adjusted p < 0.05 were considered significant.

Table S3. scRNAseq: GO term enrichment of upregulated DEGs in *SCN2A^+/G^*^1744^*^X^* organoids, related to Figure 2. GO term enrichment was performed with g:Profiler (GOSt over-representation analysis, a one-sided hypergeometric test), with multiple-comparison correction by g:Profiler’s g:SCS algorithm; terms with adjusted p < 0.05 were considered significant.

Table S4. scRNAseq: GO term enrichment of downregulated DEGs in *SCN2A^+/G^*^1744^*^X^* organoids, related to Figure 2. GO term enrichment was performed with g:Profiler (GOSt over-representation analysis, a one-sided hypergeometric test), with multiple-comparison correction by g:Profiler’s g:SCS algorithm; terms with adjusted p < 0.05 were considered significant.

Table S5. scRNAseq: GO term enrichment of upregulated DEGs in *SCN2A^-/-^* organoids, related to Figure 2. GO term enrichment was performed with g:Profiler (GOSt over-representation analysis, a one-sided hypergeometric test), with multiple-comparison correction by g:Profiler’s g:SCS algorithm; terms with adjusted p < 0.05 were considered significant.

Table S6. scRNAseq: GO term enrichment of downregulated DEGs in *SCN2A^-/-^* organoids, related to Figure 2. GO term enrichment was performed with g:Profiler (GOSt over-representation analysis, a one-sided hypergeometric test), with multiple-comparison correction by g:Profiler’s g:SCS algorithm; terms with adjusted p < 0.05 were considered significant.

Table S7. Information on gRNAs and off-target sequencing, related to Figure S2.

## Notes

### Competing Interest Statement

The authors have declared no competing interest.

### Summary of Updates

Major revisions to the original manuscript and is now accepted-in-principle at Nature Communications. Figures 3 and 4 were updated to include new pharmacological experiments linking sodium channel activity to neurogenesis. Figure 5 was updated with more mechanistic data linking SCN2A loss-of-function to the SHH pathway. Figure 6 updated with new electrophysiological characterization of SCN2A mutant neurons. Supplemental data have been updated with new supporting data for the above as well as further mechanistic characterization of SCN2A's role in SHH-driven IN neurogenesis. Main text has been updated to reviewer suggestions and figure legends now describe the statistics in compliance to Nature Communications' standards.

