## Supplemental Information for "The sodium channel SCN2A regulates cortical excitatory and inhibitory neurogenesis"

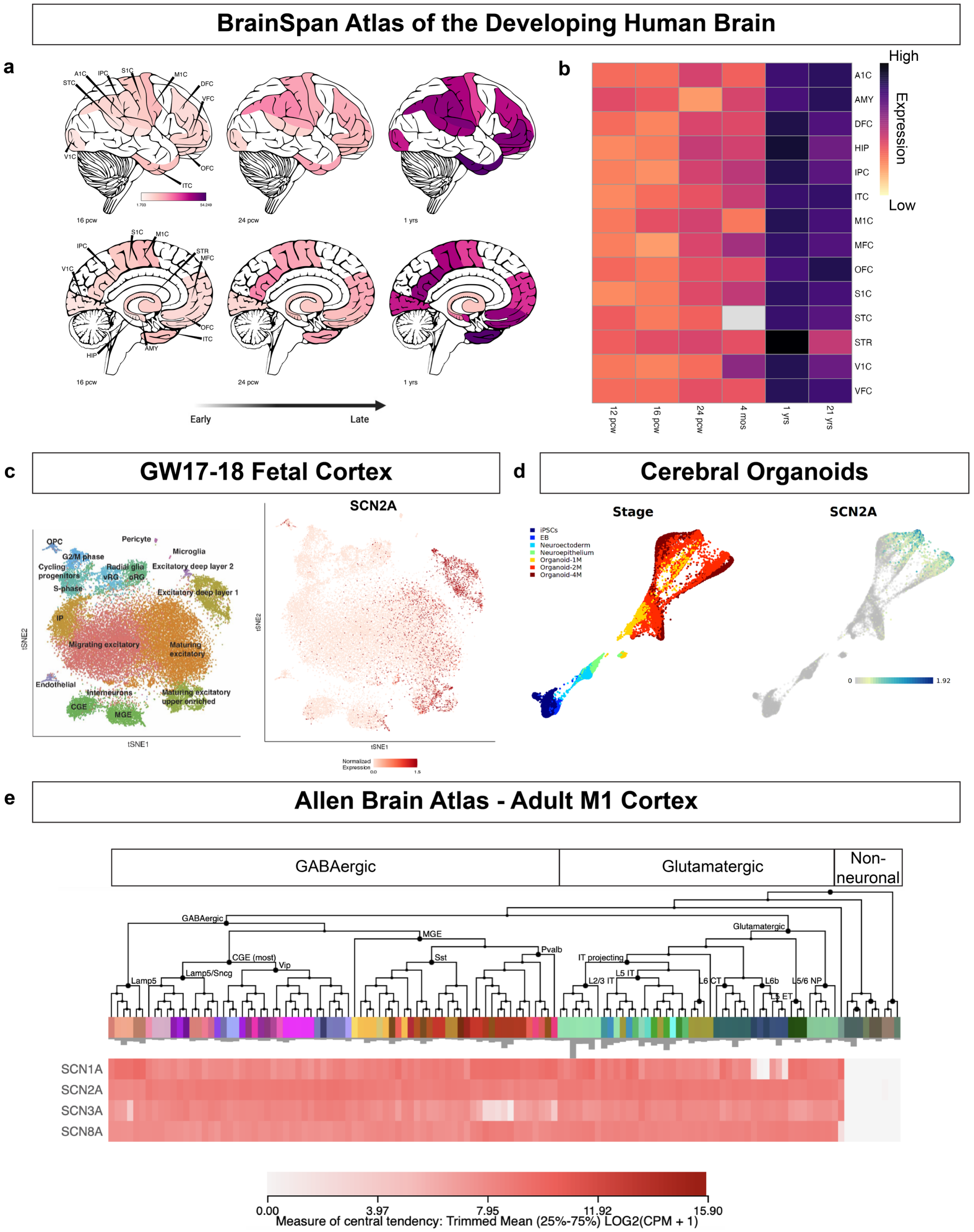


**Fig. S1.** ***SCN2A*** **is expressed in the prenatal and adult brain and in cerebral organoids.**
**a)** Representation of the expression of *SCN2A* in the human brain from the BrainSpan transcriptomic dataset (published in Miller et al.[28]).
**b)** Heatmap of the expression of *SCN2A* across early development in different regions.
**c)** Expression of *SCN2A* in mid-gestation prenatal cortex single-cell RNAseq data (published in Polioudakis et al.[29]).
**d)** Expression of *SCN2A* in cerebral organoid single-cell RNAseq data (published in Kanton et al.[33]).
**e)** *SCN2A* and other CNS voltage-gated sodium channels expression in adult human cortex (from the Allen Brain Atlas, published in Hodge et al.[34]).

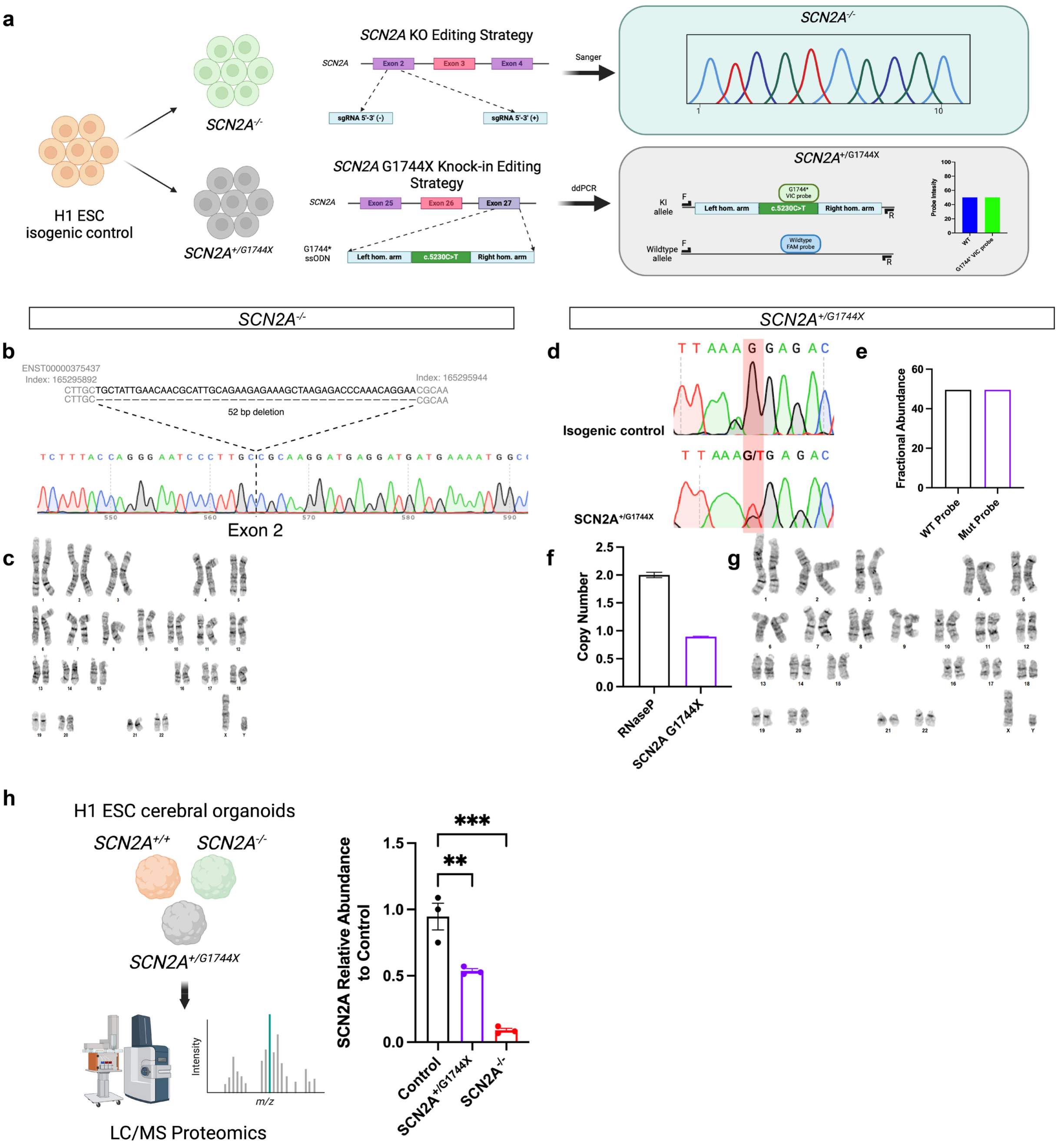


**Fig. S2. Generation and validation of H1 isogenic embryonic stem cell lines.**

**a)** Schematic showing CRISPR editing strategy to generate either a homozygous *SCN2A* knock-out (Exon 2) or heterozygous knock-in of an ASD patient-specific mutation G1744X (Exon 27). Validation was performed using sanger sequencing (*SCN2A*$^{-/-}$) or ddPCR (*SCN2A*$^{+/G1744X}$).

**b)** Sanger sequencing of homozygous KO of *SCN2A*.

**c)** Karyotype analysis for the *SCN2A*$^{-/-}$ line used in this study.

**d)** Sanger sequencing of isogenic control and *SCN2A*$^{+/G1744X}$ cells. Highlighted in red is the heterozygous detection of the knock-in allele.

**e)** ddPCR analysis of *SCN2A*$^{+/G1744X}$ cells showing equal detection of wildtype and knock-in alleles.

**f )** ddPCR analysis of *SCN2A*$^{+/G1744X}$ cells showing 1 copy of the knock-in allele and 2 copies of an autosomal gene RNaseP.

**g)** Karyotype analysis for the *SCN2A*$^{+/G1744X}$ line used in this study.

**h)** Schematic showing mass spectrometry analysis of Day 90 cerebral organoids and quantification of SCN2A protein detected (N = 1 batch, 3 organoids per genotype). Data represent mean ± SEM. Two-tailed Student's t-test. Created in BioRender. Singh, K. (2026) https://BioRender.com/0otxclw.

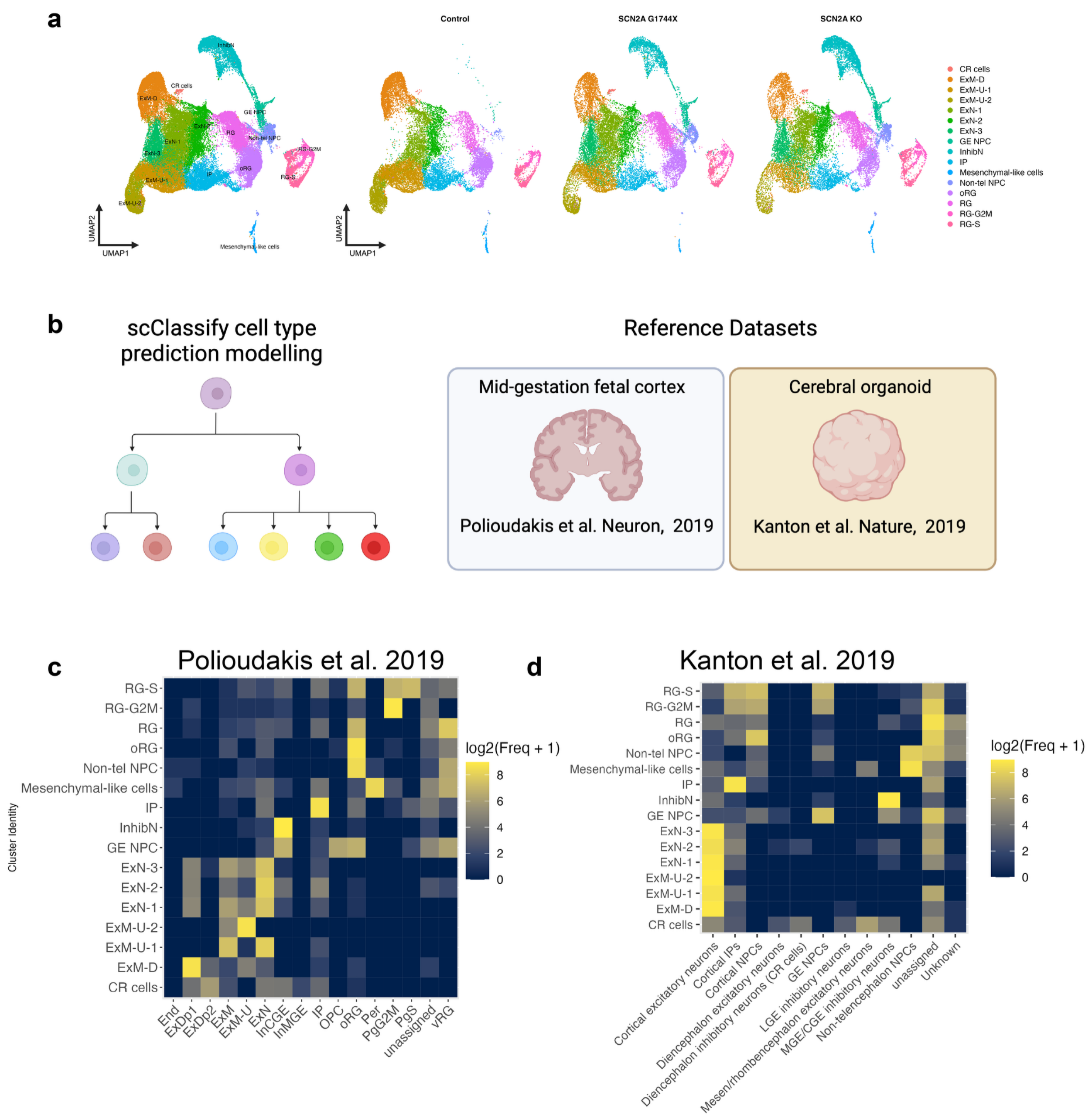


**Fig. S3. Cell type annotation and reference mapping in cerebral organoids**
**a)** UMAP plots of Day 90 cerebral organoids by genotype.
**b)** Schematic of reference mapping single-cell RNAseq data using scClassify.
**c)** Heatmap of cell type predictions using mid-gestation prenatal cortex data from Polioudakis et al.29.
**d)** Heatmap of cell type predictions using cerebral organoid data from Kanton et al.[33].
Created in BioRender. Singh, K. (2026) https://BioRender.com/0otxclw.

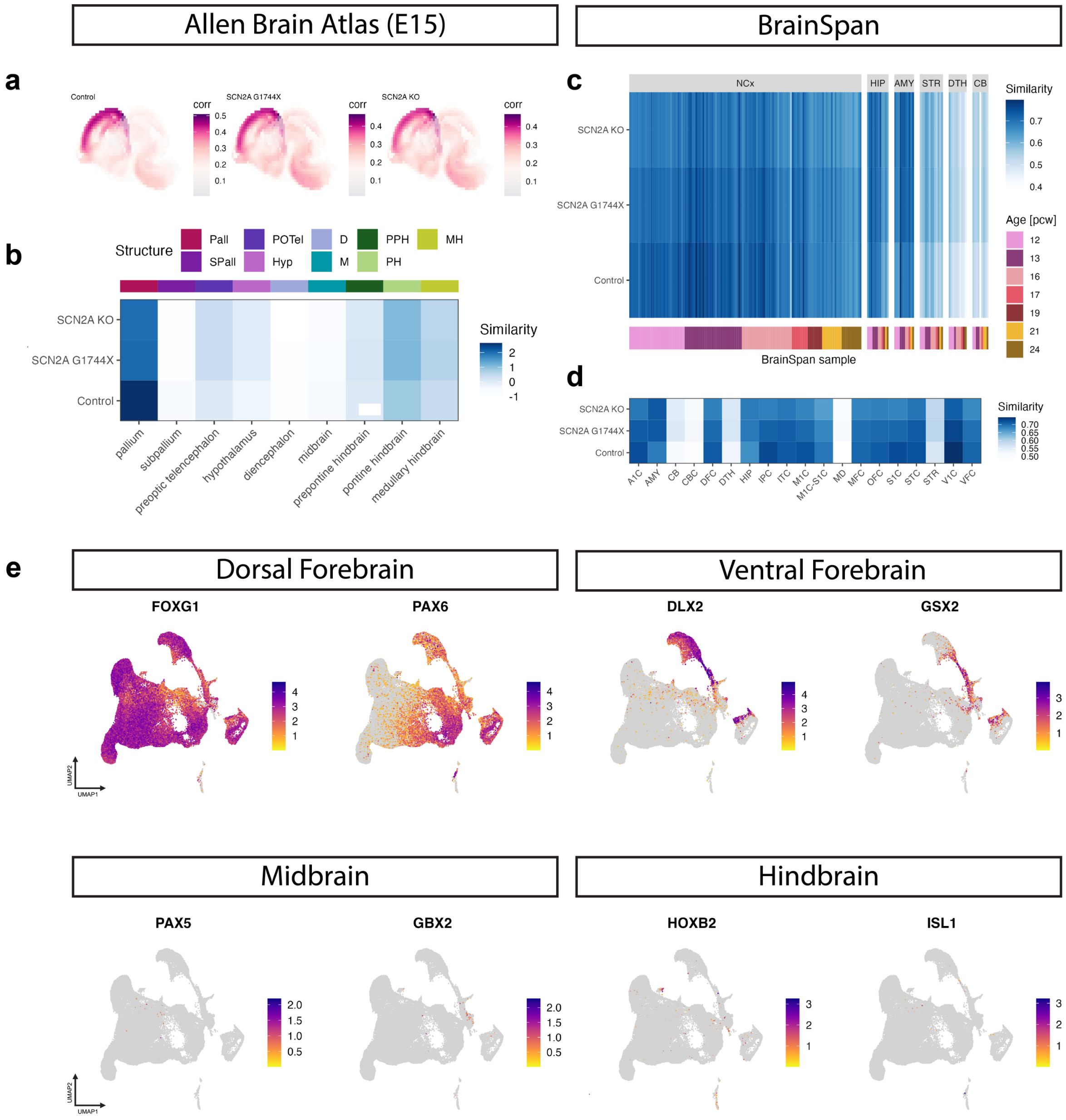


**Fig. S4. Additional annotation and regional identity mapping**

**a,b)** VoxHunt spatial brain mapping of Day 90 cerebral organoids onto the Allen Brain Institute E15.5 mouse brain data separated by genotype.

**c, d)** VoxHunt prediction of brain regional identity of cerebral organoids based on mid-gestation human brain.

**e)** Feature plots showing expression of selected regional markers from cerebral organoids.

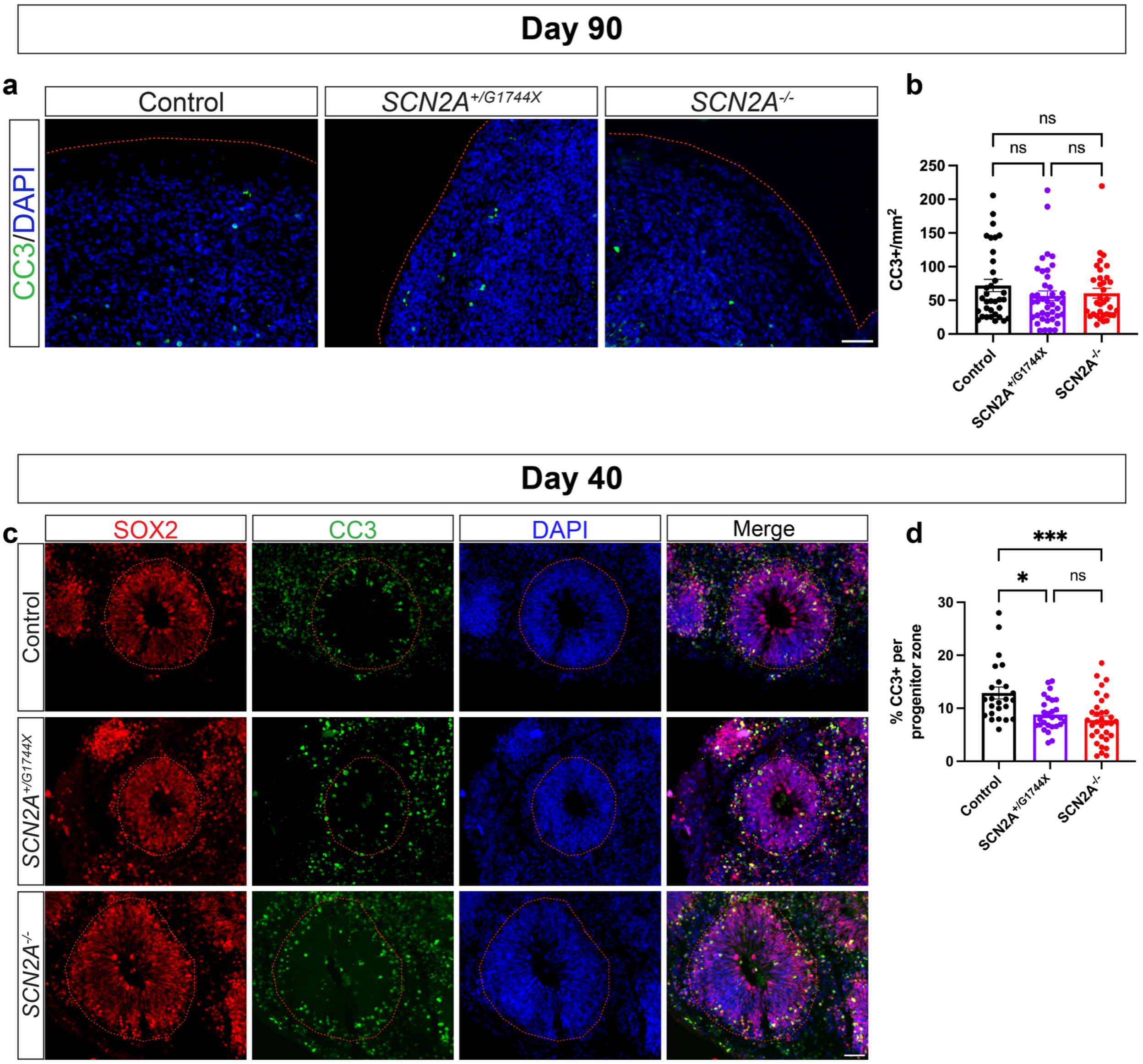


**Fig. S5. Loss of *SCN2A* does not alter cell viability in mature organoids**

**a)** Immunostaining for apoptotic marker Cleaved Caspase-3 (CC3) in Day 90 organoids. Scale bar = 100 µm.

**b)** Quantification of density of CC3$^{+}$ cells in Day 90 organoids (N = 3 independent batches, Control n = 34 FOVs from 12 organoids, *SCN2A*$^{+/G1744X}$ n = 41 FOVs from 12 organoids, *SCN2A*$^{-/-}$ n = 34 FOVs from 12 organoids, mixed-effects models with Genotype as a fixed effect and batch as a random effect).

**c)** Immunostaining for CC3 in SOX2$^{+}$ progenitor zones. Scale bar = 50 µm.

**d)** Quantification of the percentage of CC3$^{+}$ cells in progenitor zones of Day 40 organoids (N = 3 independent batches, Control n = 24 progenitor zones from 12 organoids, *SCN2A*$^{+/G1744X}$ n = 27 progenitor zones from 9 organoids, *SCN2A*$^{-/-}$ n = 33 progenitor zones from 10 organoids, mixed-effects models with Genotype as a fixed effect and batch as a random effect).

All data are reported as mean ± SEM. Mixed-effects models with Type III Wald $\chi^2$ test for fixed effects, post-hoc Tukey test. *$p < 0.05$, **$p < 0.01$, ***$p < 0.001$, ****$p < 0.0001$; n.s., not significant.

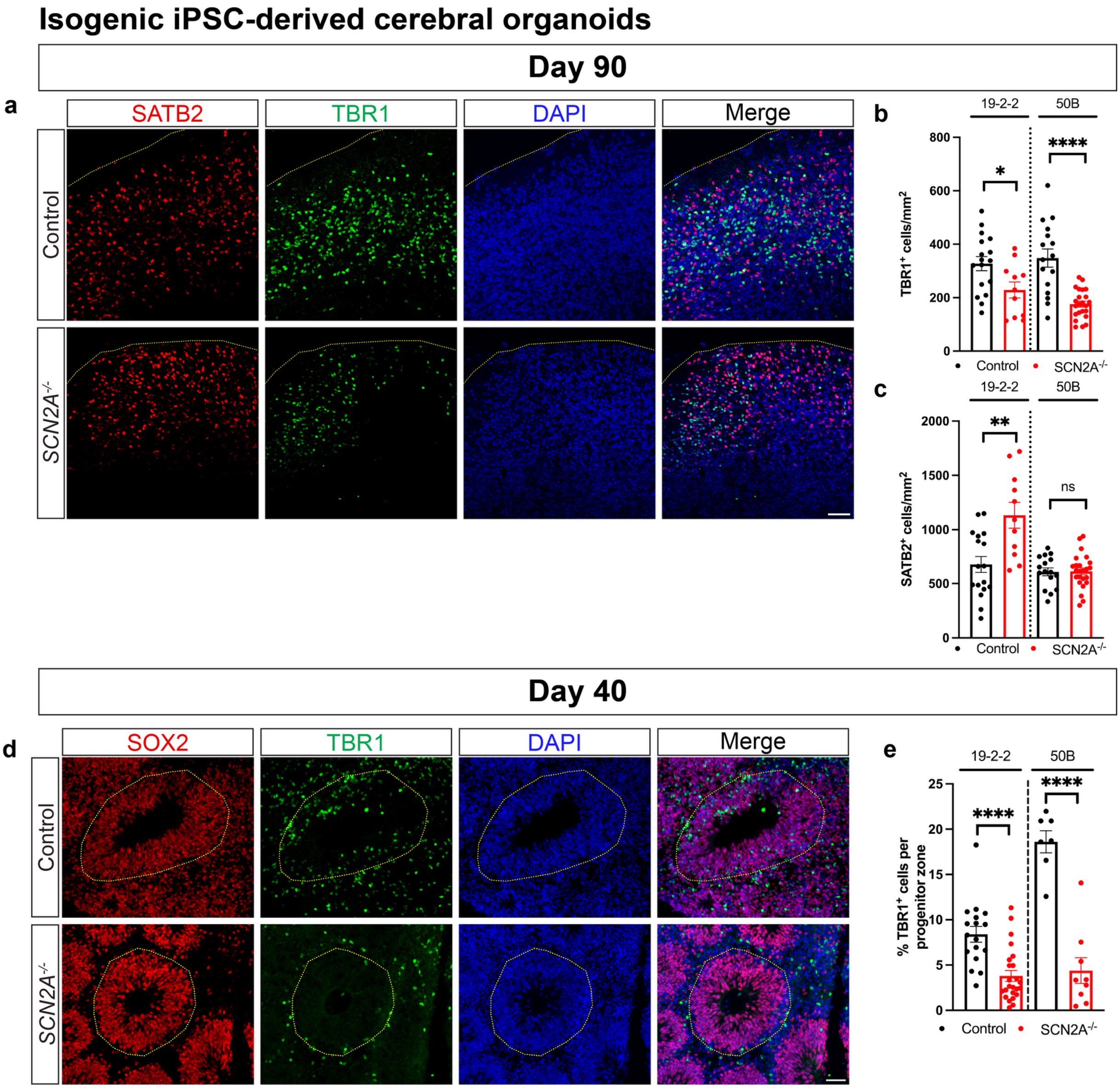


**Fig. S6. Loss of *SCN2A* impairs excitatory neuron production across development in isogenic iPSC-derived cerebral organoids**

**a)** Immunostaining for the upper layer neuron marker SATB2 and deep layer neuron marker TBR1 in Day 90 isogenic iPSC-derived organoids derived from the 19-2-2 and 50B cell lines. Scale bar = 100 µm.

**b, c)** Quantification of density of TBR1$^+$ or SATB2$^+$ neurons in Day 90 organoids by cell line (19-2-2: N = 1 batch, Control n = 17 FOVs from 4 organoids, *SCN2A*$^{-/-}$ n = 11 FOVs from 4 organoids; 50B: N = 1 batch, Control n = 16 FOVs from 4 organoids, *SCN2A*$^{-/-}$ n = 24, mixed-effects models with Genotype as a fixed effect and batch as a random effect).

**d)** Immunostaining for the deep layer neuron marker TBR1 and neural progenitor marker SOX2 in Day 40 organoids derived from the 19-2-2 and 50B cell lines. Scale bar = 50 µm.

**e)** Quantification of the percentage of TBR1$^+$ neurons in SOX2$^+$ progenitor zones by cell line (19-2-2: N = 1 batch, Control n = 17 progenitor zones from 4 organoids, *SCN2A*$^{-/-}$ n = 24 progenitor zones from 4 organoids; 50B: N = 1 batch, Control n = 7 progenitor zones from 4 organoids, *SCN2A*$^{-/-}$ n = 9 progenitor zones from 4 organoids, mixed-effects models with Genotype as a fixed effect and batch as a random effect).

All data are reported as mean ± SEM. Mixed-effects models. *p < 0.05, **p < 0.01, ***p < 0.001, ****p < 0.0001; n.s., not significant.

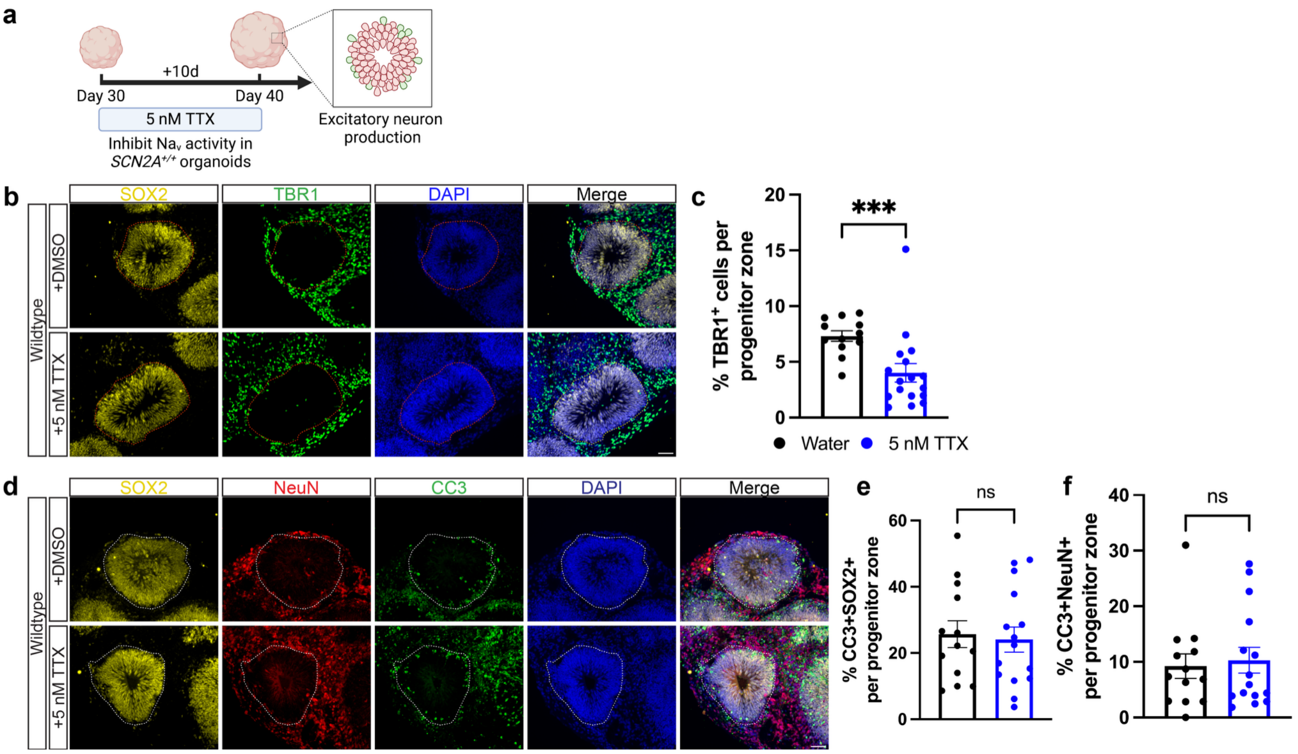


**Fig. S7. Voltage-gated sodium channel-dependent neurogenesis in cerebral organoids**
**a)** Schematic of pharmacological inhibition of VGSCs in Day 40 Control organoids using 5 nM of TTX.
**b)** Immunostaining for TBR1 and SOX2 in Day 40 organoids. Scale bar = 50 µm.
**c)** Quantification of the percentage of $TBR1^+$ neurons in $SOX2^+$ progenitor zones in TTX-treated organoids (N = 1 batch, Water n = 12 progenitor zones from 3 organoids, TTX n = 17 progenitor zones from 4 organoids, mixed-effects models with condition as the fixed effect and organoid as a random effect).
**d)** Immunostaining for NeuN, SOX2 and CC3 in Day 40 organoids. Scale bar = 50 µm.
**e, f)** Quantification of the percentage of CC3+SOX2+ (e) or CC3+NeuN+ (f) per progenitor zone in TTX-treated organoids (N = 1 batch, Water n = 13 progenitor zones from 2 organoids, TTX n = 15 progenitor zones from 3 organoids, mixed-effects models with condition as the fixed effect and organoid as a random effect).
All data are reported as mean ± SEM. Mixed-effects models. *$p < 0.05$, **$p < 0.01$, ***$p < 0.001$, ****$p < 0.0001$; n.s., not significant.
Created in BioRender. Singh, K. (2026) https://BioRender.com/0otxclw.

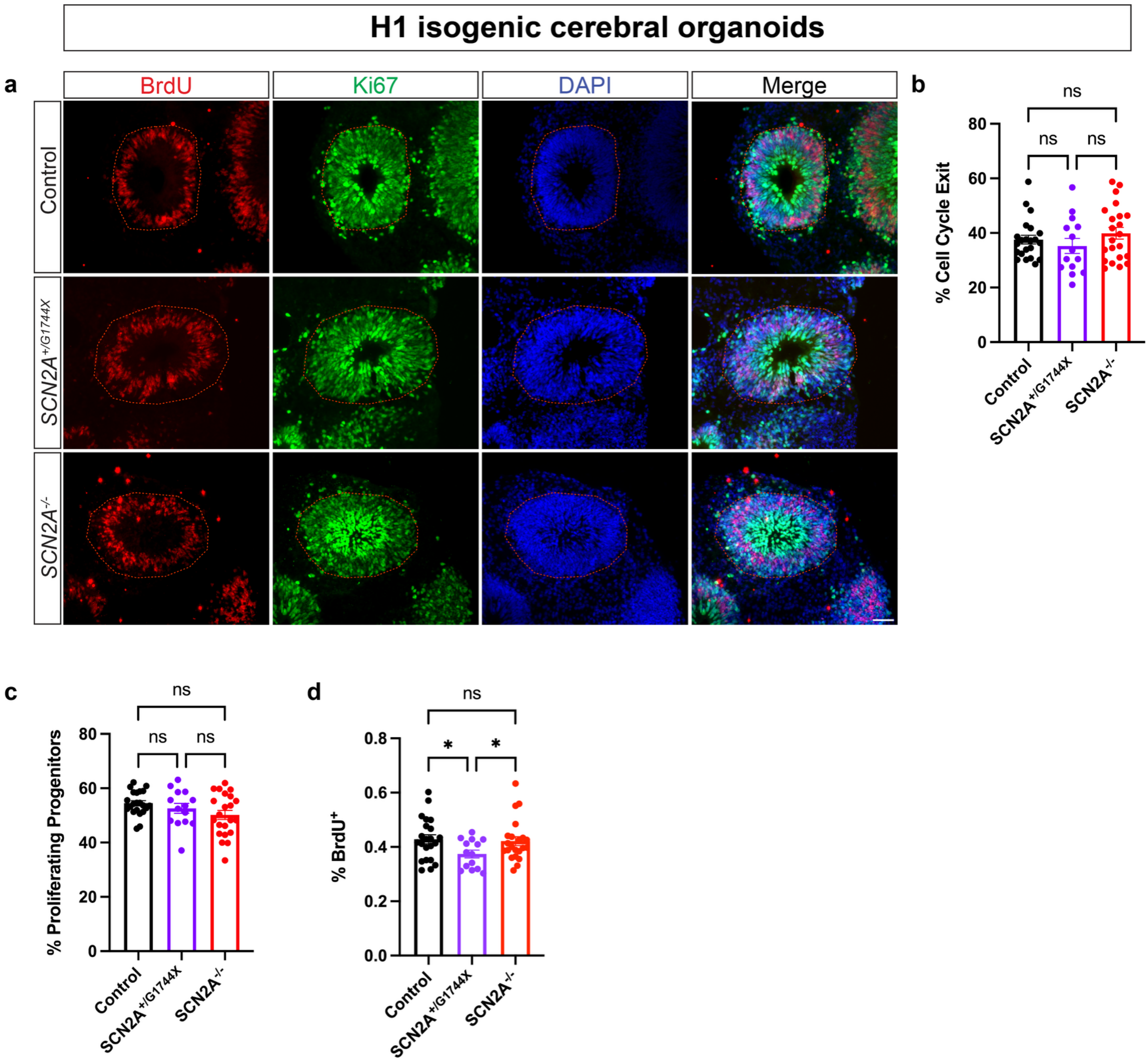


**Fig. S8.** ***SCN2A*** **does not regulate neural progenitor proliferation or cell cycle exit,**
**a)** Immunostaining for BrdU, Ki67 and SOX2 in Day 40 H1-ESC organoids. Scale bar = 50 µm.
**b-d)** Quantification of cell cycle exit ($BrdU^+Ki67^-$/total $BrdU^+$), percentage of proliferating progenitors ($Ki67^+SOX2^+$) and percentage of $BrdU^+$ cells (N = 1 batch, Control n = 22 progenitor zones from 4 organoids, *SCN2A*$^{+/G1744X}$ n = 14 progenitor zones from 4 organoids, *SCN2A*$^{-/-}$ n = 22 progenitor zones from 4 organoids, mixed-effects models with Genotype as a fixed effect and each progenitor zone nested per genotype).
All data are reported as mean ± SEM. Mixed-effects models with Type III Wald $\chi^2$ test for fixed effects, post-hoc Tukey test. *p < 0.05, **p < 0.01, ***p < 0.001, ****p < 0.0001; n.s., not significant.

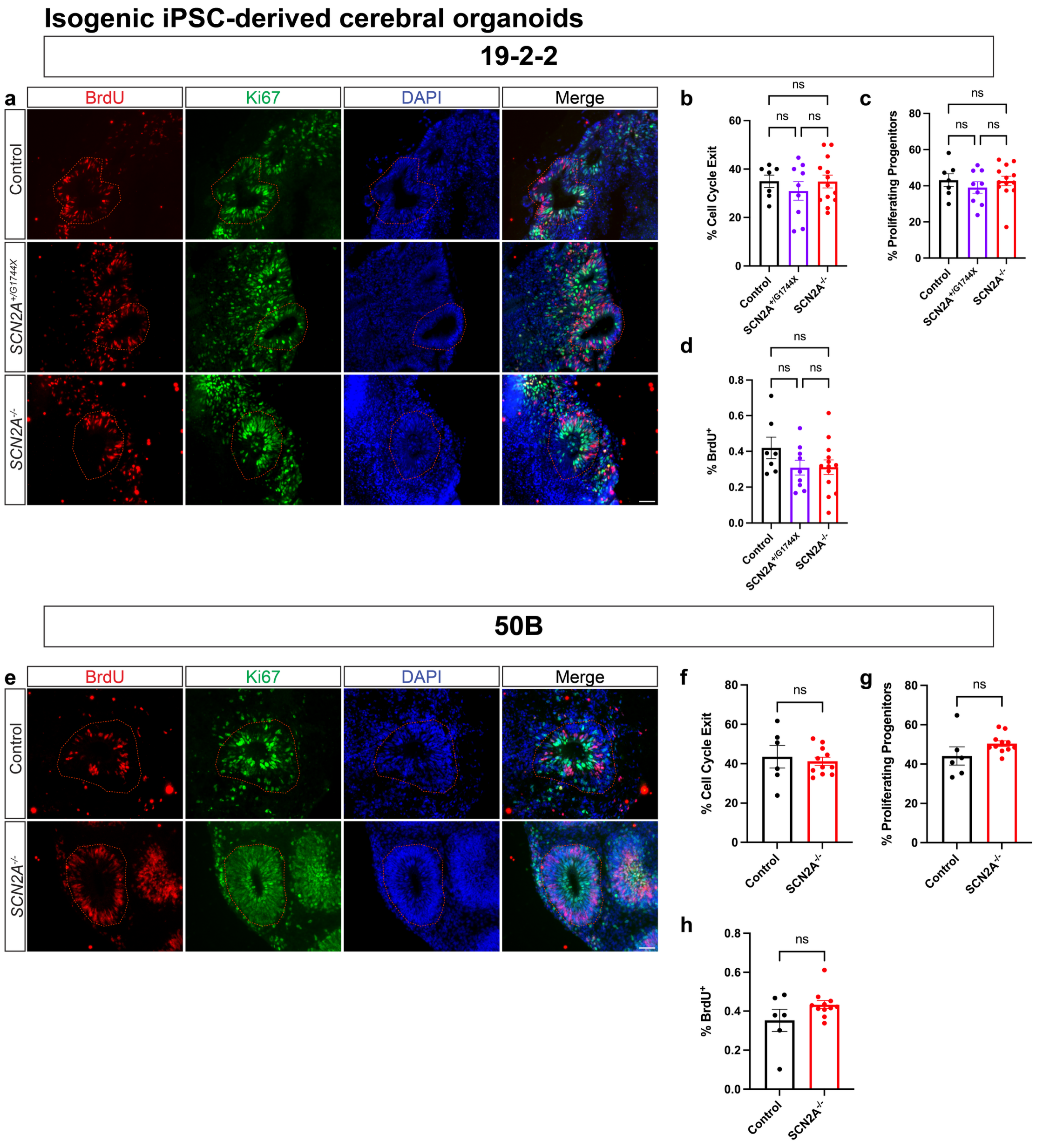


**Fig. S9. *SCN2A* does not regulate neural progenitor proliferation or cell cycle exit in isogenic iPSC-derived cerebral organoids**

**a)** Immunostaining for BrdU and Ki67 in Day 40 organoids. Scale bar = 50 µm.

**b-d)** Quantification of cell cycle exit, percentage of $BrdU^+$ cells and percentage of proliferating progenitors (N = 1 batch, Control n = 7 progenitor zones from 4 organoids, $SCN2A^{+/G1744X}$ n = 9 progenitor zones from 4 organoids, $SCN2A^{-/-}$ n = 13 progenitor zones from 4 organoids, mixed-effects models with Genotype as a fixed effect and each progenitor zone nested per genotype). Mixed-effects models with Type III Wald $\chi^2$ test for fixed effects, post-hoc Tukey test.

**e)** Immunostaining for BrdU and Ki67 in Day 40 organoids. Scale bar = 50 µm.

**f-h)** Quantification of cell cycle exit, percentage of $BrdU^+$ cells and percentage of proliferating progenitors (N = 1 batch, Control n = 6 progenitor zones from 4 organoids, $SCN2A^{-/-}$ n = 11 progenitor zones from 4 organoids, mixed-effects models with Genotype as a fixed effect and each progenitor zone nested per genotype).

All data are reported as mean ± SEM. *p < 0.05, **p < 0.01, ***p < 0.001, ****p < 0.0001; n.s., not significant.

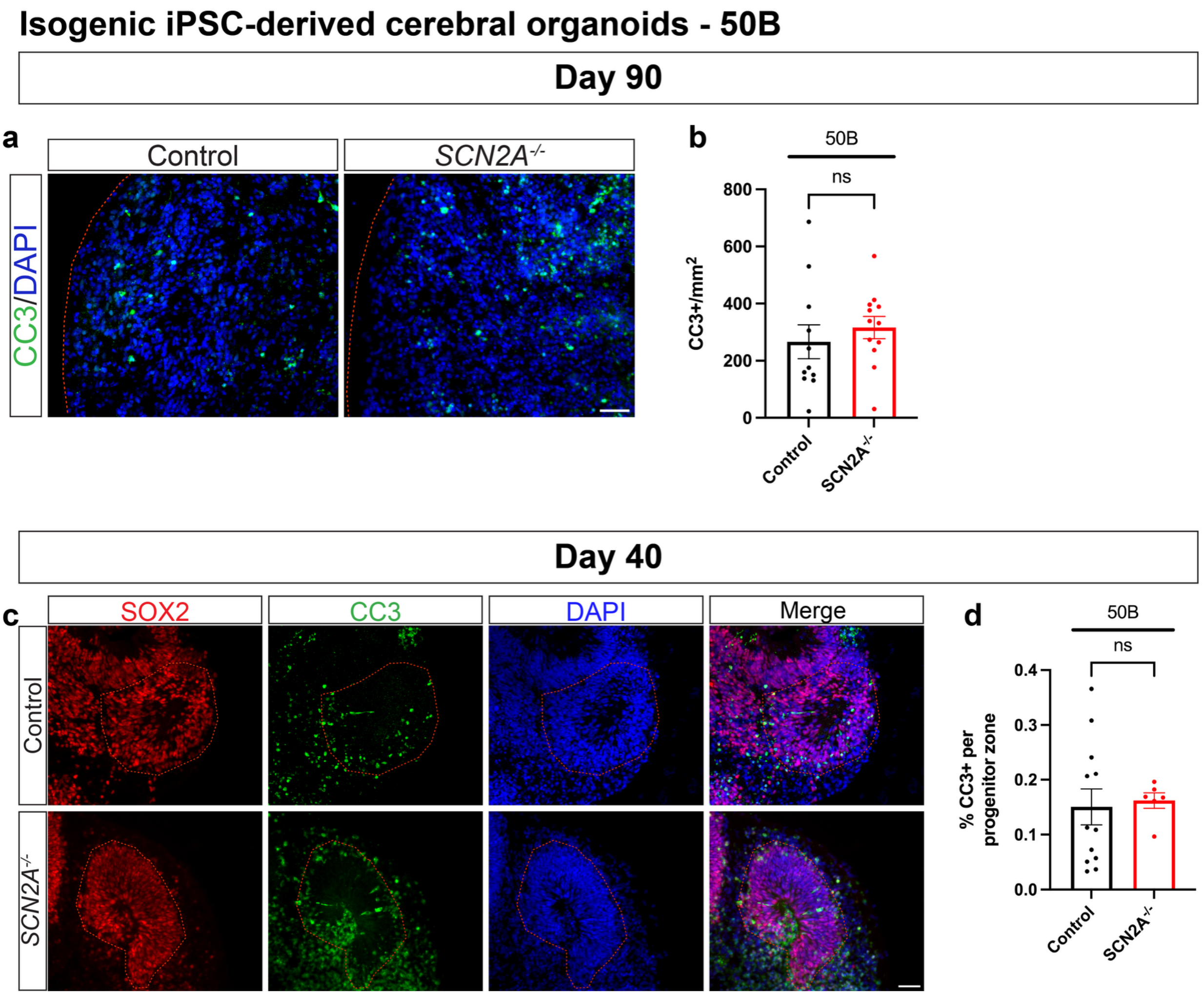


**Fig. S10. Loss of *SCN2A* does not alter cell viability in mature isogenic iPSC-derived cerebral organoids**
**a)** Immunostaining for apoptotic marker Cleaved Caspase-3 (CC3) in Day 90 organoids. Scale bar = 100 µm.
**b)** Quantification of density of CC3$^+$ cells in Day 90 organoids (N = 3 independent batches, Control n = 11 FOVs from 3 organoids, *SCN2A*$^{-/-}$ n = 12 FOVs from 3 organoids, mixed-effects models with Genotype as a fixed effect and each FOV nested per genotype).
**c)** Immunostaining for CC3 in SOX2$^+$ progenitor zones. Scale bar = 50 µm.
**d)** Quantification of the percentage of CC3$^+$ cells in progenitor zones of Day 40 organoids (N = 3 independent batches, Control n = 12 progenitor zones from 3 organoids, *SCN2A*$^{-/-}$ n = 6 progenitor zones from 3 organoids, mixed-effects models with Genotype as a fixed effect and each progenitor zone nested per genotype).
All data are reported as mean ± SEM. Mixed-effects models. *$p < 0.05$, **$p < 0.01$, ***$p < 0.001$, ****$p < 0.0001$; n.s., not significant.

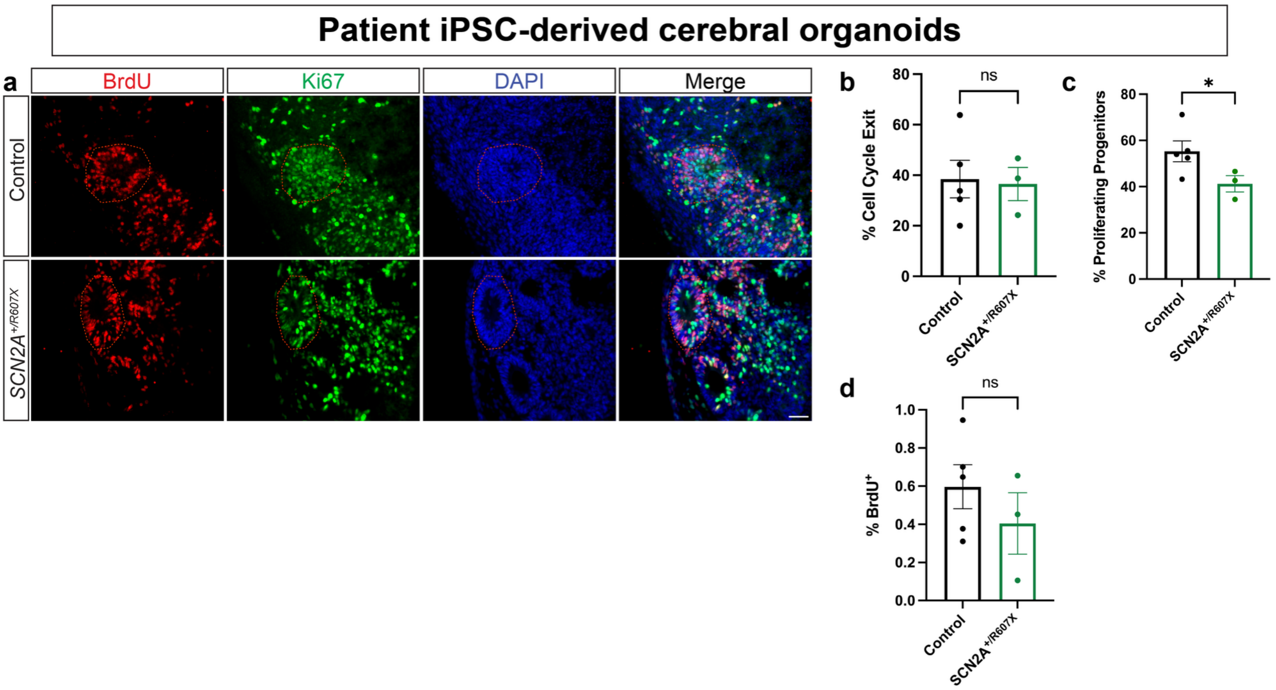


**Fig. S11.** ***SCN2A*** **does not regulate neural progenitor proliferation or cell cycle exit in patient iPSC-derived cerebral organoids**

**a)** Immunostaining for BrdU and Ki67 in Day 40 organoids. Scale bar = 50 µm.

**b-d)** Quantification of cell cycle exit, percentage of BrdU$^{+}$ cells and percentage of proliferating progenitors (N = 1 batch, Control n = 5 progenitor zones from 3 organoids, *SCN2A*$^{+/R607X}$ n = 3 progenitor zones from 3 organoids, mixed-effects models with Genotype as a fixed effect and each progenitor zone nested per genotype).

All data are reported as mean ± SEM. Mixed-effects models. *$p < 0.05$, **$p < 0.01$, ***$p < 0.001$, ****$p < 0.0001$; n.s., not significant.

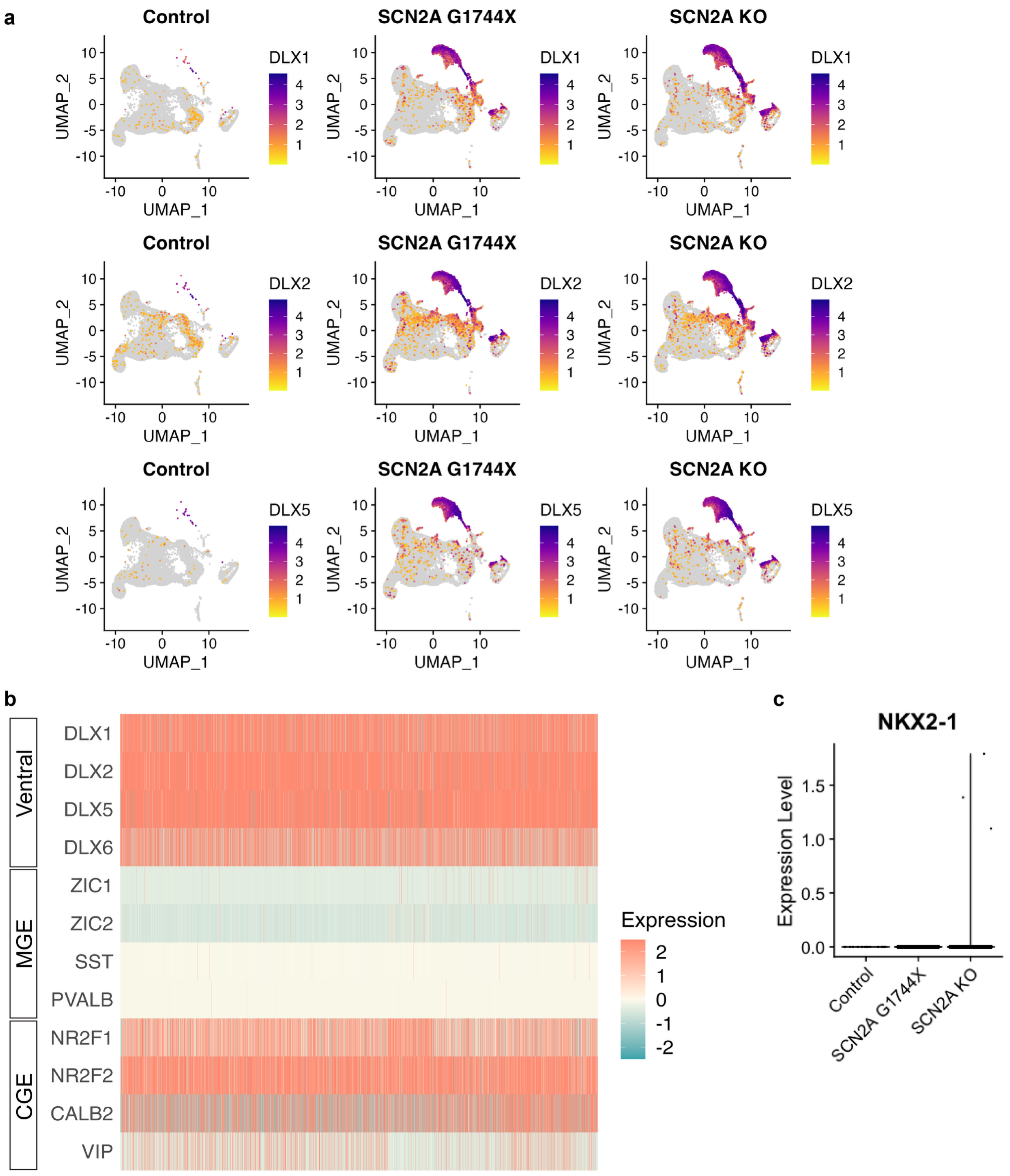


**Fig. S12. Informatic characterization of inhibitory neurons**
**a)** Feature plots showing expression of the ventral lineage DLX transcription factors separated by genotype.
**b)** Heatmap showing expression of markers for either ventral, MGE, or CGE lineages in InhibNs.
**c)** Violin plot showing minimal expression of the MGE lineage marker NKX2-1 in InhibNs.

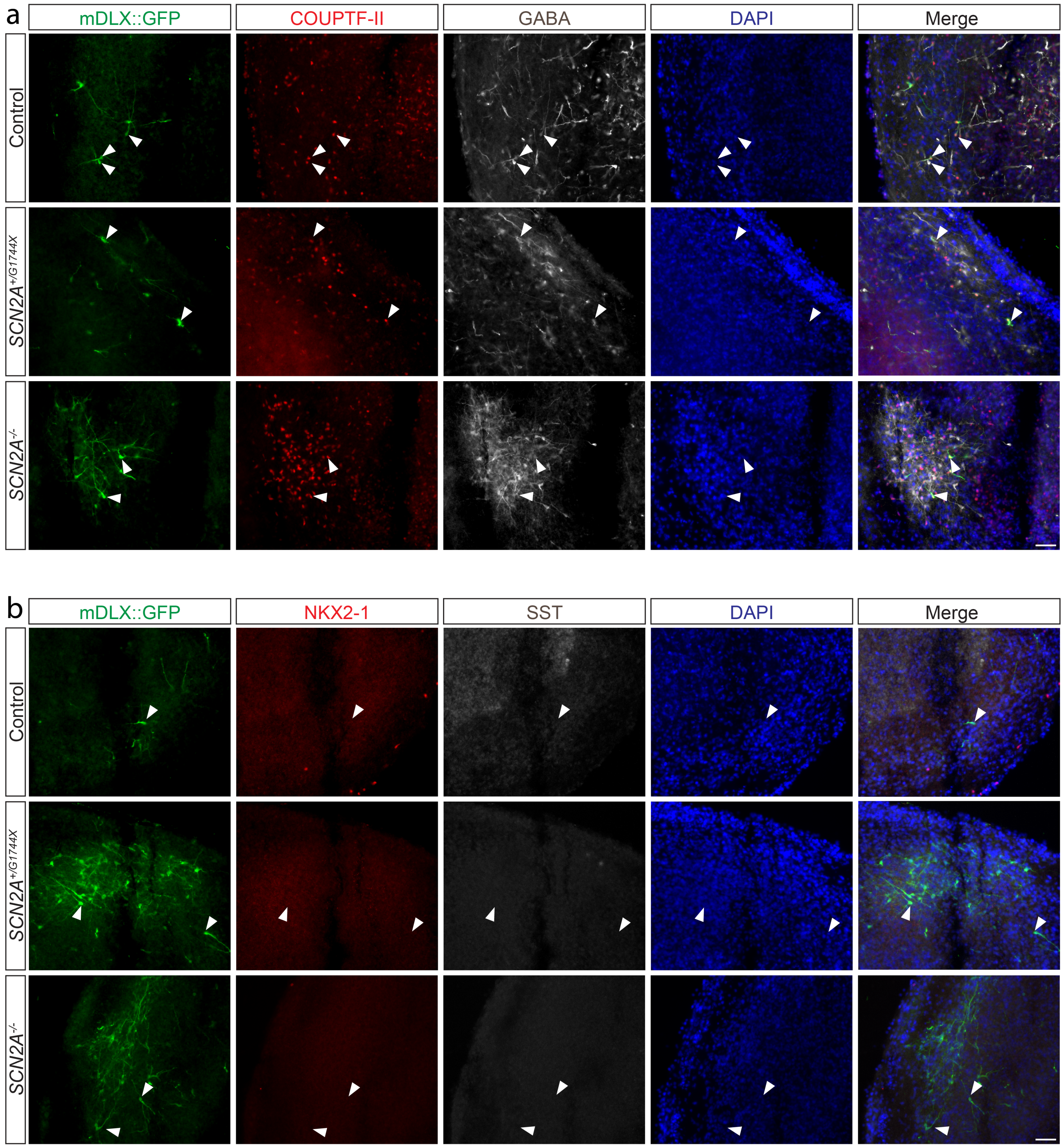


**Fig. S13. Cerebral organoid INs infected with AAV-mDLX-GFP are GABAergic.**
**a)** Immunostaining for AAV-mDLX-GFP-labelled inhibitory neurons, the neurotransmitter GABA, and the CGE marker COUPTF-II in H1 cerebral organoids. Scale bar = 50 µm.
**b)** Immunostaining for AAV-mDLX-GFP-labelled inhibitory neurons, somatostatin (SST), and the MGE marker NKX2-1 in H1 cerebral organoids. Scale bar = 50 µm.
Arrows denote cells positive or negative for the respective markers.

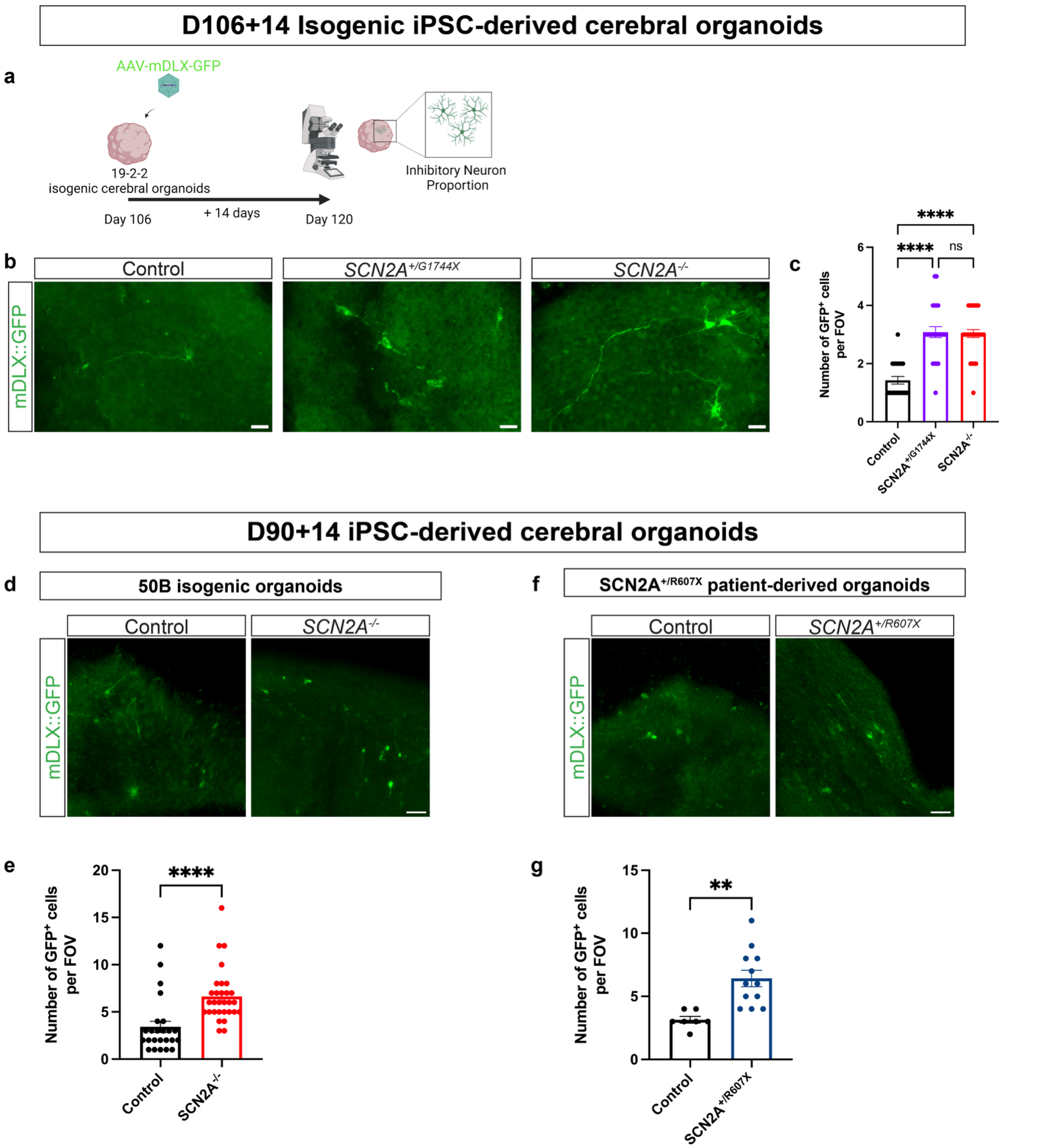


**Fig. S14. Loss of *SCN2A* leads to precocious production of CGE-like inhibitory neurons in iPSC-derived isogenic and patient-derived cerebral organoids**

**a)** Schematic of AAV viral labelling of inhibitory neurons.

**b)** Immunostaining for AAV-mDLX-GFP-labelled inhibitory neurons in 19-2-2 cerebral organoids. Scale bar = 20 μm.

**c)** Quantification of number of GFP$^{+}$ cells per FOV (N = 1 batch, Control n = 21 FOVs from 4 organoids, *SCN2A*$^{+/G1744X}$ n = 24 FOVs from 4 organoids, *SCN2A*$^{-/-}$ n = 30 FOVs from 4 organoids, mixed-effects models with Genotype as a fixed effect and each FOV nested per genotype).

**d)** Immunostaining for AAV-mDLX-GFP-labelled inhibitory neurons in 50B cerebral organoids. Scale bar = 50 μm.
**e)** Quantification of number of $GFP^+$ cells per FOV (N = 1 batch, Control n = 24 FOVs from 4 organoids, *SCN2A*$^{-/-}$ n = 30 FOVs from 4 organoids, mixed-effects models with Genotype as a fixed effect and each FOV nested per genotype).
**f)** Immunostaining for AAV-mDLX-GFP-labelled inhibitory neurons in *SCN2A*$^{+/R607X}$ patient-derived cerebral organoids. Scale bar = 50 μm.
**g)** Quantification of number of $GFP^+$ cells per FOV (N = 1 batch, Control n = 7 FOVs from 4 organoids, *SCN2A*$^{+/R607X}$ n = 12 FOVs from 4 organoids, mixed-effects models with Genotype as a fixed effect and each FOV nested per genotype).
All data are reported as mean ± SEM. For (c), mixed-effects models were used followed by Type III Wald $\chi^2$ test for fixed effects and post-hoc Tukey test. For (e and g), mixed-effects models were used. *$p < 0.05$, **$p < 0.01$, ***$p < 0.001$, ****$p < 0.0001$; n.s., not significant. Created in BioRender. Singh, K. (2026) https://BioRender.com/0otxclw.

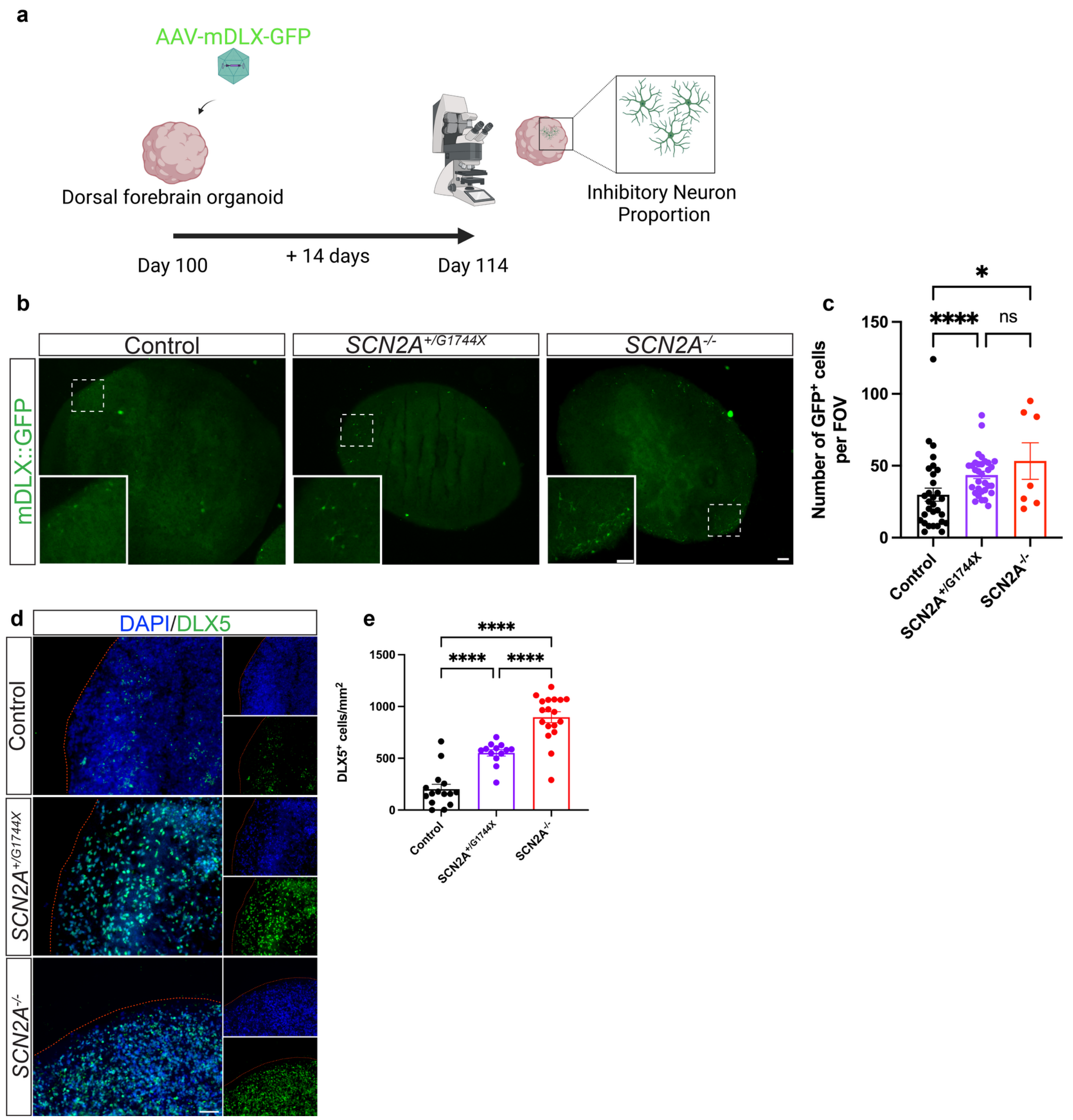


**Fig. S15. Loss of *SCN2A* leads to precocious production of CGE-like inhibitory neurons in H1 ESC-derived dorsal forebrain organoids**

**a)** Schematic of viral labelling of inhibitory neurons in dorsal (guided) forebrain organoids.

**b)** Immunostaining for AAV-mDLX-GFP-labelled inhibitory neurons in H1 cerebral organoids. Scale bar = 50 µm.

**c)** Quantification of number of $GFP^+$ cells per FOV (N = 1 batch, Control n = 30 FOVs from 4 organoids, $SCN2A^{+/G1744X}$ n = 32 FOVs from 4 organoids, $SCN2A^{-/-}$ n = 7 FOVs from 4 organoids, mixed-effects models with Genotype as a fixed effect and each FOV nested per genotype).

**d)** Immunostaining for the inhibitory neuron marker DLX5 in Day 100 organoids. Scale bar = 100 µm.

**e)** Quantification of density of $DLX5^+$ neurons in Day 100 organoids (N = 1 batch, Control n = 15 FOVs from 4 organoids, $SCN2A^{+/G1744X}$ n = 13 FOVs from 4 organoids, $SCN2A^{-/-}$ n = 18 FOVs from 4 organoids, mixed-effects models with Genotype as a fixed effect and each FOV nested per genotype).

All data are reported as mean ± SEM. Mixed-effects models with Type III Wald $\chi^2$ test for fixed effects, post-hoc Tukey test. *$p < 0.05$, **$p < 0.01$, ***$p < 0.001$, ****$p < 0.0001$; n.s., not significant. Created in BioRender. Singh, K. (2026) https://BioRender.com/0otxclw.

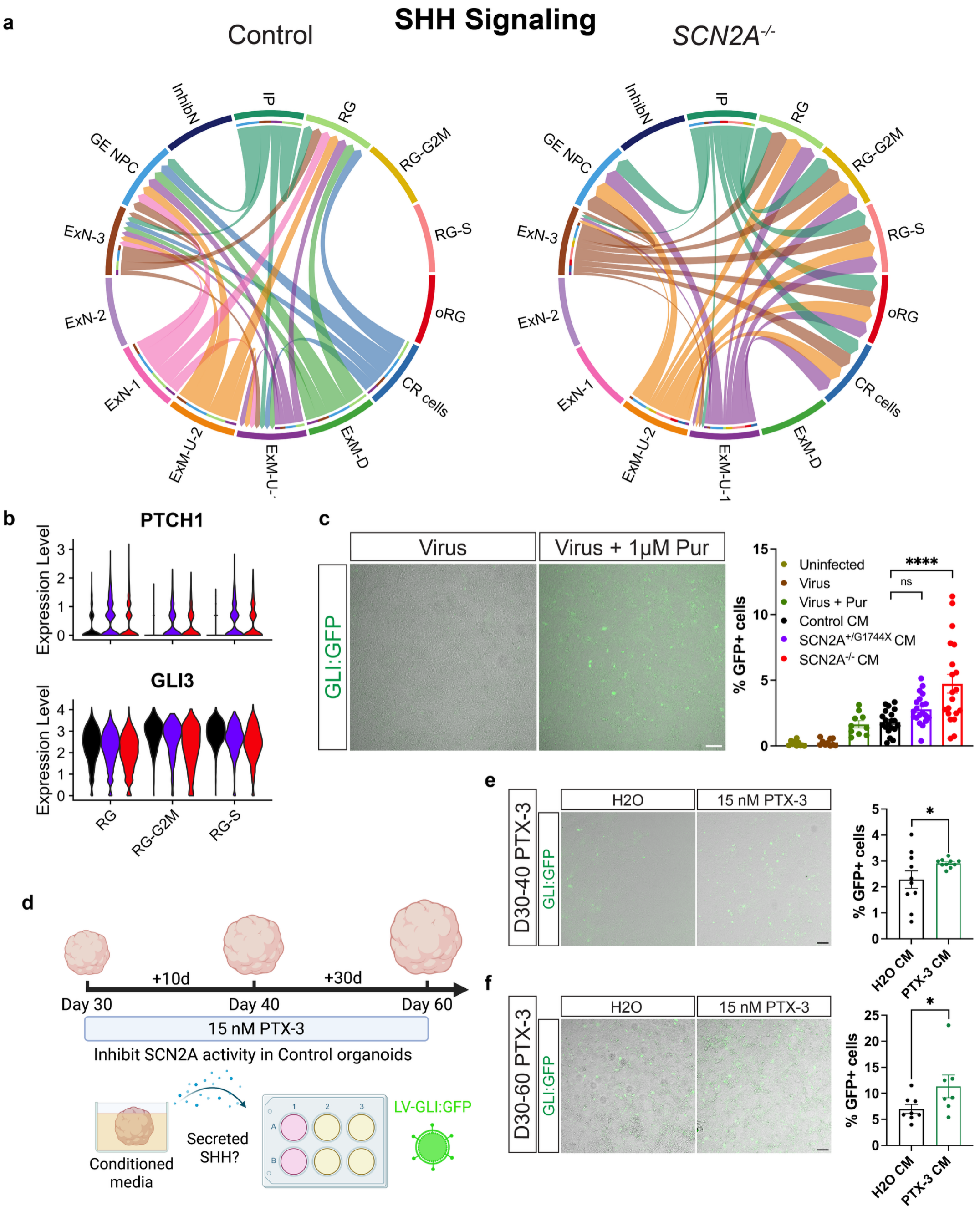


**Fig. S16. Loss of *SCN2A* or its inhibition results in elevated SHH signaling.**

**a)** Chord plots summarizing SHH signaling patterns from source to target cell types between Control and SCN2A KO organoids.

**b)** Violin plots of key differentially expressed SHH pathways genes in radial glia cell types.

**c)** Fluorescent images of infected HEK293T cells treated with and without purmorphamine and quantification of the percent GFP+ cells from wells treated with conditioned media from Day 60 SCN2A organoids (N = 1 batch, Uninfected, Virus, and Virus + Pur n = 5 FOVs, Control, *SCN2A*$^{+/G1744X}$ and *SCN2A*$^{-/-}$ CM n = 10 FOVs from 2 wells. Mixed-effects models with Genotype as a fixed effect and each FOV nested per genotype).

**d)** Schematic showing treatment of HEK293T cells transduced with LV-GLI:GFP and treated with conditioned media from Control and 15 nM PTX-3 treated organoids.

**e)** Fluorescent images of infected HEK293T cells treated with H2O or 10 day PTX-3 conditioned media and quantification of the percent GFP+ cells (N = 1 batch, H2O CM and PTX-3 CM n = 10 FOVs from 2 wells. Mixed-effects models with Condition as a fixed effect and each FOV nested per condition).

**f)** Fluorescent images of infected HEK293T cells treated with H2O or 30 day PTX-3 conditioned media and quantification of the percent GFP+ cells (N = 1 batch, H2O CM n = 8 FOVS from 2 wells and PTX-3 CM n = 7 FOVs from 2 wells. Mixed-effects models with Condition as a fixed effect and each FOV nested per condition).

All data are reported as mean ± SEM. Mixed-effects models with Type III Wald $\chi^2$ test for fixed effects, post-hoc Tukey test. *p < 0.05, **p < 0.01, ***p < 0.001, ****p < 0.0001; n.s., not significant. Created in BioRender. Singh, K. (2026) https://BioRender.com/0otxclw.

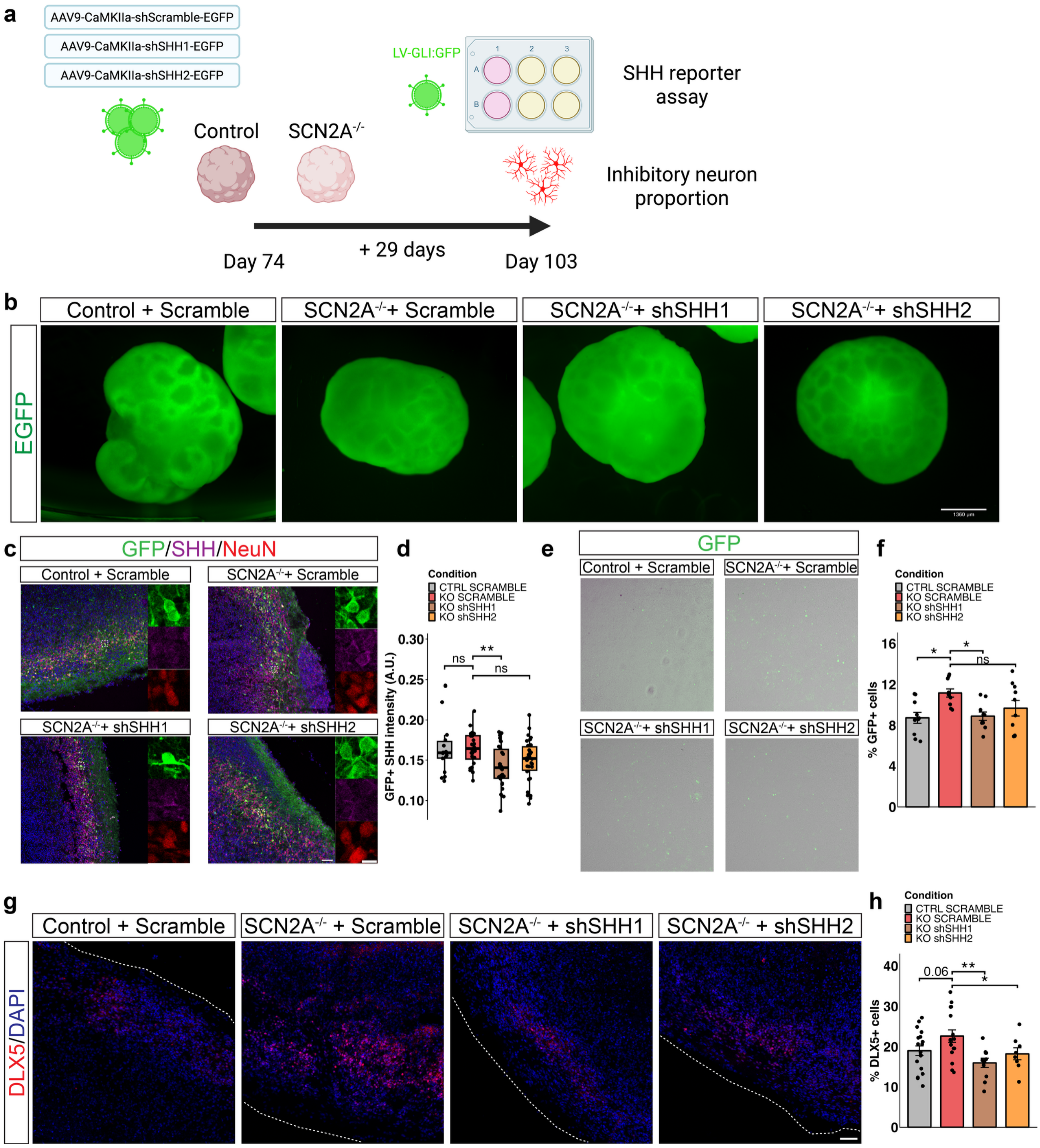


**Fig. S17. Elevated SHH signal and inhibitory neurogenesis is driven by *SCN2A*$^{-/-}$ neurons.**
**a)** Schematic showing SCN2A organoids transduced with shRNAs against SHH.
**b)** Fluorescent images of infected SCN2A organoids expressing EGFP and shRNAs after 21 days.
**c)** Immunostaining for GFP, NeuN and SHH in shRNA-infected organoids 29 days post-transduction. Scale bar = 100 µm.
**d)** Quantification of SHH intensity in GFP$^{+}$ cells (N = 1 batch, Control Scramble n = 18 FOVs from 4 organoids, *SCN2A*$^{-/-}$ Scramble n = 24 FOVs from 5 organoids, *SCN2A*$^{-/-}$ shSHH1 n = 30 FOVs from 6 organoids,

*SCN2A*$^{-/-}$ shSHH2 n = 30 FOVs from 6 organoids, mixed-effects models with Condition as a fixed effect and each FOV nested per organoid). Boxplots are median, quartiles and 1.5 IQR whiskers.
**e, f)** Fluorescent images of infected HEK293T cells treated with shRNA-infected organoid conditioned media and quantification of the percent GFP+ cells (N = 1 batch, Control Scramble n = 10 FOVs from 2 wells, *SCN2A*$^{-/-}$ Scramble n = 10 FOVs from 2 wells, *SCN2A*$^{-/-}$ shSHH1 n = 10 FOVs from 2 wells, *SCN2A*$^{-/-}$ shSHH2 n = 10 FOVs from 2 wells, mixed-effects models with Condition as a fixed effect and each FOV nested per well).
**g)** Immunostaining for the inhibitory neuron marker DLX5 in Day 90 organoids. Scale bar = 50 μm.
**h)** Quantification of % of DLX5+ neurons in shRNA-infected organoids (N = 1 batch, Control Scramble n = 18 FOVs from 4 organoids, *SCN2A*$^{-/-}$ Scramble n = 16 FOVs from 4 organoids, *SCN2A*$^{-/-}$ shSHH1 n = 11 FOVs from 4 organoids, *SCN2A*$^{-/-}$ shSHH2 n = 8 FOVs from 2 organoids, mixed-effects models with Condition as a fixed effect and each FOV nested per organoid).
All data are reported as mean ± SEM. Mixed-effects models with Type III Wald $\chi^2$ test for fixed effects, post-hoc Tukey test. *p < 0.05, **p < 0.01, ***p < 0.001, ****p < 0.0001; n.s., not significant. Created in BioRender. Singh, K. (2026) https://BioRender.com/0otxclw.

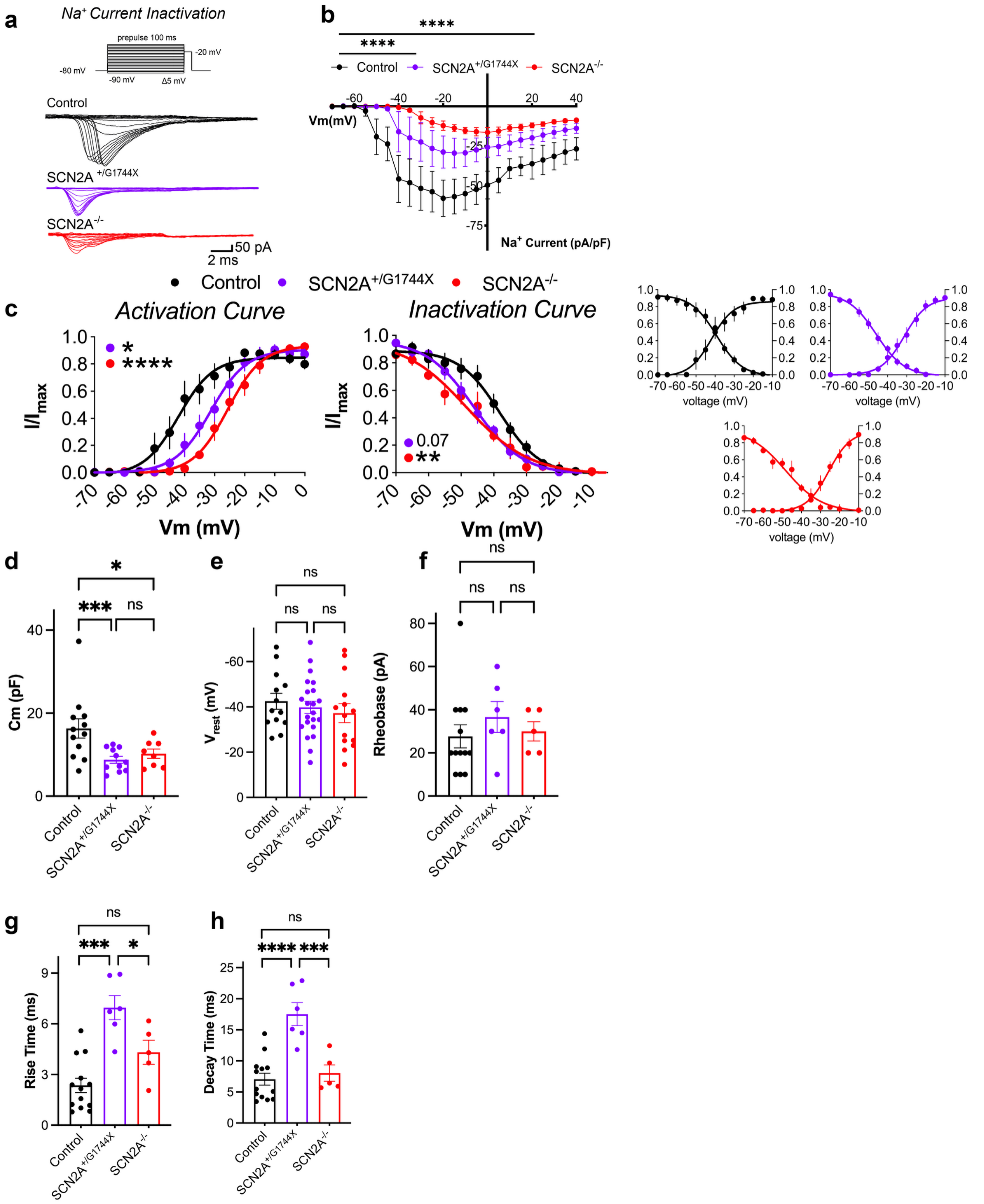


**Fig. S18. Additional electrophysiological characterization of *SCN2A* mutant organoids at Day 60.**

**a)** Representative sodium current inactivation traces.

**b)** Sodium current densities plotted against voltage (N = 2 independent batches, Control n = 12 neurons from 4 organoids, $SCN2A^{+/G1744X}$ n = 11 neurons from 4 organoids, $SCN2A^{-/-}$ n = 8 neurons from 3 organoids, mixed-effects models with Genotype and voltage as fixed effects and organoid as a random effect, Type III Wald $\chi^2$ test for Genotype and post-hoc Tukey test).

**c)** Sodium channel activation and inactivation curves in Control and SCN2A mutant neurons in Day 60 organoids (N = 2 independent batches, Control n = 12 neurons from 3 organoids, *SCN2A*$^{+/G1744X}$ n = 11 neurons from 4 organoids, *SCN2A*$^{-/-}$ n = 8 neurons from 3 organoids). Top graphs are overlaid data while bottom graphs are individual genotypes. Mixed-effects models with Genotype and voltage as fixed effects and organoid as a random effect, Type III Wald $\chi^2$ test for Genotype and post-hoc Tukey test.
**d)** Capacitance of H1 cerebral organoid neurons (N = 2 independent batches, Control n = 12 neurons from 4 organoids, *SCN2A*$^{+/G1744X}$ n = 11 neurons from 4 organoids, *SCN2A*$^{-/-}$ n = 8 neurons from 3 organoids).
**e)** Resting membrane potential of H1 cerebral organoid neurons (N = 2 independent batches, Control n = 13 neurons from 3 organoids, *SCN2A*$^{+/G1744X}$ n = 22 neurons from 4 organoids, *SCN2A*$^{-/-}$ n =14 neurons from 4 organoids).
**f)** Rheobase of H1 cerebral organoid neurons (N = 2 independent batches, Control n = 13 neurons from 3 organoids, *SCN2A*$^{+/G1744X}$ n = 6 neurons from 3 organoids, *SCN2A*$^{-/-}$ n = 5 neurons from 2 organoids).
**g, h)** Action potential-like depolarization rise (g) and decay (h) times recorded from H1 cerebral organoids neurons (N = 2 independent batches, Control n = 13 neurons from 3 organoids, *SCN2A*$^{+/G1744X}$ n = 6 neurons from 3 organoids, *SCN2A*$^{-/-}$ n = 5 neurons from 2 organoids).
All data are reported as mean ± SEM. Mixed-effects models using organoid as a random effect with Type III Wald $\chi^2$ test for fixed effects, post-hoc Tukey test. *p < 0.05, **p < 0.01, ***p < 0.001, ****p < 0.0001; n.s., not significant.

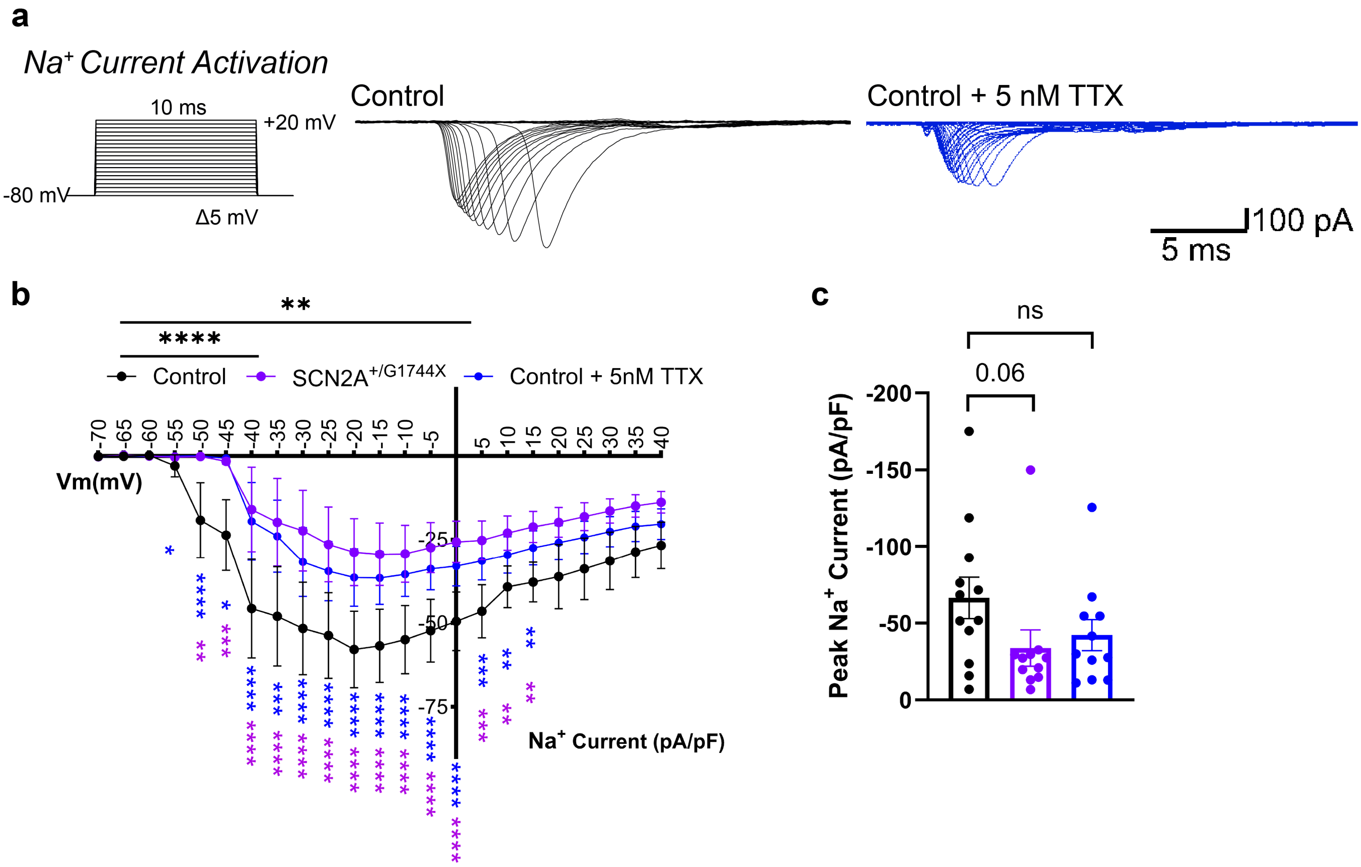


**Fig. S19. Electrophysiological characterization of 5nM TTX treatment in Control organoids at Day 60.**
**a)** Representative sodium current traces of Control and Control + 5 nM TTX cerebral organoids.
**b)** Sodium current densities normalized to cell capacitance (N = 2 independent batches, Control n = 12 neurons from 4 organoids, Control + 5 nM TTX n =11 neurons from 2 organoids, *SCN2A*$^{+/G1744X}$ data is from Fig 6c, mixed-effects models with Genotype and voltage as fixed effects including their interaction and organoid as a random effect). Black asterisks display significance across voltage steps while colored asterisks represent the respective condition significance at specific voltage steps.
**c)** Quantification of peak sodium current densities normalized to cell capacitance (N = 2 independent batches, Control n = 12 neurons from 4 organoids, Control + 5 nM TTX n =11 neurons from 2 organoids, *SCN2A*$^{+/G1744X}$ data is from Fig 6c, mixed-effects models with Genotype as fixed effects and organoid as a random effect).
All data are reported as mean ± SEM. Mixed-effects models with Type III Wald $\chi^2$ test for fixed effects, post-hoc Tukey test. *p < 0.05, **p < 0.01, ***p < 0.001, ****p < 0.0001; n.s., not significant.

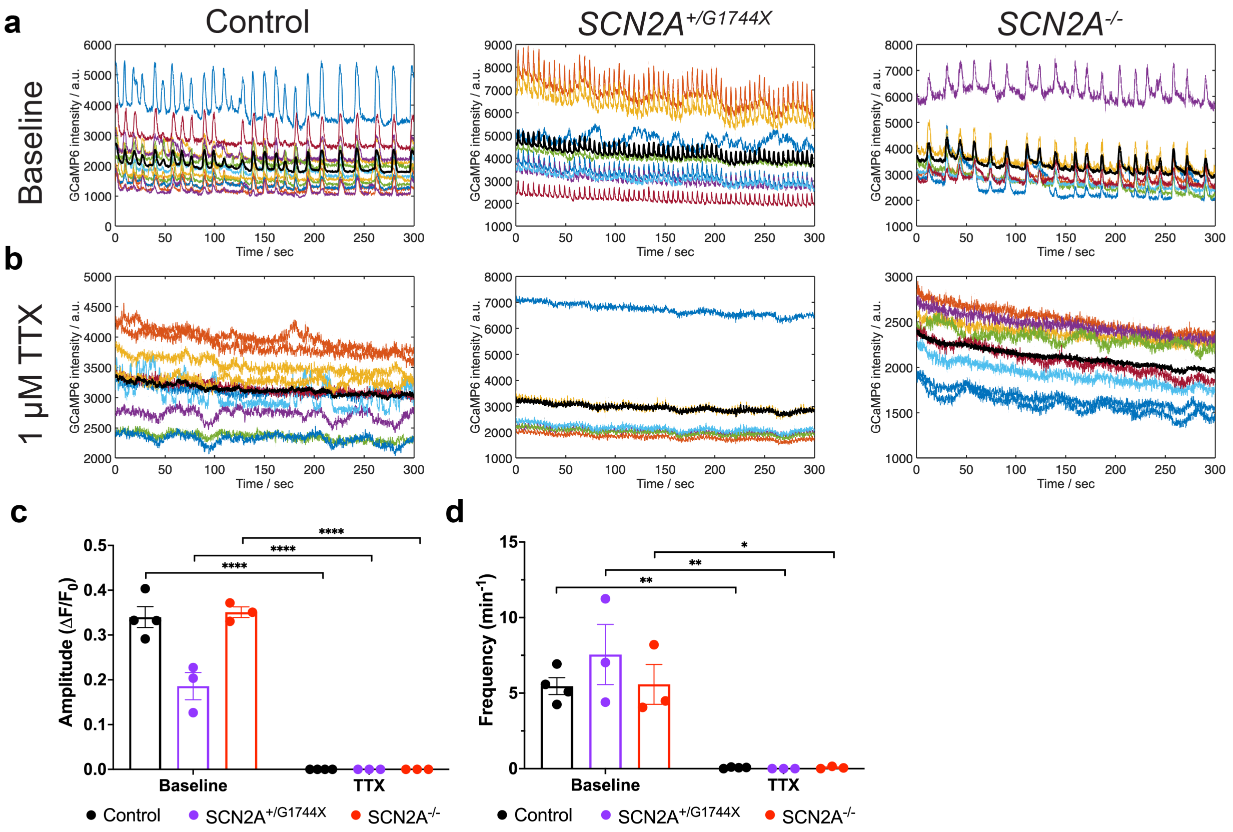


**Fig. S20. Spontaneous calcium transients obtained from GCaMP6-infected organoids are TTX-sensitive.**

**a)** Representative colored raw GCaMP6 intensity traces of Day 90 organoids at baseline with averaged intensities plotted in black.

**b)** Representative colored raw GCaMP6 intensity traces of Day 90 organoids treated with 1 μM TTX with averaged intensities plotted in black.

**c, d)** Quantification of amplitude (c) and frequency (d) of spontaneous calcium transients from Day 90 organoids at baseline and after treatment with TTX (N = 1 batch, Baseline: Control n = 4 FOVs from 2 organoids, *SCN2A$^{+/G1744X}$* n = 3 FOVs from 2 organoids, *SCN2A$^{-/-}$* n = 3 FOVs from 2 organoids; TTX: Control n = 4 FOVs from 2 organoids, *SCN2A$^{+/G1744X}$* n = 3 FOVs from 2 organoids, *SCN2A$^{-/-}$* n = 3 FOVs from 2 organoids, mixed-effects models with Genotype and treatment as fixed effects and each FOV nested within genotypes). All data are reported as mean ± SEM. Mixed-effects models with Type III Wald $\chi^2$ test for fixed effects, post-hoc Tukey test. *p < 0.05, **p < 0.01, ***p < 0.001, n.s., not significant.
